# Ciprofloxacin resistance in *Klebsiella pneumoniae*: phenotype prediction from genotype and global distribution of resistance determinants

**DOI:** 10.1101/2025.09.24.678318

**Authors:** Kara K. Tsang, Iana Amke, David M. Aanensen, Jabir Abdulahi, Alexander Aiken, Michael A. Bachman, Stephen Baker, Katherine Barry, Gherard Batisti Biffignandi, Emília Maria Medeiros Andrade Belitardo, Iván Bloise, Ilhem Boutiba, Chanté Brand, Sylvain Brisse, Susana Campino, Rafael Cantón, Alexandra Chiaverini, Daniela Maria Cirillo, Taane G. Clark, Teresa M. Coque, Jukka Corander, Marta Corbella, Alessandra Cornacchia, Jennifer Cornick, Annapaula Correia, Aline Cuénod, Vo Thi Trang Dai, Nicola D’Alterio, Sophia David, Federico Di Marco, Pilar Donado-Godoy, Jenny Draper, Adrian Egli, Refath Farzana, Nicholas A. Feasey, Edward J. Feil, Maria Laura Ferrando, Brian M. Forde, Aasmund Fostervold, Ebenezer Foster-Nyarko, Tommaso Giani, Claire L. Gorrie, Yukino Gütlin, Patrick N. A. Harris, Brekhna Hassan, Marisa Haenni, Wataru Hayashi, Eva Heinz, Marta Hernández-García, Marit Andrea Klokkhammer Hetland, Le Nguyen Minh Hoa, Nguyen Thi Hoa, Le Thi Hoi, Benjamin P. Howden, Odion O. Ikhimiukor, Jonathan R. Iredell, Dana Itani, Adam W. J. Jenney, Håkon Pedersen Kaspersen, Shizuo Kayama, Fahad Khokhar, Norikazu Kitamura, Appiah-Korang Labi, Margaret M.C. Lam, Val F. Lanza, Fernando Lázaro-Perona, Thongpan Leangapichart, Melese Hailu Legese, Le Thi Lien, Małgorzata Ligowska-Marzęta, Samuel Lipworth, Iren Høyland Löhr, S. Wesley Long, Giovanni Lorenzin, Alicia Fajardo Lubian, Amy J. Mathers, Andrew G. McArthur, Nubwa Medugu, Adane Mihret, Adriano Souza Santos Monteiro, Patrick Musicha, Geetha Nagaraj, Bernd Neumann, Mae Newton-Foot, Anderson O. Oaikhena, Iruka N. Okeke, João Perdigão, Luis G.C. Pacheco, Sally R. Partridge, David L. Paterson, Oliver Pearse, My H. Pham, Francesco Pomilio, Niclas Raffelsberger, Andriniaina Rakotondrasoa, KL Ravikumar, Joice Neves Reis, Sandra Reuter, Leah W. Roberts, Carla Rodrigues, Charlene Rodrigues, Jesus Rodriguez-Baño, Gian Maria Rossolini, Ørjan Samuelsen, Francesca Saluzzo, Kirsty Sands, Davide Sassera, Helena M.B. Seth-Smith, Andrea Spitaleri, Varun Shamanna, Norelle L. Sherry, Sonia Sia, Hanen Smaoui, Anton Spadar, Jörg Steinmann, Nicole Stoesser, Motoyuki Sugai, Yo Sugawara, Marianne Sunde, Arnfinn Sundsfjord, Göte Swedberg, Lamia Thabet, Nicholas R. Thomson, Harry A. Thorpe, M. Estée Torok, Van Dinh Trang, Nguyen Vu Trung, Paul Turner, Jay Vornhagen, Nguyen Hoang Vu, Boaz D. Wadugu, Timothy R. Walsh, Andrew Whitelaw, Hayley Wilson, Kelly L. Wyres, Corin Yeats, Koji Yahara, Meriam Zribi, Kathryn E. Holt, the KlebNET-GSP AMR Genotype-Phenotype Group

## Abstract

**BACKGROUND:** Ciprofloxacin resistant *Klebsiella pneumoniae* is common or emerging in many geographies, and knowledge of local resistance rates is important for empirical therapy. Whilst there are known *K. pneumoniae* ciprofloxacin resistance determinants, there is a lack of systematic data on the effect of determinants, alone and in combination, and there are no publicly accessible tools for predicting resistance from whole genome sequence data.

**METHODS:** The KlebNET-GSP AMR Genotype-Phenotype Group aggregated a matched genotype-phenotype dataset of n=12,167 *K. pneumoniae* species complex (*Kp*SC) isolates from 27 countries between 2001-2021. We developed a rules-based classifier to predict ciprofloxacin resistance by categorizing the number of quinolone resistance determining regions mutations in *gyrA* and *parC*, the number of plasmid-mediated quinolone resistance genes, and the presence/absence of *aac(6ʹ)-Ib-cr* (which can acetylate ciprofloxacin).

Predictive performance was assessed using the discovery dataset, for which we re-phenotyped discrepant isolates; and validated using externally contributed datasets (n=7,030 *Kp*SC isolates).

**RESULTS:** The rules-based classifier predicted R vs S/I with categorical agreement, sensitivity, and specificity >96%, and major/very major error rates <4%. Performance was similar across diverse *Kp*SC sources (human, animal, other), species, and intra-species lineages. External validation of the classifier yielded overall 93.12% categorical agreement [95% confidence interval (CI), 92.50-93.74%], 8.65% major errors [95% CI, 7.34-9.97%], and 6.20% very major errors [95% CI, 5.51-6.90%]. We implemented the classifier in Kleborate, a command-line tool that is integrated into the Pathogenwatch web platform. Using this to assess the global distribution of ciprofloxacin resistance determinants in *Kp*SC genomes available in Pathogenwatch (n=31,319, from 109 countries between years 2000-2023), we observed a significant positive association between national quinolone consumption rates and predicted ciprofloxacin resistance (R^2^=0.20, p=0.004).

CONCLUSIONS

Ciprofloxacin resistance phenotypes can be reasonably predicted from genotypes, which is sufficient for informing surveillance. However, unexplained resistance remains and accuracy is insufficient for clinical applications. We demonstrate the value of aggregating genotype-phenotype data to explore resistance mechanisms and develop predictors, but highlight complexities in combining phenotype data from different assays and standards.

## INTRODUCTION

*Klebsiella pneumoniae* cause a wide range of infections, particularly in healthcare settings. In hospitals, *K. pneumoniae* most commonly cause urinary tract, respiratory tract, and wound or soft tissue infections, which can progress to bloodstream infection if not effectively treated^1^.

In addition, hypervirulent strains of *K. pneumoniae* are a common cause of community-acquired pyogenic liver abscess, often in healthy individuals and can metastasize to distant sites, including most commonly the eye, lung and central nervous system^2,3^.

Antimicrobial resistance (AMR) is common in *K. pneumoniae*, and infections with carbapenem-resistant or third-generation cephalosporin (3GC)-resistant *K. pneumoniae* were associated with an estimated 630,000 deaths in 2021^4,5^. 3GC-resistant and carbapenem-resistant *Enterobacterales*, including *Klebsiella*, have been designated a critical priority for research and development by the World Health Organization (WHO)^6,7^.

Treatment of *K. pneumoniae* infection would ideally be informed by phenotypic antimicrobial susceptibility testing (AST) of the causative agent. However, a majority of infections (particularly in resource-limited settings that lack culture-based diagnostics and AST) are treated empirically, based on the clinical syndrome^8^ (e.g. sepsis, where WHO regimens vary by age group but consist of some combination of penicillins, aminoglycosides and 3GCs, either as dual therapy or monotherapy, though when available, drugs such as carbapenems are often utilised in settings where resistance to first and second line agents is widespread).

Ciprofloxacin is a relatively cheap and broad-spectrum orally-active antibacterial that appears in the WHO essential medicines list. It is classed as an ‘Access’ antibiotic in the WHO AWaRe book^8^, where it is recommended for empirical treatment of hospital-acquired urinary tract infection (UTI) and pyogenic liver abscess^8^. Ciprofloxacin can also be used to treat urinary, complicated intra-abdominal, and lower respiratory tract infections^9^. As such, ciprofloxacin use is significant globally, including in low-and middle-income countries where resistance to other accessible first-line drugs such as penicillins, 3GCs and gentamicin are increasing, more expensive antibiotics are not affordable, specific infections such as enteric fever are common, or intravenous delivery is demanding for healthcare systems.

Ciprofloxacin-resistant *K. pneumoniae* is common or emerging in many geographic areas^10^, and was associated with an estimated 430,000 deaths in 2021^4,5^. Since the 2000s, the ciprofloxacin resistance rate in *K. pneumoniae* has been well-known and exceeded 5% in North America, Europe, and Asia^11–14^. In the most recent WHO Global AMR Surveillance System (GLASS) report, the per-country ciprofloxacin resistance rates from over 50 countries had a median of over 40% in bloodstream infections (total infections 84,809) and urinary tract infections (total infections 287,972)^10^. Ciprofloxacin and other fluoroquinolones are bactericidal, as they inhibit bacterial DNA synthesis and replication by interfering with the binding of DNA by DNA gyrase (encoded by *gyrA* and *gyrB*) and topoisomerase IV (encoded by *parC* and *parE*)^15–18^.

Ciprofloxacin resistance is most often mediated by stepwise accumulation of mutations in the quinolone resistance determining regions (QRDR) of *gyrA* and *parC*, and occasionally of *gyrB* or *parE*^19^. Mutations in genes encoding porins (*ompK35* and *ompK36* on the chromosome)^20^ or efflux pumps (e.g., horizontally acquired *qepA* and *oqxAB* on the chromosome) can also influence the uptake and excretion of ciprofloxacin and other antibiotics in the bacterial cell^21–23^. Plasmid-mediated quinolone resistance (PMQR) mechanisms (sometimes referred to as transferable mechanisms of quinolone resistance-TMQR^24^) inhibit quinolone binding via target site modifications^25^ through the acquisition of mobile quinolone resistance genes. The *qnr* genes (such as *qnrA*, *qnrB*, and *qnrS*), which encode proteins that protect the quinolone targets to confer PMQR, were first reported in *K. pneumoniae* in 1998^16,26,27^. An efflux pump discovered in 2007, *qepA*, also confers PMQR^23^. One acquired aminoglycoside acetyltransferase gene variant, *aac(6ʹ)-Ib-cr,* is also able to acetylate ciprofloxacin and is reported to mediate resistance to fluoroquinolones in some organisms^21,28,29^. In *K. pneumoniae*, it is not as clear whether *aac(6ʹ)-Ib-cr* confers decreased susceptibility to fluoroquinolones since previous experimental efforts have not considered the joint contribution of QRDR mutations in this context^30,31^.

Bacterial whole-genome sequencing (WGS) is increasingly used to support AMR surveillance and infection control, and genome data can provide insights into not only rates of resistance but also transmission dynamics and the emergence and spread of resistance within bacterial populations^32–34^. It is therefore desirable to understand the combinatorial effects of the known genetic determinants of resistance to predict resistance phenotypes from WGS data. Sequence-based predictions are also potentially useful in settings where phenotypic testing is not possible, such as sequence-based diagnostics (based on metagenomics, amplicon-sequencing or micro-arrays) where no isolate is available for AST^35–37^, or to support the development of empiric treatment guidelines where genomic surveys have been undertaken. Sequence-based predictions rely on using bacterial genome features (i.e., annotated genes, mutations, *k*-mers) and their corresponding antibiogram data to generate an antibiotic susceptibility prediction. A rules-based method makes predictions using expert-curated knowledge with supportive data to generate rules based on the presence/absence of known resistance determinants. There are more complicated prediction methods, such as logistic regression and other machine learning algorithms, where it can be difficult to understand which genetic determinants drive a particular phenotype and how they interact in combination^38^.

Whilst a range of ciprofloxacin resistance determinants are known in *K. pneumoniae*, there is a lack of systematic data on the effect of specific determinants, alone and in combination, and there are no publicly accessible tools for predicting resistance from *K. pneumoniae* WGS data. Several studies reported genotypes and phenotypes of local cases of ciprofloxacin resistant *K. pneumoniae*^39–43^, but these lack the sample size and diversity needed to tease apart the effects of specific determinants and combinations or to develop convincing predictors.

Another factor is the difficulty of interpreting *Enterobacterales* ciprofloxacin susceptibility particularly when the AST measurement is in the area of technical uncertainty^44^. Two studies have explored the use of machine learning to predict minimum inhibitory concentration (MIC) of ciprofloxacin and other drugs from *K. pneumoniae* WGS data. The first used n=1,664 isolates from a single facility in Texas, USA, 86% of which had ciprofloxacin MIC >2 mg/L and most of which belonged to two healthcare-associated carbapenem-or 3GC-resistant clones (ST258, ST307) to model ciprofloxacin resistance^45^. The second study supplemented this data with data from additional isolates from diverse sources in Pavia, Italy^46^ to reach a total of n=3,158 isolates with available ciprofloxacin MICs^47^. These studies support that ciprofloxacin resistance phenotypes can in principle be predicted from WGS data, and the second study estimated heritability (*h*^2^) for ciprofloxacin MIC above 80%.

However, these models lack interpretability, or external validation on large diverse datasets, and were not implemented as predictors that others can apply to interpret their own data.

Here, we investigate ciprofloxacin resistance genotype-phenotype relationships in a diverse collection of n=12,167 sequenced isolates of the *K. pneumoniae* species complex (*Kp*SC, which comprises *K. pneumoniae* and 4 closely related species), with matched ciprofloxacin phenotype data, contributed by 26 study groups covering 27 countries as part of the KlebNET-Genomic Surveillance Platform (GSP) AMR Genotype-Phenotype Group. We develop a rules-based classifier to predict ciprofloxacin resistance from WGS data, implemented in the widely-used Kleborate genotyping tool, and demonstrate external validation on a further 7,030 isolates from 19 study groups across six continents. Finally, we evaluated the global distribution of quinolone resistance mechanisms in 31,319 *Kp*SC isolates from 109 countries and the association of these with quinolone consumption rates.

## METHODS

### *K. pneumoniae* species complex isolate collection

In the discovery phase, we used 12,167 *Kp*SC genomes with matched antibiotic susceptibility testing (AST) data (summary in **Table S1**). The collection includes 10,634 *K. pneumoniae*, 1,037 *Klebsiella variicola* subsp. *variicola* (*Kvv*), 286 *Klebsiella quasipneumoniae* subsp. *similipneumoniae* (*Kqs*), 189 *Klebsiella quasipneumoniae* subsp. *quasipneumoniae* (*Kqq*), 9 *Klebsiella quasivariicola*, 11 *Klebsiella variicola* subsp. *tropica,* and 1 *Klebsiella africana*.

These diverse isolates were collected from 26 study groups across 27 countries (spanning all continents except Antarctica), have a diverse range of isolation sources (86% human, 9% animal, 3% environment, 0.7% marine, and 0.7% food/feed) and temporal span (collected from 2001-2021) (**Figure S1**).

### Genome sequence analysis

Illumina reads were assembled into genomes using Unicycler^48^ (v0.4.8) (accessions and assembly metrics in **Table S2**). All assemblies met the pre-agreed KlebNET-GSP criteria of <5% contamination (assessed using KmerFinder^49^ v3.2); Kleborate-designated species match of “strong”; ≤500 contigs; genome size 4,969,898–6,132,846 bp; and G+C content in the range 56.35%–57.98%, in addition to other *Kp*SC species-specific criteria as described https://bigsdb.pasteur.fr/klebsiella/genomes-quality-criteria/. Contig metrics across the dataset were: contig count, mean 132.9, minimum (min) 2, maximum (max) 499, standard deviation (sd) 70.7; N50, mean 335,221 bp, min 19,327 bp, max 5,474,158 bp, sd 436,968 bp; genome size, mean 5,516,257 bp, min 4,991,333 bp, max 6,132,239 bp, sd 185,779 bp; G+C content, mean 57.23%, sd 0.25%.

Kleborate^50^ (v3.0.0) was used to identify the presence of QRDR substitutions (e.g., in GyrA and ParC, which we refer to as QRDR mutations hereafter since the substitutions are caused by mutations in the QRDRs of *gyrA* and *parC*); PMQR genes (e.g., *qnr* and *qep* genes); *aac(6ʹ)-Ib-cr* (reported in the AGly column); porin defects associated with AMR (i.e., loss of OmpK35 or OmpK36, and insertions in loop 3 of OmpK36). Kleborate was also used to call 7-locus multi-locus sequence types (STs)^51^. Kleborate genotype results are included in **Table S2**. To designate imprecise matches to acquired resistance gene alleles, Kleborate appends ‘*’ (to indicate ≥90% and <100% identity at amino acid level across the full length of the reference allele) or ‘?’ (to indicate ≥80% and <100% coverage of the reference allele). These symbols are retained in the individual-variable analysis presented here, and absence of these symbols indicates a perfect match at the protein level. As imprecise matches were rare (n=83/18,884, 0.4%), they were excluded from regression modelling. Note the Kleborate database includes two reference alleles for some resistance genes, these are labelled with the suffix ‘.v1’ and ‘.v2’ in all outputs. Two reference alleles are included for *aac(6ʹ)-Ib-cr*, labelled v1 (accession MW579371.1) and v2 (NG_052213.1); both harbour the W102R and D179Y mutations required for acetylation of fluoroquinolones^29^. The distribution of hits for these individual alleles, and their imprecise matches, are shown separately in the single-variable analyses; as all showed the same association with resistance, all *aac(6ʹ)-Ib-cr* hits were combined into a single variable for regression modelling and rules-based classifier testing.

To investigate ciprofloxacin resistant isolates that had no known quinolone resistance genetic determinants, we also screened for variants of potentially relevant genes using BLASTN to compare each allele to a wildtype reference sequence: *ompK35* (WP_004141771.1, NZ_CP073076.1:2568274-2569353)*, ompK36* (WP_002913005.1, NZ_CP046939.1:1613497-1614600)*, gyrA* (WP_004184961.1, NZ_CP073076.1:812319-812678)*, parC* (WP_004900648.1, NZ_CP073076.1:5234743-5235102), and the *oqxAB* operon (including *oqxA* (WP_002914189.1, NZ_JAALAM010000003.1:40277-41452), *oqxB* (WP_117254594.1, NZ_UKBZ01000005.1:386801-389953), *oqxR* (WP_000888203.1, NZ_JAKRRC010000012.1:123786-124265)*, rob* (WP_085783298.1, NZ_CP083032.1:1141725-1142090)*, norB* (WP_004148137.1, NZ_CP132670.1:2862207-2863595)*, rarA* (WP_002914178.1, NZ_JASEKY020000003.1:6592-7476). We estimated the copy numbers of the intrinsic, chromosomal genes, *oqxA* and *oqxB,* by analysing Illumina read sets, calculating the ratio of read depth for each gene vs the mean read depth of the seven *K. pneumoniae* MLST loci, using SRST2^52^ (v0.2.0) to perform the mapping and depth calculations. Lastly, we used NCBI’s AMRFinderPlus^53^ (v3.12.8, specifying organism option to *Klebsiella pneumoniae*) to search for known resistance-associated mutations in *oqxR*.

### Susceptibility phenotyping data and interpretation

Ciprofloxacin susceptibility phenotypes were determined by the contributing laboratories using a range of phenotypic methods including disk diffusion (n=3073), agar dilution (n=498), gradient diffusion (n=4), semi-automated AST methods (Vitek2 (n=4,640), Phoenix (n=3,620)), and broth microdilution via Sensititre (n=332), that were performed according to CLSI^54^ or EUCAST^55^ guidelines (see isolate-level data in **Table S3)**. Currently, both EUCAST and CLSI define isolates with MIC ≤0.25 mg/L as susceptible (S) and >0.5 mg/L as resistant (R). EUCAST defines the epidemiological cut-off (ECOFF) value for the wildtype (WT) MIC distribution as ≤0.125 mg/L, with 95% confidence interval 0.06-0.25 mg/L. Using the ECOFF point estimate of MIC ≤0.125 mg/L would result in just 3% of our data being classified as WT. However, using the upper bound of MIC ≤0.25 mg/L, which is the same as the susceptibility breakpoint, better fits the observed bimodal distribution of our own MIC data (**Figure S2**) and results in 44% of isolates being classified as WT. Therefore, we define non-wild type (NWT) as >0.25 mg/L. For disk diffusion, EUCAST and CLSI use disk content of 5 µg (has remained unchanged since 2009 for EUCAST^56^ and 1985 for CLSI^57^) and agree the zone diameter breakpoint of <22 mm for R (which is also equal to the ECOFF) but differ slightly on the definition of S (≥26 mm for CLSI, ≥25 mm for EUCAST). We interpreted each dataset according to the methods with which the data were generated.

However in practice, only n=3 isolates assessed using CLSI had a disk diffusion zone diameter of 25 mm, thus only n=3/3,073 (0.098%) isolates would have been classified differently if we were to apply the EUCAST cut-off to all disk diffusion data.

EUCAST guidance notes that for surveillance, one should never group intermediate / susceptible at increased exposure (I) and R, only S/I. In our rules-based classifier, the prediction outcomes are WT S, NWT I, and NWT R. Overall, we define the primary outcome variable for our rules-based and logistic regression classifiers as R / NWT R (defined by MIC or disk diffusion, as described above), and present as a secondary outcome data on prediction of NWT (equivalent to I/R or NWT I/NWT R).

To improve the quality of the phenotypic data before analysis, we sought to re-test isolates that had at least one QRDR mutation but were recorded as phenotypically S (n=88), and isolates with no known quinolone resistance determinants but were recorded as phenotypically R (n=252). We were unable to re-test all isolates as there were resource and storage constraints. Of those with QRDR mutations but originally recorded S, n=28 were re-tested and n=14 (50%) were confirmed R. Of those with no known determinants but recorded R, n=122 were re-tested and n=104 (85.2%) were confirmed to be S. All analyses presented use the latest, re-tested data, and isolates that could not be re-tested retained their original classification.

### External validation of a rules-based ciprofloxacin resistance classifier

We implemented the rules-based classifier in Kleborate (v3.0.0; cipro_prediction branch), to facilitate independent external validation on datasets that were not used to develop the classifier. A total of 7,030 genomes contributed from 19 studies were considered in the external validation set. All genomes were analysed using the same Kleborate code (v3.0.0; cipro_prediction branch) to generate genotyping and ciprofloxacin resistance prediction results, then compared to matched AST data using R code (available at <u>github.com/klebgenomics/cipropaper/</u>), which includes filtering out assemblies not meeting the KlebNET-GSP genome assembly quality control criteria (as previously described). To support inclusion of unpublished datasets without requiring sharing of raw data, we gave dataset owners the option to either run the validation analysis themselves using the code we provided (EUSCAPE/EURECA and IRCCS San Raffaele Scientific Institute datasets), or to share with us the genetic data (reads, assemblies, or Kleborate output) and matched antibiogram data for validation by the study analysis team (all other datasets).

A summary of the included datasets, including assembly method and AST methods, are given in **Table S4**. Strain-level genotypes and phenotypes for these datasets are provided in **Tables S5 and S6**. Genotype profiles and predictions were cross-tabulated against phenotype data (S/I/R calls re-interpreted using the EUCAST clinical breakpoints noted above), to calculate evaluation metrics and plot genetic vs. phenotypic data. Phenotype re-testing on the internal validation datasets showed that isolates with borderline phenotype measures were frequently misclassified, but we were unable to conduct re-testing on external validation samples. To reduce the impact of phenotype assay variability we calculated performance metrics excluding isolates phenotyped as ‘I’ (i.e. MIC=0.5 mg/L, thus within one doubling distribution of either ‘S’ or ‘R’ categories; or disk zone 22-25 mm). Pooled metrics were calculated by including all isolates with phenotype calls of S or R. Per-dataset metrics were also calculated for datasets with at least 25 high quality genomes with calls of S or R, including more than 10 of each category (S and R).

### Public genome and quinolone consumption data

Kleborate v3.2.4 was implemented in Pathogenwatch (v23.4.4)^58^, and calls made from publicly available *Kp*SC genomes were downloaded on August 11, 2025 from the public Pathogenwatch collection (which includes assemblies of all KpSC read sets available in NCBI as of March 29, 2023). In total, 51,592 of these genomes passed the KlebNET-GSP genome quality control criteria described above; these had variable metadata, and we included only those with year (from 2000 onwards), country, BioProject accession, and Life Identification Number (LIN) code (n=42,643). To mitigate sampling bias and reduce over-representation of clonal isolates (i.e., outbreak investigation), one random isolate was selected from each group of isolates with the same LIN code and ciprofloxacin rules-based classifier genotype profile that were collected from the same BioProject, country, and year. As a result, n=31,319 unique genomes, which includes representation from 109 countries and years 2000-2023, were used for downstream analysis. The frequency of ciprofloxacin resistance genotypes and predictions per country, year, and ST were calculated using R v4.2.3. Estimates of national annual quinolone consumption data, for the years 2000-2018, were downloaded from the Global Research on Antimicrobial Resistance (GRAM) project^59^.

## Statistical analysis

Associations between genetic determinants and ciprofloxacin phenotypes were assessed individually in single-variable analyses, using Wilcoxon rank sum test for association with log_2_(MIC) or disk diffusion zone diameter (two-sample tests, implemented in base R function *wilcox.test*) and Fisher’s exact test to calculate odds ratio (OR) and p-value for association with categorical phenotypes (R vs S/I, NWT vs WT, implemented in base R function *fisher.test*).

Associations between genetic determinants and ciprofloxacin phenotypes were also assessed in multivariable analyses, using Firth’s logistic regression (implemented in the R package logistf v1.26.0). Model fit was investigated using Akaike Information Criterion (AIC, using the logistf package function *extractAIC*), and penalized likelihood ratio test to compare model pairs (with vs without interaction terms, using the logistf package function *anova.logistf*).

Predictive performance of regression models was assessed by comparing the predictions generated by applying the fitted model to the same data (using the logistf package function *predict.logistf*) with the observed phenotypes. Metrics reported are: categorical agreement, the proportion of genomes for which the predicted category (R vs S/I, or NWT vs WT) matched the observed phenotype category; sensitivity, the proportion of R genomes predicted as R; specificity, the proportion of S/I genomes predicted as S/I; major errors (ME), the proportion of S/I isolates genomes that were classified as R (i.e., false-resistant rate); and very major errors (VME), the proportion of R genomes that were classified as S/I (i.e., false-susceptible rate). Target values for an antimicrobial susceptibility diagnostic test are: categorical agreement >90%, ME rate <3%, VME rate <1.5%^60^.

The same metrics were used to assess the overall performance of the rules-based classifier, by comparing the predicted class against the observed phenotype category. Performance metrics were evaluated for the full dataset (n=12,167 genomes), and separately for subsets of data stratified by species or ST.

The positive predictive value (PPV) for individual genetic variants was calculated as the proportion of variant-positive isolates that were phenotypically resistant; 95% confidence interval for PPV was calculated as 95% confidence interval of this proportion, i.e. p ± 1.96*sqrt(p*(1-p)/N). In addition, the predictive value of each component genotype profile used by the rules-based classifier was assessed by calculating PPV, the proportion of genomes with the genotype profile that were phenotypically R.

We used linear regression to model the relationship between national annual quinolone consumption (measured as Defined Daily Doses (DDD) per 1000 inhabitants per day), and national annual ciprofloxacin resistance prevalence (estimated by Kleborate v3.2.4) and mean numbers of ciprofloxacin resistance determinants, using the *lm* function in R v4.2.3, and analysis of variance (ANOVA) for model comparison using the *anova* function.

All statistical analyses were performed in R v4.4.1. Code is available at <u>github.com/klebgenomics/cipropaper</u> (DOI:10.5281/zenodo.17193873).

## RESULTS

### Ciprofloxacin genotype-phenotype relationships

Our discovery dataset comprised 12,167 *K. pneumoniae* species complex isolates with matched whole genome sequence and ciprofloxacin susceptibility data, in the form of either disk diffusion (n=3,073, 25%) or MIC measures (n=9,094, 91%; of which n=8,260 were assessed using semi-automated platforms). The distributions for MIC and zone diameter values are shown in **Figures S2-3**, together with the clinical breakpoints used for interpretation. In total, n=5,388 (44%) isolates were classified as wildtype (WT; all ciprofloxacin susceptible (S)) and n=6,779 (56%) as non-wildtype (NWT; of which n=602 (5%) were intermediate / susceptible at increased exposure (I) and n=6,177 (51%) were resistant (R)).

We identified QRDR mutations in n=4,742 genomes (39.0%), PMQR genes (*qnr* allele only (n=4,019)*, qep* allele only (n=7), *qnr* and *qep* alleles (n=11)) in n=4,037 (33.2%) genomes, and *aac(6ʹ)-Ib-cr* in n=2,924 (24.0%) genomes; n=5,671 (46.6%) of genomes lacked any of these determinants. The frequencies of each determinant and their association with R and NWT phenotypes are shown in **Table S7**. All resistance determinants were rare amongst susceptible isolates (QRDR 0.8%, PMQR 1.8%, *aac(6ʹ)-Ib-cr* 2.5%), and were significantly more prevalent amongst resistant isolates (QRDR 75.7%, PMQR 60.8%, *aac(6ʹ)-Ib-cr* 44.3%; see **Table S7**). However, given that many genomes carried multiple resistance determinants, we were interested in teasing apart the contributions of specific determinants alone and in combination.

Eight individual QRDR mutations across 112 isolates were observed ‘solo’, i.e., in the absence of other QRDR mutations, PMQR genes, or *aac(6ʹ)-Ib-cr*. These include four mutations at GyrA-83 (83F, 83Y, 83L, 83I; each were independently associated with resistance, see **Figure 1a, Table S7**) and four mutations at GyrA-87 (87N, 87G, 87Y, 87H; the latter was observed in a single susceptible genome, but the other three mutations were independently associated with resistance, see **Figure 1a, Table S7**). ParC-80 and ParC-84 mutations were not observed solo, all detected instances were in genomes with a GyrA-83 mutation. Amongst isolates with a GyrA-83 mutation, GyrA-87 mutation, and no PMQR genes or *aac(6ʹ)-Ib-cr*, the presence of a ParC-80 or ParC-84 mutation was associated with a significant increase in MIC (median 2 mg/L vs 1 mg/L, p=0.002 using Wilcoxon rank sum test) and reduction in disk diffusion zone diameter (median 6 mm vs 20 mm, p=5×10^-9^) (see **Figures S4-5**).

**Figure 1.**
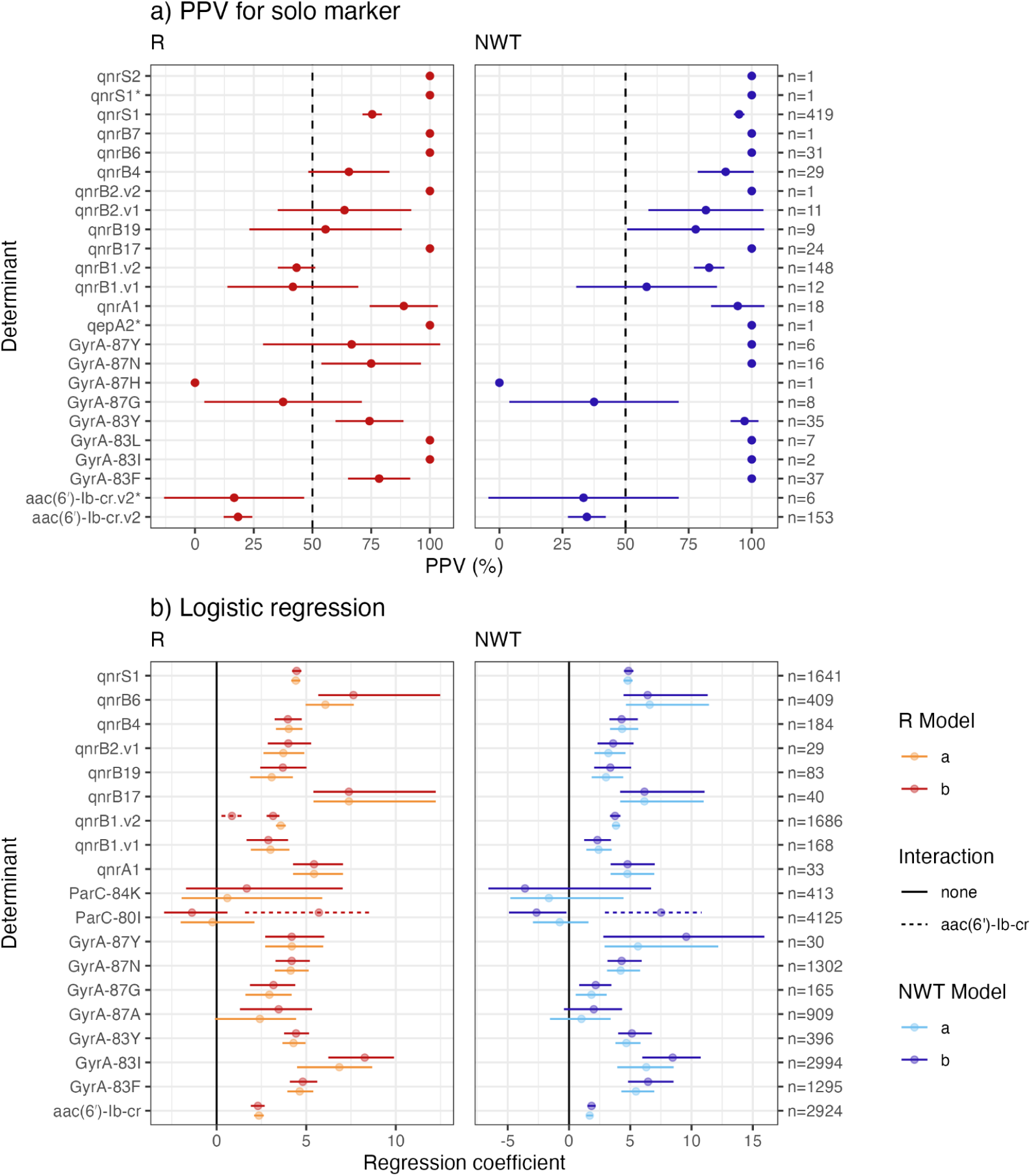
Measures of association with ciprofloxacin resistance for individual genetic determinants. **(a)** Positive predictive value (PPV) for each genetic determinant ‘solo’, i.e. amongst genomes carrying no other QRDR mutation, PMQR gene or *aac(6ʹ)-Ib-cr*, for the category R (resistant) or NWT (non-wildtype). Plotted lines indicate 95% confidence interval. The total number of genomes in which each determinant was found solo, amongst which the PPV is calculated, is indicated on the right. **(b)** Statistically significant coefficients of association with response variables R or NWT, estimated using logistic regression, for all determinants identified in ≥20 genomes (p<0.05). For each response variable (R, NWT), two models were fit: a, simple additive model including all determinants identified in ≥20 genomes; b, additive model with an interaction term for each determinant with *aac(6ʹ)-Ib-cr.* Coefficients for interaction terms are only plotted where there were significant (p>0.05) positive associations with the response variable.

Amongst isolates lacking any PMQR genes or *aac(6ʹ)-Ib-cr* (n=7,457), there was a significant association between the presence of any QRDR mutation and ciprofloxacin MIC (median 2 mg/L vs 0.25 mg/L, p<2×10^-16^), disk diffusion zone diameter (median 9 mm vs 29 mm, p<2×10^-16^), NWT (OR 827 [95% CI 525-1,264], p<2×10^-16^), and clinical resistance (OR 1522 [1,050-2,420], p<2×10^-16^). The number of QRDR mutations was also significantly associated with MIC (median 0.25 mg/L with no mutations, median 1 mg/L with one mutation, median 2 mg/L with two or more mutations; p<2×10^-16^ using log-linear regression; see **Figure 2a**) and disk diffusion zone diameter (median 29 mm with no mutations, median 21 mm with one mutation, median 6 mm with two or more mutations; p<2×10^-16^ using linear regression; see **Figure S6**).

**Figure 2.**
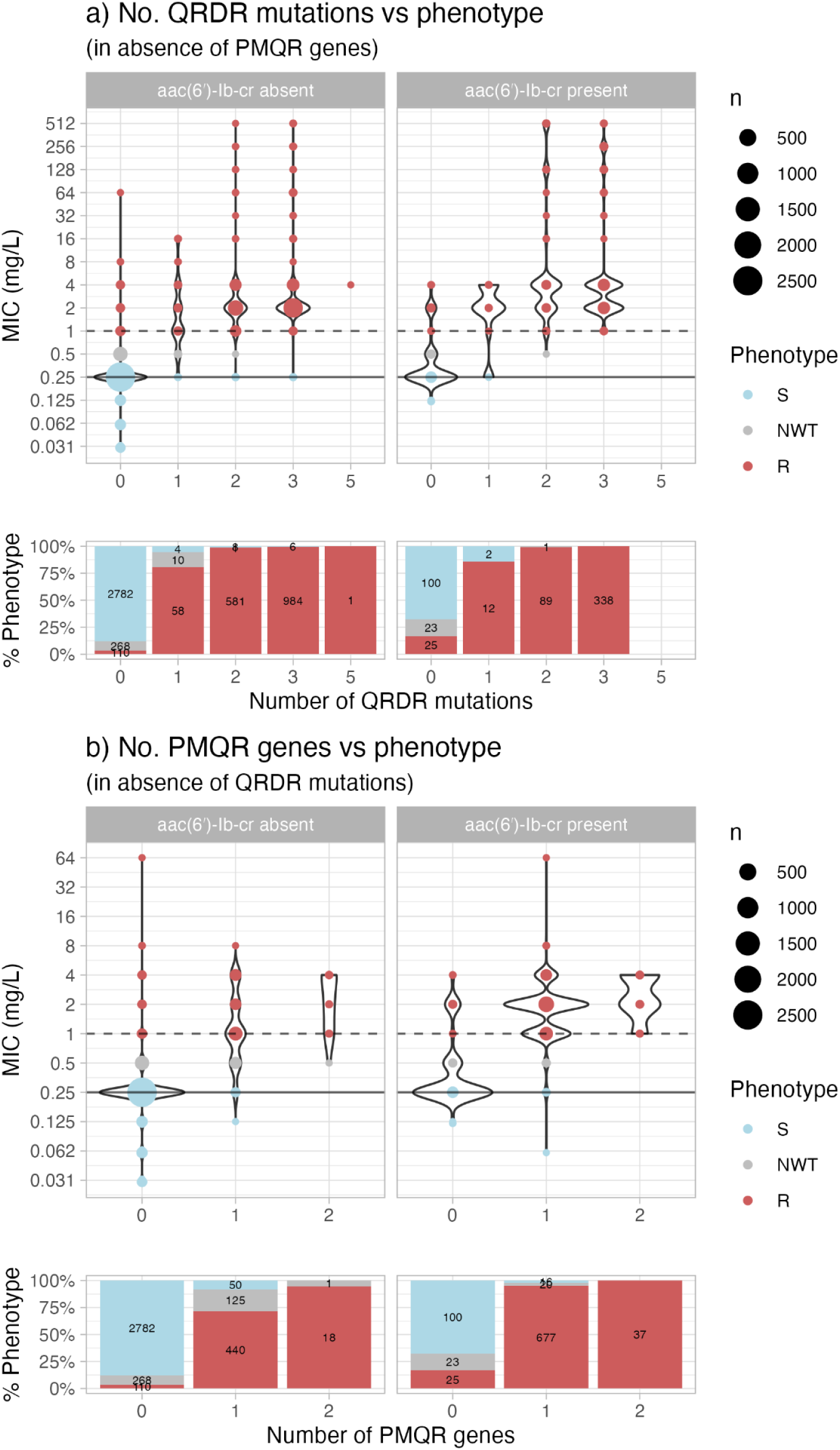
Number of QRDR mutations or PMQR genes vs phenotype, for isolates with MIC measurements (n=9,094). In the absence or presence of *aac(6ʹ)-Ib-cr*, the MIC distributions of genomes with (a) QRDR mutations or (b) PMQR genes and no other fluoroquinolone resistance determinant. The number of susceptible (S), intermediate (I), and resistant (R) genomes is written in each stacked bar and circle sizes represent the number of genomes.

Eleven PMQR genes (ten *qnr* genes and *qepA2)* were identified solo across 706 isolates (i.e., in isolates lacking any QRDR mutations or an *aac(6ʹ)-Ib-cr* gene) (**Figure 1a, Table S6**). *qepA2*, *qnrB7* and *qnrS2* were each observed once amongst such isolates, and all three isolates were ciprofloxacin resistant (MIC 1 mg/L, 1 mg/L, and 4 mg/L, respectively per gene). *qnrS1, qnrB1, qnrB2, qnrB4, qnrB6, qnrB17, qnrB19* and *qnrA1* were each observed alone in ≥12 isolates, and were each independently associated with resistance (odds ratios ≥6, p-values ≤1×10^-3^; see **Table S8, Figures 1b, S7, S8**). Amongst isolates lacking any QRDR mutations or *aac(6ʹ)-Ib-cr* (n=6,401), there was a significant association between presence of any PMQR gene and ciprofloxacin MIC (median 1 mg/L vs 0.25 mg/L, p<2×10^-16^), disk diffusion zone diameter (median 21 mm vs 29 mm, p<2×10^-16^), NWT (OR 115 [87-156], p<2×10^-16^), and clinical resistance (OR 94 [74-120], p<2×10^-16^). The number of PMQR genes was also significantly associated with MIC (median 0.25 mg/L with no PMQR genes, median 1 mg/L with one PMQR gene, median 2 mg/L with two PMQR genes, p<2×10^-16^ using log-linear regression; see **Figure 2b**) and disk diffusion zone diameter (median 29 mm with no PMQR genes, median 21 mm with one PMQR gene, median 10 mm with two PMQR genes; p<2×10^-16^ using linear regression; see **Figure S9**).

Exact matches to *aac(6ʹ)-Ib-cr* reference sequences were found in 2,879 genomes (21.2%; n=2,823 v2 and n=56 v1, see **Methods**) and imprecise matches were found in 29 genomes (0.24%, all v2). Matches to the v2 allele were identified solo, in the absence of other fluoroquinolone resistance determinants, and were associated with resistance (**Figure 1a, Table S7**); v1 was not identified solo. All hits to *aac(6ʹ)-Ib-cr* were combined into a single variable to indicate presence of an *aac(6ʹ)-Ib-cr* gene in all further analyses. In the absence of any QRDR mutations or PMQR genes, presence of *aac(6ʹ)-Ib-cr* was associated with a significant upward shift in the MIC distributions (p=7×10^-14^ using Wilcoxon rank sum test), but the median MIC was the same with or without *aac(6ʹ)-Ib-cr* (0.25 mg/L), and the majority of *aac(6ʹ)-Ib-cr* positive isolates did not exceed the clinical breakpoints (17% R, 33% NWT, 67% S; **Figure 3**). Similar results were evident for disk diffusion diameter (median 23.5 mm vs 29 mm, p=5×10^-5^; 33% R, 58% NWT, 42% S) although the number of genomes were smaller (see **Figure 3**). There was however a significant association between presence of *aac(6ʹ)-Ib-cr* and ciprofloxacin NWT (OR 5 [4-8], p<2×10^-16^), and clinical resistance (OR 9 [6-14], p<2×10^-16^).

**Figure 3.**
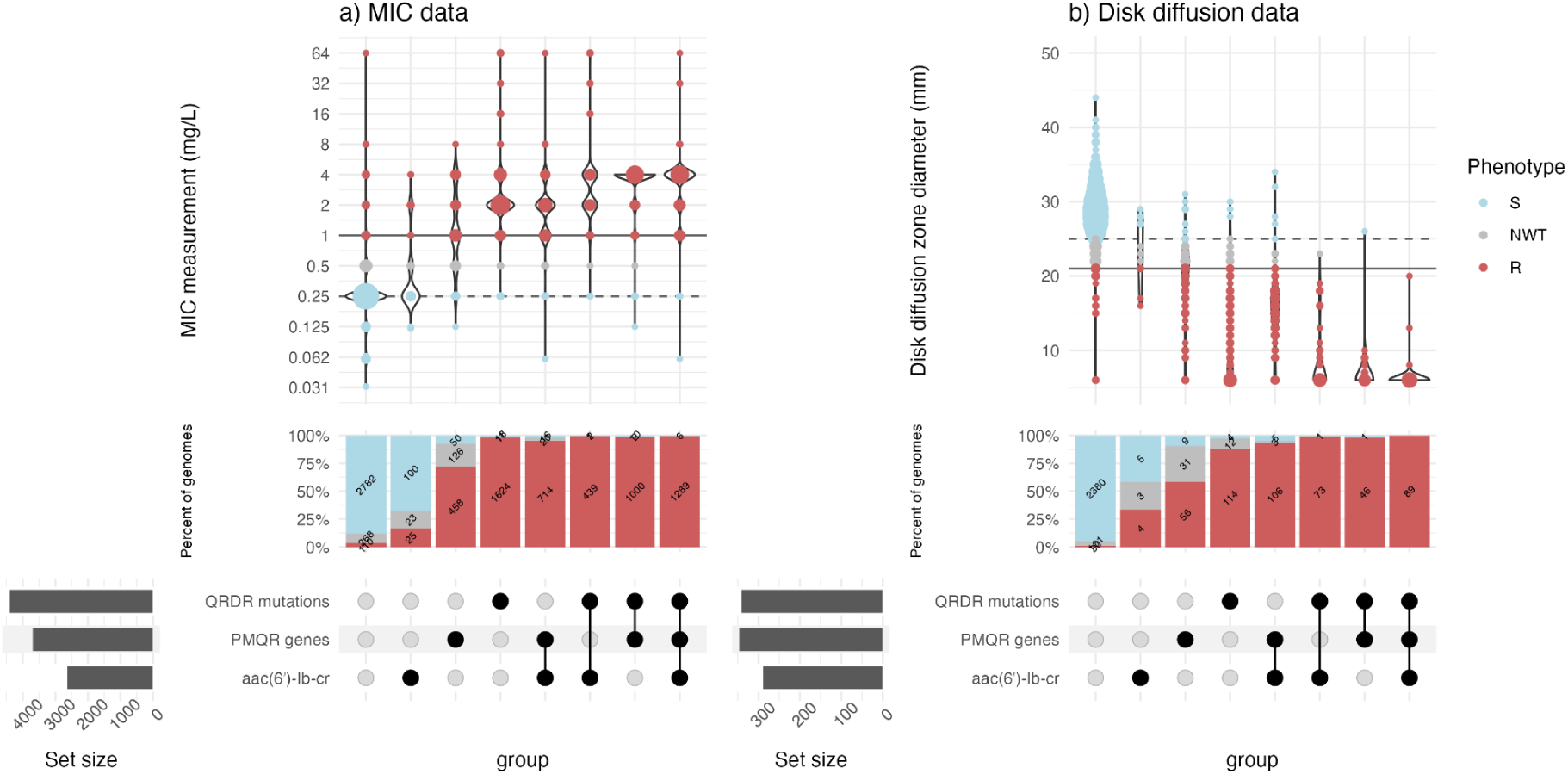
Combined effects of QRDR mutations, PMQR genes, and *aac(6’)-Ib-cr* on ciprofloxacin susceptibility. The (a) MIC or (b) disk diffusion distributions of genomes with QRDR mutations, PMQR genes, or *aac(6ʹ)-Ib-cr*. Included genomes have no other fluoroquinolone resistance determinant. Filled circles in the upset plot indicate the genotype as indicated in the row label. The number of susceptible (S), intermediate (I), and resistant (R) genomes is written in each stacked bar and circle sizes represent the number of genomes.

### Predicting resistance

We explored multivariable logistic regression models for prediction of R or NWT, with predictors being either (1) individual determinants (with minimum frequency ≥20 genomes) or (2) the number of QRDR mutations and number of PMQR genes; with *aac(6ʹ)-Ib-cr* as either (a) an individual predictor or (b) an interaction term (summarised in **Table 1**).

**Table 1.**
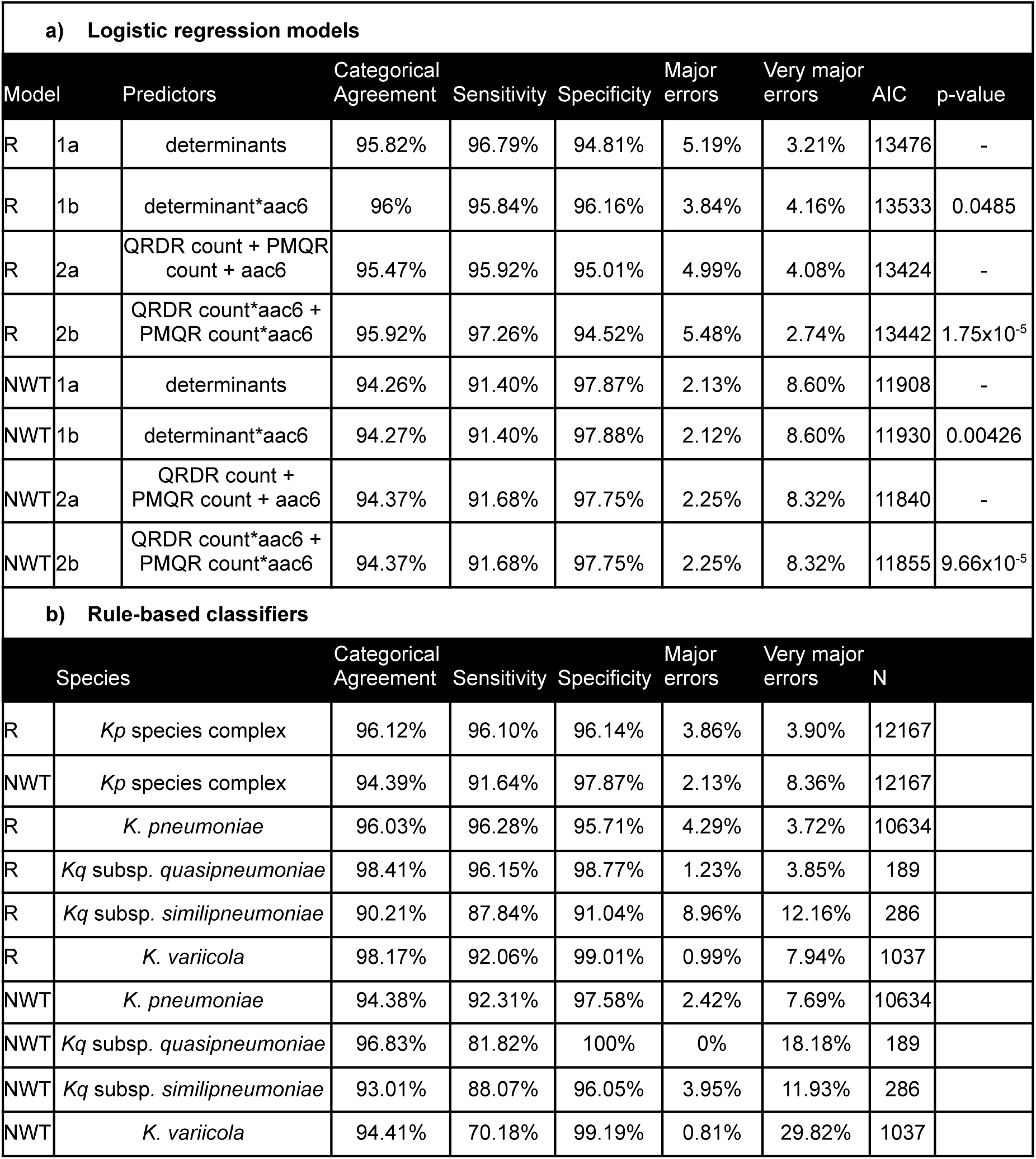
Predictive models for ciprofloxacin resistance. **(a)** Summary of logistic regression models predicting resistance (R vs S/I) or non-wild type (NWT vs WT). Model predictors include determinants (all fluoroquinolone resistance determinants except *aac(6ʹ)-Ib-cr*), aac6 (*aac(6ʹ)-Ib-cr*), the number of quinolone resistance determining region mutations (QRDR count), and the number of plasmid-mediated quinolone resistance genes (PMQR count). For each pair of models (a, b; distinguished by inclusion of interaction terms for *aac(6ʹ)-Ib-cr* in b), model comparison was done using the likelihood ratio test (p-value). AIC, Akaike information criterion. **(b)** Summary of predictive classifiers for ciprofloxacin resistance. Major error (S isolates called as R); Very major error (R isolates called as S); *Kp*, *K. pneumoniae*; *Kq*, *K. quasipneumoniae*.

The individual-determinant multivariable logistic regression models (**Figure 1b, Table S8**) support that most QRDR mutations and PMQR genes with sufficient observations (≥20 genomes) were independently significantly associated with ciprofloxacin resistance (and NWT), consistent with the single variable analyses (**Table S7**). The exceptions were ParC variants, which as noted above were only found in combination with GyrA mutations so are not independently associated with resistance. For the individual-determinant models, those with *aac(6ʹ)-Ib-cr* interaction terms (Model b) provided little evidence of better fit (higher AIC (13,533 for R vs. 11,930 for NWT); p=0.049 for R and p=0.004 for NWT using ANOVA to compare models, see **Table 1**), and only one interaction showed a significant positive correlation with resistance (*qnrB1*.v2:*aac(6ʹ)-Ib-cr*, p=0.0039, see **Figure 1b, Table S8**). The counts-based multivariable logistic regression models (i.e., Models 2a and 2b that include the number of QRDR mutations and PMQR genes in **Table 1a**) support that QRDR mutation count, PMQR gene count, and *aac(6ʹ)-Ib-cr* are each independently associated with ciprofloxacin resistance (**Table S9**). The model with *aac(6ʹ)-Ib-cr* interaction terms provided better fit (p<1×10^-6^ using ANOVA), but the interaction terms were negatively associated with resistance (**Table S9**).

Overall, the regression models showed similar results in terms of resistance classification (**Table 1a**), with high categorical agreement (95.47–95.92%), sensitivity (95.84–97.26%) and specificity (94.52-96.16%). The NWT classification models showed lower sensitivity (91.40–91.68%) and higher specificity (97.75–97.88%); including interaction terms made no difference to classification as WT/NWT (**Table 1a**).

Based on these observations, we developed a rules-based classifier for ciprofloxacin that categorises all genomes into one of ten genotype profiles on the basis of the number of QRDR mutations (0, 1, >1), the number of PMQR genes (0, 1, >1), and the presence/absence of *aac(6ʹ)-Ib-cr* (**Table 2**). Genomes with no determinants, *aac(6ʹ)-Ib-cr* solo, or solo GyrA-87G or GyrA-87H (which had solo PPV <50%, see **Figure 1a**), are predicted as WT S. While having no known ciprofloxacin resistance determinants compared to only having *aac(6ʹ)-Ib-cr* are both predicted as WT S, they are distinguished as separate genotype profiles to show that having *aac(6ʹ)-Ib-cr* solo shifts the R PPV from 2.52% to 18.12% (**Table 2**).

**Table 2.** Simple rules-based classifier for ciprofloxacin R and NWT prediction. Genotype profile describes the number of quinolone-resistance determining regions’ (QRDR) mutations, plasmid-mediated quinolone resistance (PMQR) genes, and presence (1) or absence (0) of *aac(6ʹ)-Ib-cr* identified in a genome (* indicates the gene may be present or absent). ^ GyrA-87G and GyrA-87H are not included in the QRDR count, and *qnrB1* is excluded from the single-PMQR count. N, total genomes with this profile. Prediction describes the predicted phenotype associated with each genotype profile. WT=wildtype, NWT=non-wildtype, R=resistant, S=susceptible. PPV=positive predictive value calculated for each genotype profile, MIC=minimum inhibitory concentration, IQR=interquartile range (based on n=9,094 isolates with MIC data). MIC distributions for each profile are shown in **Figure 5**.

| Genotype profile |  |  | N | Prediction | R PPV | NWT PPV | MIC (mg/L),<br>median [IQR] |
| --- | --- | --- | --- | --- | --- | --- | --- |
| QRDR | PMQR | *aac(6')-Ib-cr* |  |  |  |  |  |
| 0^ | 0 | 0 | 5680 | WT S | 2.52%<br>(n=143/5680) | 9.01%<br>(n=512/5680) | 0.25<br>[0.25-0.25] |
| 0 | 0 | 1 | 160 | WT S | 18.12%<br>(n=29/160) | 34.38%<br>(n=55/160) | 0.25<br>[0.25-0.5] |
| 0 | *qnrB1* | 0 | 160 | NWT I | 43.12%<br>(n=69/160) | 81.25%<br>(n=130/160) | 0.5<br>[0.5-1] |
| 1 | 0 | 0 | 103 | NWT R | 77.67%<br>(n=80/103) | 99.03%<br>(n=102/103) | 1<br>[1-2] |
| 1 | 0 | 1 | 23 | NWT R | 86.96%<br>(n=20/23) | 91.3%<br>(n=21/23) | 2<br>[1-2] |
| >1 | 0 | * | 2164 | NWT R | 99.21%<br>(n=2147/2164) | 99.31%<br>(n=2149/2164) | 2<br>[2-4] |
| 0 | 1^ | 0 | 546 | NWT R | 77.47%<br>(n=423/546) | 94.69%<br>(n=517/546) | 1<br>[1-2] |
| 0 | 1 | 1 | 820 | NWT R | 94.63%<br>(n=776/820) | 97.44%<br>(n=799/820) | 2<br>[1-2] |
| 0 | >1 | * | 68 | NWT R | 97.06%<br>(n=66/68) | 100%<br>(n=68/68) | 2<br>[2-4] |
| >0 | >0 | * | 2443 | NWT R | 99.22%<br>(n=2424/2443) | 99.3%<br>(n=2426/2443) | 4<br>[4-4] |

Genomes with solo *qnrB1* are predicted as NWT I (based on solo PPV, see **Figure 1a**). Genomes with multiple determinants are predicted as NWT R (based on **Figure 3**). Some genotypic categories are *aac(6ʹ)-Ib-cr* agnostic as there is no significant difference in PPV or phenotypic distributions when comparing the inclusion / exclusion of the gene.

Overall performance of the classifier was slightly better than the logistic regression models for prediction of R vs S/I, yielding categorical agreement, sensitivity, and specificity >96%, and ME and VME <4% (see **Table 1b**). As a classifier of NWT vs WT, the rules-based classifier performed very similarly to the logistic regression models, with high specificity (97.87%) and low ME (2.13%), but lower sensitivity (91.64%) and high VME (8.36%) (**Table 1b**).

**Table 2** shows the PPV for the different genotype profiles used by the rules-based classifier; notably the PPV was much lower for genotype profiles with a solo QRDR mutation (77.67% PPV for R, 99.03% for NWT) or solo PMQR gene (77.47% PPV for R, 94.69% PPV for NWT) compared with genotype profiles comprising multiple determinants (**Table 2, Figure S10**). Most of the ‘misclassifications’ (95%, n=220/231, across 18 datasets of 12,167 isolates) were caused by isolates with assay measurements lying between the clinical breakpoints (0.5mg/L MIC, 22/23 mm disk diffusion zone diameter), but classified genetically as NWT R (see **Figure 4**). In most cases, these genomes carried a single QRDR mutation (9%, n=20), solo *qnrB1* (13%, n=28), solo *qnrS1* (63%, n=138), or another solo PMQR gene (11%, n=24), and as such the phenotype may be explained by poor expression and/or genomic context of the acquired gene.

**Figure 4.**
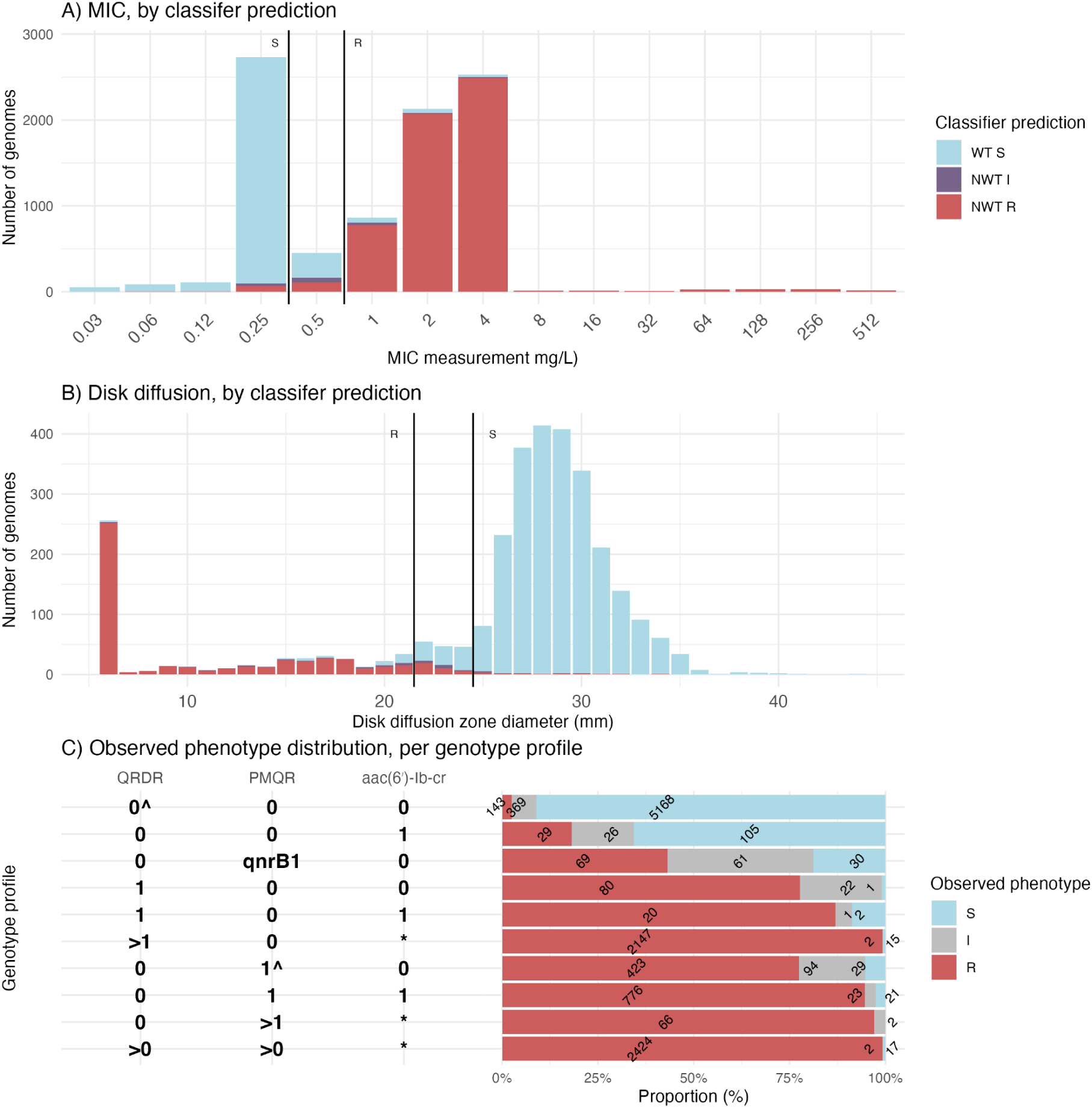
Ciprofloxacin AST distributions of genomes coloured by the phenotype category predicted by the rule-based classifier. The (A) MIC and (B) disk diffusion distribution of genomes coloured by the rules-based classifier prediction. Vertical lines show EUCAST ciprofloxacin breakpoints. C) Frequency of true, observed phenotypes observed for genomes falling into each genotype profile used in the classifier. The rules-based classifier assigns each genome into a genotype profile depending on the number of quinolone resistance determining region mutations (QRDR), and the number of plasmid-mediated quinolone resistance genes (PMQR), and the presence (+) or absence (-) of aac6 (*aac(6ʹ)-Ib-cr*). ^ indicates that GyrA-87G and GyrA-87H are not included in the QRDR count, and qnrB1 is excluded from the single-PMQR count. The number of susceptible (S), intermediate (I), and resistant (R) genomes is written in each stacked bar.

Figure 5 shows the MIC distribution for the different genotype profiles used by the rules-based classifier, and **Table 2** includes the median MIC and interquartile range for each profile. These, together with the PPV, provide useful supportive evidence to aid interpretation of a specific genotype profile, alongside the categorical classifications of WT/NWT and S/R.

**Figure 5.**
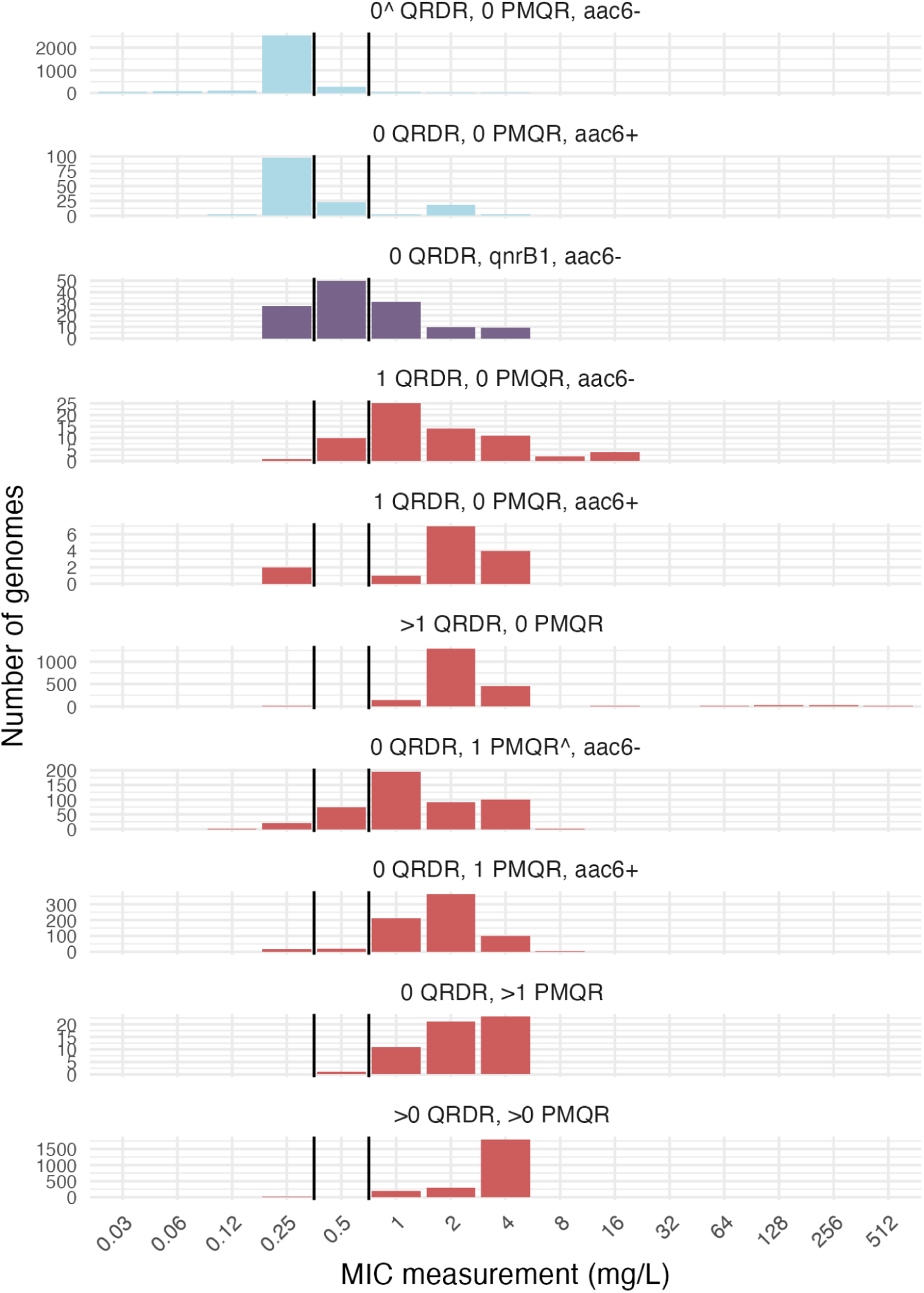
Ciprofloxacin MIC distribution for each genotype profile category used by the rule-based classifier. Vertical lines indicate the S and R breakpoints. The rules-based classifier assigns each genome into a genotype profile depending on the number of quinolone resistance determining region mutations (QRDR), and the number of plasmid-mediated quinolone resistance genes (PMQR), and the presence (+) or absence (-) of aac6 (*aac(6ʹ)-Ib-cr*). ^ indicates that GyrA-87G and GyrA-87H are not included in the QRDR count, and *qnrB1* is excluded from the single-PMQR count. Colours represent the rules-based classifier prediction: blue = WT S, purple = NWT I, and red = NWT R.

As this dataset (n=12,167 genomes) included all *Kp*SC (comprising 1,992 distinct STs), we assessed performance separately for each species with ≥100 genomes. This yielded very similar metrics for each species, although performance was notably better for *K. pneumoniae* (which accounts for 87% of the data), *Kqs,* and *Kvv* (see **Table 1b**). We also assessed performance within subgroups (STs, countries, source type, AST platform type, host infection status) (Figure 6). Sensitivity and specificity were high (>89%) regardless of source (human, animal, environment), AST platform type, and host infection status. Using different semi-automated AST platforms varied performance metrics (e.g., across n=3,620 isolates from three studies assessing MICs using the Phoenix platform, categorical agreement was 97.62%, with 2.37% ME and 2.38% VME; compared with respective values of 95.28%, 8.77% and 3.12% across n=4,640 isolates from 13 studies assessing MICs using the Vitek platform). Categorical agreement, ME, and VME values across n=3,073 isolates from six studies using the disk diffusion method were 96.65%, 1.96%, and 10.23%, respectively.

**Figure 6.**
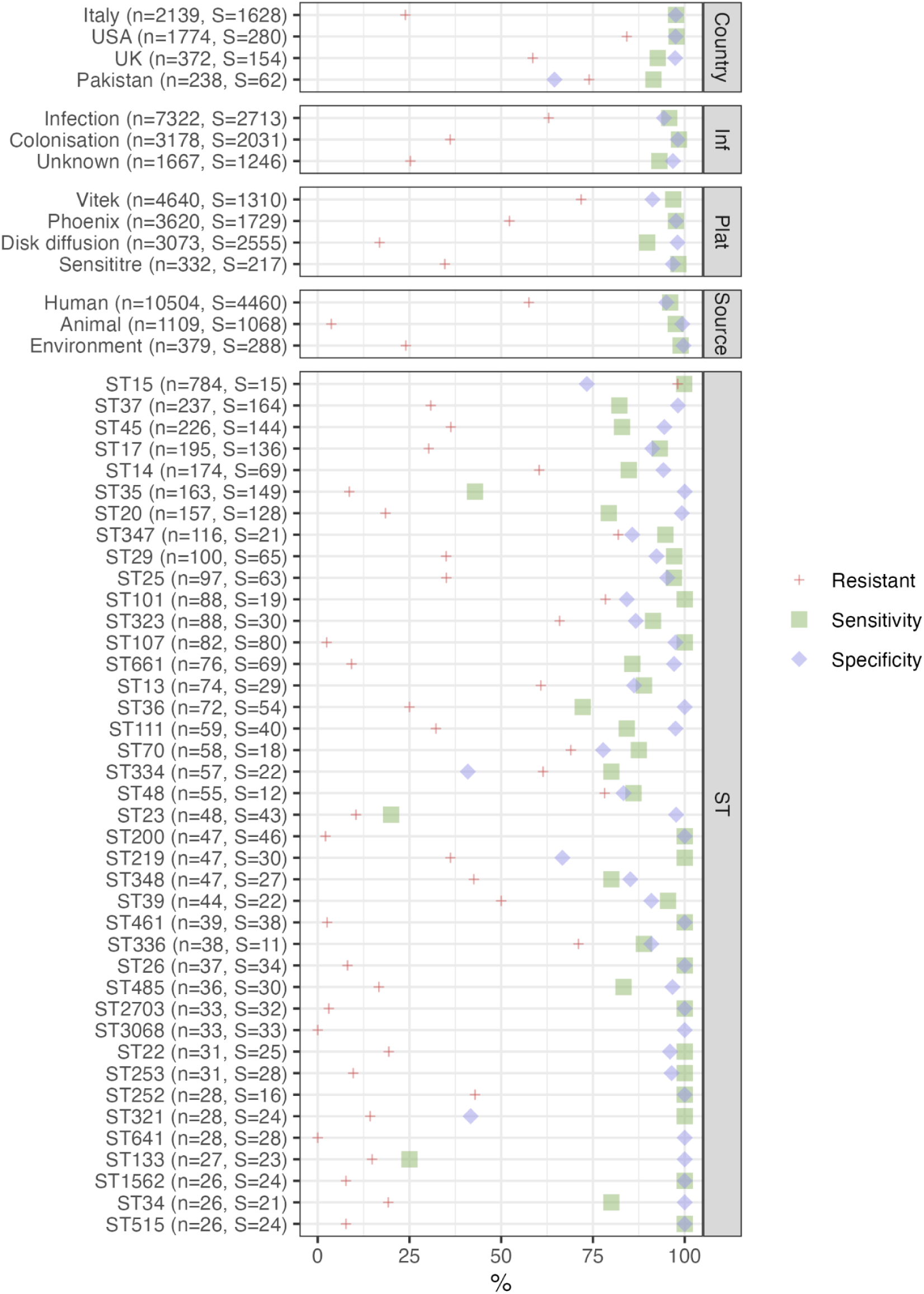
Sensitivity and specificity of the rules-based classifier stratified by isolate metadata. Sensitivity and specificity were calculated after the isolates were categorised in terms of geography, infection status (Inf), source, antibiotic susceptibility testing platform type (Plat), and sequence type (ST). The percent of resistant genomes of each category is represented by a red cross. The total number of genomes (n=) and number of susceptible genomes (S=) are printed next to the metadata category. Only metadata categories with more than 10 susceptible genomes, >25 total genomes, and have representation from three or more datasets are included. Datasets with <100% resistant genomes are displayed (since 100% resistant genomes result in 0% specificity).

There is also some variability in performance metrics across geographical sources (average sensitivity 94.8%, average specificity 89.2%) and STs (average sensitivity 87.9%, average specificity 98.0%).

### Unexplained phenotypes

There were 140 phenotypically resistant isolates for which we did not detect any QRDR mutations, PMQR genes, or *aac(6ʹ)-Ib-cr* (i.e., n=140/6,177, 2.3% of resistance phenotypes).

Over half of these isolates (n=73, 52%) had borderline phenotype assay results (n=53 with MIC=1 mg/L and n=20 with disk diffusion zone diameter 20-21 mm), and were from diverse lineages (n=84 unique STs across 140 isolates). Amongst the 140 isolates with unexplained resistance, n=11 had *ompK35* loss/truncation, n=4 had *ompK36* loss/truncation, n=23 had two copies of *oqxA*, n=28 had two copies of *oqxB*, n=31 had inactivating mutations/truncations/loss in *oqxR, rob, rarA* or *norB*, and n=74 (53%) had none of these features detected (none of which had borderline assay results, i.e., 22/23 mm disk diffusion zone diameter or 0.5 mg/L MIC). Of these features, only *ompK35*/*ompK36* loss was significantly associated with resistance amongst the n=5,671 genomes lacking QRDR mutations and *qnr/qep* or *aac(6ʹ)-Ib-cr* genes (see **Table S10**), but these had a low PPV (<25%). Overall, these results do not support expansion of the classifier to include additional genetic determinants.

Forty-one phenotypically susceptible isolates carried a QRDR mutation (other than GyrA-87G or H) and/or >1 *qnr*/*qep* gene (other than *qnrB1*). Most of these isolates (n=30, 73%) had borderline phenotype assay results (n=30 with MIC=0.25 mg/L and n=4 with disk diffusion zone diameter 25-26 mm), and they were diverse in terms of ST (n=33 unique STs). Half of these genomes (n=23) carried *qnrS1* and no other known determinants; the rest harboured a single GyrA-83 mutation (n=1 GyrA-83F, n=1 GyrA-83I, n=2 GyrA-83Y), a single PMQR gene (n=1 *qnrA1*, n=3 *qnrB2*, n=3 *qnrB4*) or multiple variants (n=8). Two genomes lacked both *oqxA* and *oqxB,* and one genome lacked *oqxB*, which could potentially explain the unexpected susceptible phenotype.

### External validation of the rules-based classifier

We undertook independent external validation using 19 datasets not available at the time of developing the rules-based classifier (n=6,763 *K. pneumoniae* and n=267 *Kvv* sourced from 6 continents, testing S or R via either semi-automated AST / broth microdilution methods (n=5,835), disk diffusion (n=1,155) or gradient diffusion (n=40) (see **Methods** and **Table S4**). Note that to reduce outliers due to uncertain (and often irreproducible, see re-testing data above) classification of isolates based on borderline assay results, isolates with an I phenotype (MIC=0.5 mg/L, n=491 or disk zone 22-25, n=172) were excluded from the primary analysis. A majority (94%, n=624/663) of these I isolates had either no ciprofloxacin resistance determinants (n=255, predicted S), *aac(6ʹ)-Ib-cr* only (n=92, predicted S), *qnrB1* only (n=64, predicted I), or one other PMQR (n=213, predicted R).

The pooled performance metrics were 93.12% categorical agreement [95% CI, 92.50-93.74%], 8.65% ME [95% CI, 7.34-9.97%], and 6.20% VME [95% CI, 5.51-6.90%], however there was wide variability across datasets (Figure 7**, Table S11, Figure S11**). Median values per dataset (for 14 datasets with sufficient data, see **Methods**) were 93.44% categorical agreement (range, 73.97-100%), 9.46% ME (range, 0-60%), and 6.20% VME (range, 0-21.62%). One dataset (MBIRA study using Vitek, labelled H in Figure 7) was an outlier with very low categorical agreement (74.68%) and high ME (43.12%). This was attributable to n=37 ST1741 isolated from the same hospital in Tanzania in 2021, with MIC=0.25 mg/L (therefore classified as S) but harbouring *qnrS1* (therefore predicted as R).

**Figure 7.**
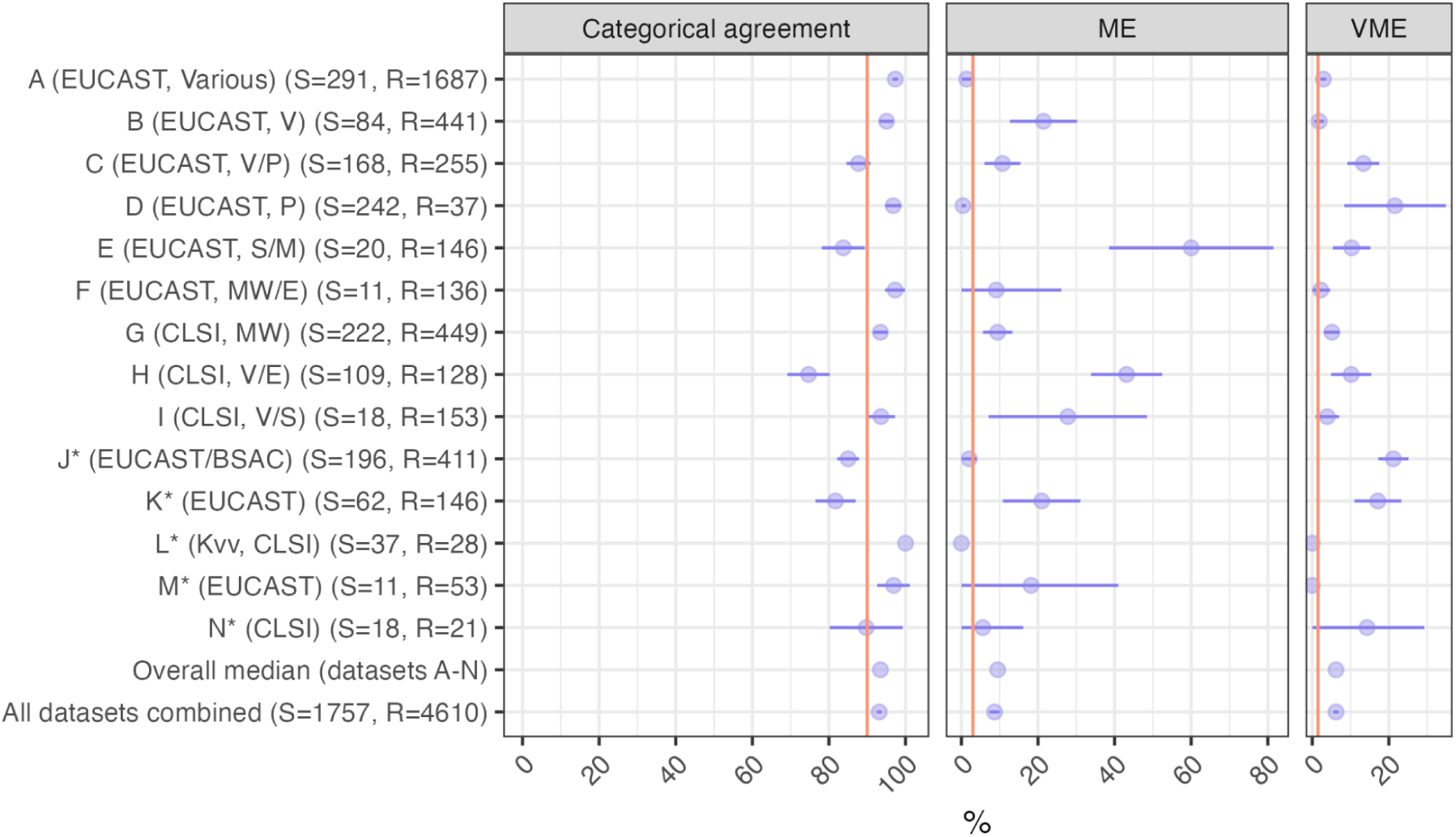
External validation of rules-based classifier. Individually plotted datasets have at least 25 high quality genomes with calls of S or R, including more than 10 of each category (S/R). Ciprofloxacin susceptibility phenotypes were measured using MIC or disk diffusion (indicated by *). The breakpoint guidelines used are indicated in the first bracket, followed by MIC method (Vitek (V), Phoenix (P), Sensititre (S), Microscan Siemens (M), Microscan WalkAway Beckman Coulter (MW), and E-test (E)). Dataset A includes data from laboratories across Europe and they used various MIC methods. The number of S and R isolates in each dataset are listed in the second bracket (S=/R=); isolates with I phenotype were excluded as they are within 1 doubling dilution of either S or R. ‘Overall median’ represents the median values for each metric, across all plotted datasets, A-N. ‘All datasets combined’ represents the values calculated by pooling S and R isolates from all available validation datasets (including datasets A-N and those that were excluded from individual plotting based on the criteria noted above). Plotted horizontal lines indicate 95% confidence interval. Plotted orange vertical lines represent the United States Food and Drug Administration AST Systems Guidance targets for categorical agreement (>90%), major errors (ME, <3%), and very major errors (<1.5%).

We hypothesise this reflects a single clone with reduced expression of *qnrS1* resulting in incomplete penetrance of the expected phenotype, whose repeated isolation in the local study is skewing the overall metrics for this dataset (for which it comprises 15% of the sample).

When the 37 isolates are excluded from the dataset, the categorical agreement increases to 90.19% and ME lowers to 20.83%. Another dataset (Oxfordshire study using Phoenix, labelled D in Figure 7) was an outlier with very high VME (21.62%), where all 8/37 of the resistant isolates (with no ciprofloxacin resistance determinants therefore classified as S) had MIC=1 mg/L, which is within one doubling dilution of the intermediate phenotype (MIC=0.5 mg/L). Few of the datasets met the US FDA target for ME <3% (n=4) or VME <1.5% (n=3). Median MICs for each genotype profile are the same for the rules-based classifier dataset (**Table 2**) compared to the pooled validation datasets (**Table S12**) with the exception of the one QRDR and *aac(6ʹ)-Ib-cr* profile (MIC 2 mg/L vs. 1 mg/L, respectively). Evaluation metrics for predicting R (including / excluding I) and NWT per dataset are detailed in **Table S11** and PPVs and median MICs of genotype profiles per dataset are listed in **Table S12**.

### Global distribution of ciprofloxacin resistance determinants

To assess the global genomic epidemiology of ciprofloxacin resistance determinants in a larger scale compared to our genotype-phenotype dataset, we downloaded publicly available Kleborate typing results for *Kp*SC genomes from Pathogenwatch (v23.4.4) (n=31,319 following cluster de-replication, see **Methods**) and applied our rules-based classifier to predict resistance. Very high fractions of *K. pneumoniae* genomes were predicted as ciprofloxacin R (68.0%) and NWT (69.8%), whereas other *Kp*SC had much lower predicted resistance rates: *Kvv* 14.8% R and 15.2% NWT, *Kqq* 54.2% R and 54.2% NWT, *Kqs* 46.5% R and 46.5% NWT, *K. africana* 0% R and 0% NWT. QRDR mutations were common in *K. pneumoniae* (identified in 50.3%), and were universally present in the most common clones (ST258, ST11, ST307, ST147, and ST15; represented by n>1,200 cluster de-replicated genomes each), which all had at least two fixed QRDR mutations (see **Figure S12**). In contrast, QRDR mutations were rare in other species (0-3.2%), including the most common clones (**Figure S13**). PMQR genes and *aac(6ʹ)-Ib-cr* were rare in *Kvv* (13.8% and 6.4%, respectively) and *K. africana* (0% and 0%), but more common in *K. pneumoniae* (37.0% and 27.1%), *Kqq* (50.0% and 23.2%) and *Kqs* (41.0% and 22.2%). Interestingly, *K. pneumoniae* ST258 nearly all lacked PMQR genes and *aac(6ʹ)-Ib-cr* (3.5% and 0.6% prevalence, respectively), whilst PMQR genes were more common in *K. pneumoniae* clones ST11, ST307, ST147, and ST15 (33.3-66.1%). For the most common *Kvv* clone, ST347, almost half would be predicted resistant (40%, n=10/25), driven by the combination of *qnrB1*/*qnrB4* and *aac(6ʹ)-Ib-cr* (70%, n=7/10) or *qnrS1*/*qnrB2* alone (30%, n=3/10); however the other common *Kvv* clones (with >10 genomes each) largely lacked these resistance determinants (80-100%) (**Figure S13**).

Genotype profiles varied by year, with an overall trend towards accumulation of resistance determinants, from <10% of genomes with any QRDR mutation or PMQR gene in 2000, to >30% frequency for each type since 2015, and >10% of genomes carrying both QRDR and PMQR since 2012 (**Figure S14**). After 2010, over 50% of genomes have a predicted MIC of >0.5mg/L, which would be interpreted as resistant (**Figure S14**). Notably, from 2000-onwards, the proportion of genomes with an MIC prediction of ≥4 mg/L generally increases over time (**Figure S14**). The available genome sample varied widely by country, however there were 40 countries with >100 genomes isolated from 2000 onwards. Using the rules-based classifier to predict resistance, there was a significant positive association between national quinolone consumption in the period 2000-2018 (R^2^=0.2, p=0.0044) (Figure 8). Interestingly, national consumption was a strong positive predictor of the mean number of QRDR mutations per *K. pneumoniae* genome (R^2^=0.23, p=0.0025) but was not associated with the mean number of PMQR genes (R^2^=2×10^-4^, p=0.93) (Figure 8).

**Figure 8.**
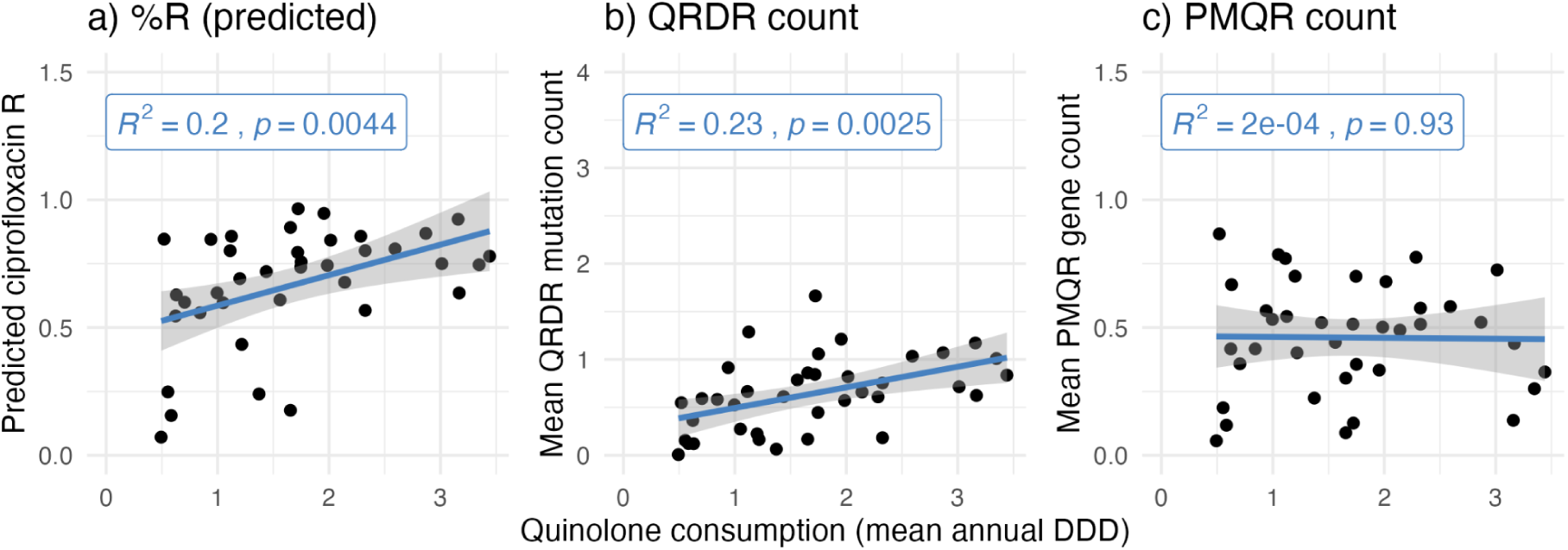
Association between ciprofloxacin genotypes and consumption, by country. Each data point represents a country for which at least 100 public *K. pneumoniae* genomes isolated from 2000 onwards were available in Pathogenwatch (n=109 countries). Plots show mean annual quinolone consumption (measured as Defined Daily Doses (DDD) per 1000 inhabitants per day) for the period 2000-2023, on the x-axis, estimated in the GRAM study^59^. Y-axes show the genome-derived data on ciprofloxacin resistance determinants, for genomes isolated from 2000 onwards. QRDR=quinolone resistance determining region, PMQR=plasmid-mediated quinolone resistance, R=resistance, predicted by applying the rule-based classifier in Kleborate.

## DISCUSSION

We aggregated a diverse collection of 12,167 *Kp*SC genomes with matched ciprofloxacin susceptibility assay results from 26 study groups. This global collaboration allowed us to assemble a much larger data set than has previously been used to investigate AMR genotype-phenotype relationships in *Kp*SC (more than three times the size of previous reports). Importantly, the dataset represents a very diverse set of *Kp*SC isolates, isolated from 27 countries (across all continents except Antarctica), sampled from a variety of human, animal, and environmental sources, and comprising 1,992 distinct STs. Using this collection, we explored the distribution and phenotype associations of known determinants of fluoroquinolone resistance. We used these observations to develop a rules-based classifier that performed at least as well as logistic regression models, implemented this classifier into the widely-used Kleborate software v3.2.4, and externally validated it using 7,030 additional genomes from 19 studies.

We focused on mutations and genes that have been shown to influence ciprofloxacin susceptibility through known mechanisms of action. Our exploratory analyses supported generalising and expanding the genotype-phenotype association by taking into account the number of QRDR mutations, the number of PMQR genes (with a few specific exceptions), and the presence/absence of *aac(6ʹ)-Ib-cr*. Notably, logistic regression models using these generalised counts performed nearly identically to more complex models utilising individual determinants; we therefore applied the counts-based approach to our rules-based classifier, which greatly simplifies the calculation of predictions. In a previously published MIC prediction model for *K. pneumoniae,* the features most correlated with ciprofloxacin MICs were integron integrase IntI1, tryptophan synthase, and TEM β-lactamase^45^. This is likely due to confounding by genetic linkage in the multi-drug resistant clones ST307 and ST258 which dominated the dataset. The same study compared the ability of machine learning based on genome-wide *k*-mers vs AMR genes alone to predict MIC and found near-identical results across all methods. This supports our simple and interpretable approach of utilising proven AMR determinants to predict ciprofloxacin resistance.

Similar approaches have been taken previously to understanding and predicting ciprofloxacin resistance in *Escherichia coli*^61,62^. The most recent and large-scale analysis in *E. coli* (n=2,875 isolates) reported cumulative effects of QRDR mutations and PMQR genes on MIC^62^. In *E. coli*, no solo variants were associated with clinical resistance, likely because *K. pneumoniae* has a higher background (WT) ciprofloxacin MIC (tentative ECOFF 0.125 mg/L [CI 0.06–0.25 mg/L], compared with tentative ECOFF 0.06 mg/L [CI 0.03–0.06 mg/L] for *E. coli*) due to its core efflux pump OqxAB, therefore a single resistance determinant need only increase the MIC by one or two doubling dilutions to exceed the tentative ECOFF. Here, we aimed to predict categorical resistance (WT S, NWT I, and NWT R) rather than MIC values, so that we could increase sample size by utilising disk diffusion data (>3,000 isolates) and avoid the unresolved challenges associated with modelling of MIC data^45,47^, while meeting the needs of most users.

Compared with the logistic regression models we constructed, the rules-based classifier yielded slightly better agreement and has the benefit of being transparent and easier to understand and interpret, especially compared with models that have interaction terms. Similarly, rules-based classification is easier to apply as an interpretation layer to generate predictions from genotypes, requiring simple logic that is easier to encode into bioinformatics or reporting pipelines than regression models are (again, especially compared with models that have interaction terms, which provided the best fit amongst the regression models).

Additionally, the approach of classifying into multiple combinatorial genotype profiles, each associated with a prediction, allows for more precise communication of the strength of evidence supporting each prediction, which can vary substantially for different profiles that are associated with the same predicted phenotype category. For example, the PPV associated with a prediction of R based on presence of a single PMQR gene (77%) is much lower than a prediction based on presence of PMQR plus *aac(6ʹ)-Ib-cr* (95%), or two PMQR genes (97%). Similarly, although we did not specifically aim to predict MIC, the genotype profiles used by the classifier are each associated with a specific MIC (Figure 5), which can be reported alongside the categorical prediction. For example, the MIC associated with presence of a single PMQR gene (1 mg/L [IQR, 1–2 mg/L]) is lower than a prediction based on the presence of PMQR plus *aac(6ʹ)-Ib-cr* (2 mg/L [IQR, 1–2 mg/L]), or two PMQR genes (2 mg/L [IQR, 2–4 mg/L]).

While our dataset was dominated by *K. pneumoniae* (87%), the rules-based classifier showed similar performance for other members of the species complex (**Table 1b**, Figure 7), although the sample sizes available to validate this are currently too small to provide high confidence. The classifier showed equivalent performance in isolates from humans, animals, and other sources, and in clinical and non-clinical isolates (Figure 6), suggesting good generalizability. Similar performance metrics were observed across diverse lineages (STs), although there was some variability which warrants further exploration into potential lineage effects, as has been recently reported for amoxicillin-clavulanic acid resistance in *E. coli*^63^ (e.g., there may be lineage-associated variation in *oqxAB* regulation in the *KpSC*).

The rules-based classifier was implemented in Kleborate^50^ (v3.2.4), a widely-used *Klebsiella* genotyping tool that is available as an open-source command-line tool and is integrated into several microbial genomics platforms including Bactopia^64^, Pathogenwatch^58^ and BIGsdb^65^.

This integration facilitated external validation on datasets outside the core consortium, without the need for testers to share their unpublished primary data, and yielded similar performance estimates as the discovery set.

Kleborate output includes not only the categorical prediction (WT/NWT, S/I/R) but also the genotype profile on which the categorical prediction is based, and the PPV and MIC values associated with that profile, as shown in **Table 2**. These data are intended to provide additional information to users with which to interpret the prediction, and is based on the 12,167 genomes reported in this paper (all of which are publicly available; i.e., it does not include additional information from the external validation data sets as this is not currently reproducible). It is envisioned that future updates of Kleborate will include enhanced quantitative data (PPV and MIC) to support the interpretation of genotype profiles, as more data is amassed and published. It is also envisioned that future versions of Kleborate will include predictions for other drugs, based on extension of this work by the KlebNET-GSP AMR Genotype-Phenotype Group.

Our rules-based classifier includes only determinants with proven mechanisms of action, which were independently associated with resistance in our study (Figure 1, **Tables S6-7**). It is possible that other variants contribute to predictable variation in ciprofloxacin resistance (e.g., upstream structures that can change expression of *aac(6ʹ)-Ib-cr*^66^ or *qnr*^67^ genes). Other considerations include investigating the genomic context of genes (i.e., plasmid or chromosome), in addition to plasmid copy numbers. The above mentioned genomic investigations are out of scope for this study, but we considered other possibilities and found no evidence to support their inclusion in a classifier, although novel mechanisms may remain to be discovered.

Notably, half of the mis-classified genomes had phenotype measures close to the resistant clinical breakpoint, and our available re-test data suggests that it is quite likely that some discrepant classifications result from mismeasured or imprecise (i.e., close to breakpoint) phenotypes rather than undiscovered genetic markers. The diverse methods and sources of phenotype measures and the inability to re-test all isolates with unexpected phenotypes (as some were not stored, or were not recoverable) are key limitations of our study. Here, we found that re-testing n=150 of 252 isolates with unexpected phenotypes (R but without known determinants, or S with known determinants) prior to modelling resulted in ‘correcting’ of phenotypes for 118 (79%). The high error rate identified amongst isolates that we were able to re-test, highlights the potential impact of re-testing on evaluation metrics, and the importance of considering phenotype measurement inaccuracies (particularly those close to the intermediate phenotype clinical breakpoints)^44,68,69^ when planning and interpreting genotype-phenotype studies. It is also therefore reasonable to expect that re-testing on a wider scale might improve the estimated error rate of the classifier. Another explanation of phenotype measurements changing after retesting (especially near the clinical phenotypic breakpoint) is that there could also be true fluctuation if there is variation in gene expression or plasmid copy number. In line with the importance of phenotype data quality, categorical agreement and error rates differed between studies (see Figures 6**, 7, S11**), and by semi-automated AST platform which demonstrates the variance between phenotypic platforms^70–74^ (Figure 6**, 7**). Assessment of performance could also be strengthened by using larger datasets with consistent phenotyping as gold-standard for comparison, with biological repeat measures for each isolate.

Our analyses highlight that a very large and diverse sample size is needed to investigate the association between individual and combinatorial effects of resistance determinants on resistance. Importantly, many of the published *Klebsiella* genome studies source their data from a hyperendemic disease or clonal outbreak, therefore while the sample size may be large, the datasets are not diverse. More data is needed from non-*K. pneumoniae* species, and in particular we did not have sufficient data to assess *K. quasivariicola*, *K. variicola* subsp. *tropica,* or *K. africana*. Lastly, another limiting factor is the need to combine phenotypic data from different laboratories that used different methodologies. To mitigate this issue, we re-interpreted all MIC and disk diffusion zone diameters using the same interpretive criteria and directly explored the association between genotype profiles and assay values, and excluded studies that were only able to contribute categorical data (S/I/R) as such data could not be consistently interpreted. This highlights the importance of storing primary AST measures, and not just interpreted categories, when extracting and archiving data for AMR research. Future research of this nature, and the deposition of AST data alongside WGS data in public archives, would benefit from availability of standard tools and formats for storing and sharing AST data^73^. Having standardized templates for antibiogram data deposition would in turn promote the accurate collection of methods and other metadata to be shared with public data repositories. Here, we aggregated and processed all AST data in the NCBI antibiogram format (https://www.ncbi.nlm.nih.gov/biosample/docs/antibiogram/), however we note that many partners did not have the in-house capacity to transform their primary data into this format.

While we are not proposing the use of WGS nor the simple rules-based classifier as a replacement for phenotypic AST, it is illustrative to compare the outputs to the accepted standards. Currently, the US FDA targets are >90% complete category agreement, <3% ME, and <1.5% VME^55^. Of the 14 datasets with sufficient data, the external validation results reported the median values of 93.44% categorical agreement (range, 73.97-100%), 9.46% ME (range, 0-60%), and 6.20% VME (range, 0-21.62%) for predicting R, see Figure 7), which exceed the criteria for categorical agreement but are far from the target ME and VME. The ranges for ME and VME indicate there are datasets for which the rules-based classifier performs without errors, which shows the potential of using WGS and the rules-based classifier for ciprofloxacin resistance prediction.

While the evaluation metrics provide a general observation for the performance of the rules-based classifier, it is essential to investigate each dataset individually because their sampling can drive evaluation metrics to extreme values. For example, all four *Kvv* datasets were small (<100 genomes) and had mostly (>50%) susceptible isolates, therefore sensitivity and VME are easily affected by a few false negatives. Generally, small datasets with an imbalance of S/R isolates can fluctuate evaluation metrics widely given only a few misclassifications. In addition, as seen with the H (CLSI) dataset, there may have been over-representation of a ST1741 clone with one PMQR gene incorrectly predicted as R, which resulted in the poor specificity and ME metrics. The ability to investigate the sampling and other genotypic information highlights the importance of having access to genomes and metadata to establish potential reasons (i.e., sampling biases) for poor evaluation metrics.

Similar to the development of the rules-based classifier, increased sampling from all *Kp*SC members and AST data collected from disk diffusion methods in the external datasets would have provided a more comprehensive view of the classifier performance.

To our knowledge, this is the first investigation of ciprofloxacin resistance using the public *Kp*SC data from Pathogenwatch, where we use our rules-based classifier to predict resistance and MICs based on genotype and correlate with national annual quinolone consumption data. We attempted to mitigate sampling bias by grouping highly similar isolates (e.g., matched genotype and metadata) from the same study and selecting one representative. However, we highlight that sampling bias will still exist as studies often select for resistant isolates (often through hospital outbreak monitoring) as opposed to unbiased sequencing of all isolates, which can drive the observed patterns of increased predicted resistance/MICs and ciprofloxacin resistance determinants since 2000. Similar to the KlebNET-GSP collection, the public data consists mostly of *K. pneumoniae*, calling attention for researchers to also select for the sequencing of other *Kp*SC species. As the public data does not include matched antibiogram data, we were unable to validate the rules-based classifier using this collection nor were we able to compare the predicted resistance/MIC to true observed phenotypes. In general, we advocate for all study groups to share antibiogram data along with their WGS public deposition as a standard template is readily available through the NCBI and ENA platforms. As such, we have publicly shared all ciprofloxacin antibiogram data for the 12,167 *Kp*SC genomes used in the development of the rules-based classifier (as a part of the KlebNET-GSP AMR Genotype-Phenotype Group collection).

Our global consortium gathered a diverse collection of isolates to study the contribution of individual and combinatorial resistance determinants in AMR genotype-phenotype relationships. We show the accessible implementation of predictors to facilitate dissemination and external validation of predictive algorithms. This approach will be applied by the KlebNET-GSP group to study other antimicrobials of clinical importance in treating *K. pneumoniae* infections, ideally with increased sample size for other species complex members, and we envisage re-evaluating and refining the rules-based classifier for ciprofloxacin as more evidence is accumulated. A similar approach has been applied recently to *Mycobacterium tuberculosis*, via the World Health Organization’s TB catalogue^75^, and has potential to be expanded to all WHO Priority Pathogens^76^.

## DECLARATIONS

## CONSENT FOR PUBLICATION

Not applicable

## AVAILABILITY OF DATA AND MATERIALS

The KlebNET-GSP AMR Genotype-Phenotype Group discovery dataset used to develop the rules-based classifier are all published with accessions in **Table S1**-**S3**. Information about the external validation datasets can be found in **Table S4-S6**. The ciprofloxacin resistance rules-based classifier is implemented for public use in Kleborate (v3.2.4) and Pathogenwatch (v23.4.4). Pathogenwatch *Kp*SC genomes and metadata were downloaded (https://cgps.gitbook.io/pathogenwatch/public-data-downloads) on August 11, 2025.

All code and analyses are available at github.com/klebgenomics/cipropaper.

## AUTHOR CONTRIBUTIONS

**Conceptualization** - EJF, KLR, INO, IHL, AF, ND, MH, AE, NF, PM, ØS, GDW, AGM, SWL, LM, SS, TL, MS, SB, AJM, PD, DMC, TW, KS, MET, NRT, SB, PT, MB, BH, AA, FL, JRI, AM, TMC, RC, JS, SR, TG, KY, PNAH, LGCP, AW, NAF, EH, KW, TGC, KEH **Data curation** - IA, HT, JC, GN, VS, AOO, OI, MAKH, AC, AC, MH, AC, HMBS, YG, KLW, SWL, LM, SS, TL, MS, CR, AR, AJM, KB, PD, FDM, KS, BH, RF, LR, HW, CLG, NLS, DI, EH

**Formal analysis** - KKT, KEH

**Investigation** - IA, KLW, INO, AOO, OI, SB, CR, IHL, AF, MAKH, ØS, AS, NR, DMC, AS, FDM, GL, GG, FS, MLF

**Resources** - MC, HT, DS, GBB, FP, JC, AS, NR, AWJJ, MMCL, AS, GL, GG, FS, MLF, FK, NVK, NVT, LTH, NTH, MHP, BW, JV, EF, CR, AC, AS, NHV, VTTD, LTL, DMA, CY, SD, SC, JC

**Software** - KKT, CY

**Validation** - AA, NM, AL, JA, BW, FL, IB, JRI, AFL, JD, SRP, MHL, GS, AM, TMC, RC, VFL, MH, BN, JS, SR, RC, JR, TG, GMR, KY, MS, SK, YS, WH, NK, PNAH, BMF, DLP, LGCP, JNR, ASSM, EMMAB, DMC, AS, FDM, GL, GG, FS, MLF, AW, CB, MN, ND, FP, AC, AC, KLR, GN, VS, NLS, SL, NAF, OP, AE, HMBS, DI, IB, HS, LT, MZ

**Writing - original draft** - KKT, KEH

**Writing - review & editing** - All authors read and approved the final manuscript.

## ACKNOWLEDGEMENTS

The KlebNET-GSP AMR Genotype-Phenotype Group would like to acknowledge the Kp-T7 study, the Kp-MDR study, the Norwegian Study Group on *Klebsiella pneumoniae*, the Kp-NORM study, the NORKAB study, the JPI-AMR consortium SpARK, the NIHR Global Health Research Unit Genomic Surveillance of Antimicrobial Resistance, the ‘Controlling Superbugs’ flagship study, the Victorian Carbapenemase-Producing *Enterobacterales* (CPE) program, the KASPAH project, Vietnam ICU WGS study, the REDUCEAMU project, the MBIRA study, AGAR GnSOP, and the Molecular Epidemiology of carbapenem and colistin resistance and extended-spectrum beta-lactamase-producing *Enterobacteriaceae* among sepsis patients in Ethiopia project, the Surveillance and Characterization of Hypervirulent and Antibiotic-Resistant *Klebsiella* spp. isolates (SCHARKI) project in Germany, COMBACTE-CARE EURECA, EuSCAPE, Malawi Liverpool Wellcome Program (MLW), Queen Elizabeth Central Hospital (QECH), MERINO trial, iberoAMRica study, SARA consortium, and MedVetKlebs.

## ABBREVIATIONS

QRDR: quinolone resistance determining regions
PMQR: plasmid-mediated quinolone resistance
*Kp*SC: Klebsiella pneumoniae species complex
S: susceptible
I: intermediate / susceptible at increased exposure
R: resistant
CI: confidence interval
AMR: antimicrobial resistance
3GC: third-generation cephalosporin
WHO: World Health Organization
AST: antimicrobial susceptibility testing
UTI: urinary tract infection
GLASS: Global AMR Surveillance System
TMQR: transferable mechanisms of quinolone resistance
WGS: whole-genome sequencing
MIC: minimum inhibitory concentration
*h*^2^: heritability
*Kvv*: Klebsiella variicola subsp. variicola
*Kqs*: Klebsiella quasipneumoniae subsp. similipneumoniae
*Kqq*: Klebsiella quasipneumoniae subsp. quasipneumoniae
ST: sequence type
ECOFF: epidemiological cut-off
WT: wildtype
NWT: non-wild type
OR: odds ratio
AIC: Akaike Information Criterion
ME: major error
VME: very major error
PPV: positive predictive value
LIN: Life Identification Number
GRAM: Global Research on Antimicrobial Resistance
ANOVA: analysis of variance

## SUPPLEMENTARY FIGURES

**Figure S1.**
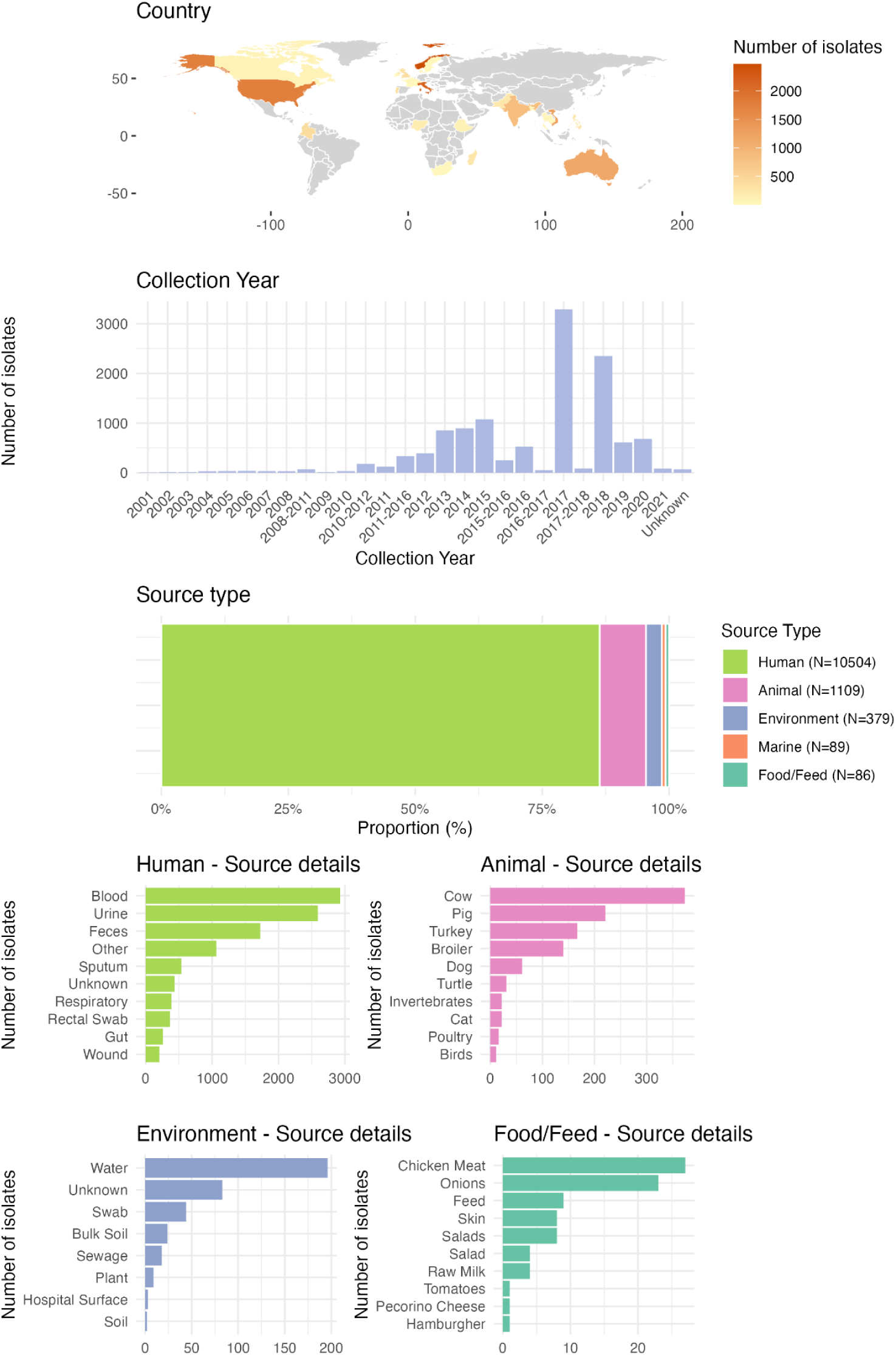
Global and diverse collection of *Kp*SC isolates with ciprofloxacin susceptibility tests (N=12,167) from the KlebNET-GSP AMR Genotype-Phenotype Group. The isolates were collected from 27 countries between 2001 and 2021 and sampled from predominantly human sources, but also include animals, environment, and food / feed sources.

**Figure S2.**
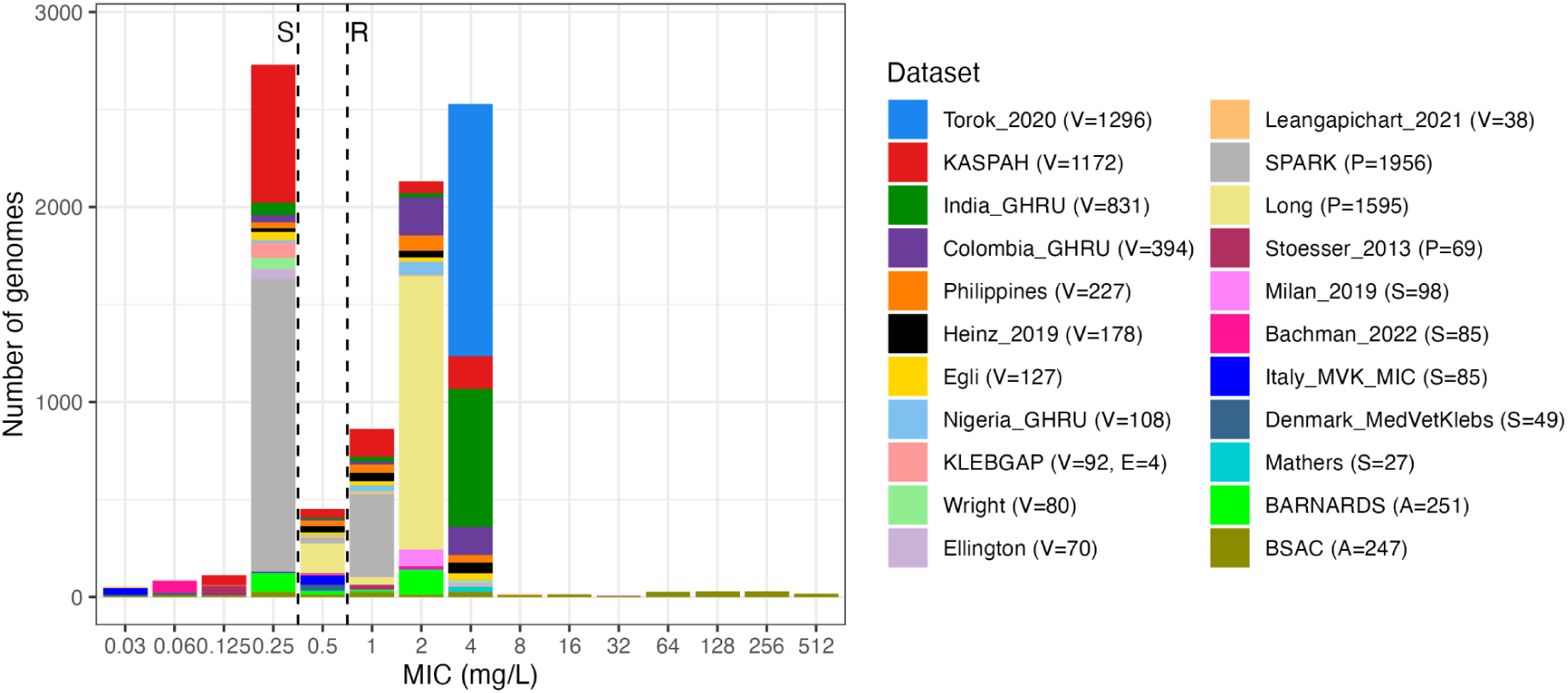
Minimum inhibitory concentration. (**MIC) distribution for discovery datasets.** The number of isolates tested using each MIC method (Vitek (V), E-test (E), Phoenix (P), Sensititre (S), and agar dilution (A)) is indicated within the bracket. All datasets have lower bounds (</≤) and upper bounds (>/≥) with the exception of the BSAC (A=251) and KLEBGAP (E=4) datasets. Vertical lines indicate the EUCAST/CLSI S breakpoint (≤0.25 mg/L) and the EUCAST/CLSI R breakpoint (>0.5 mg/L).

**Figure S3.**
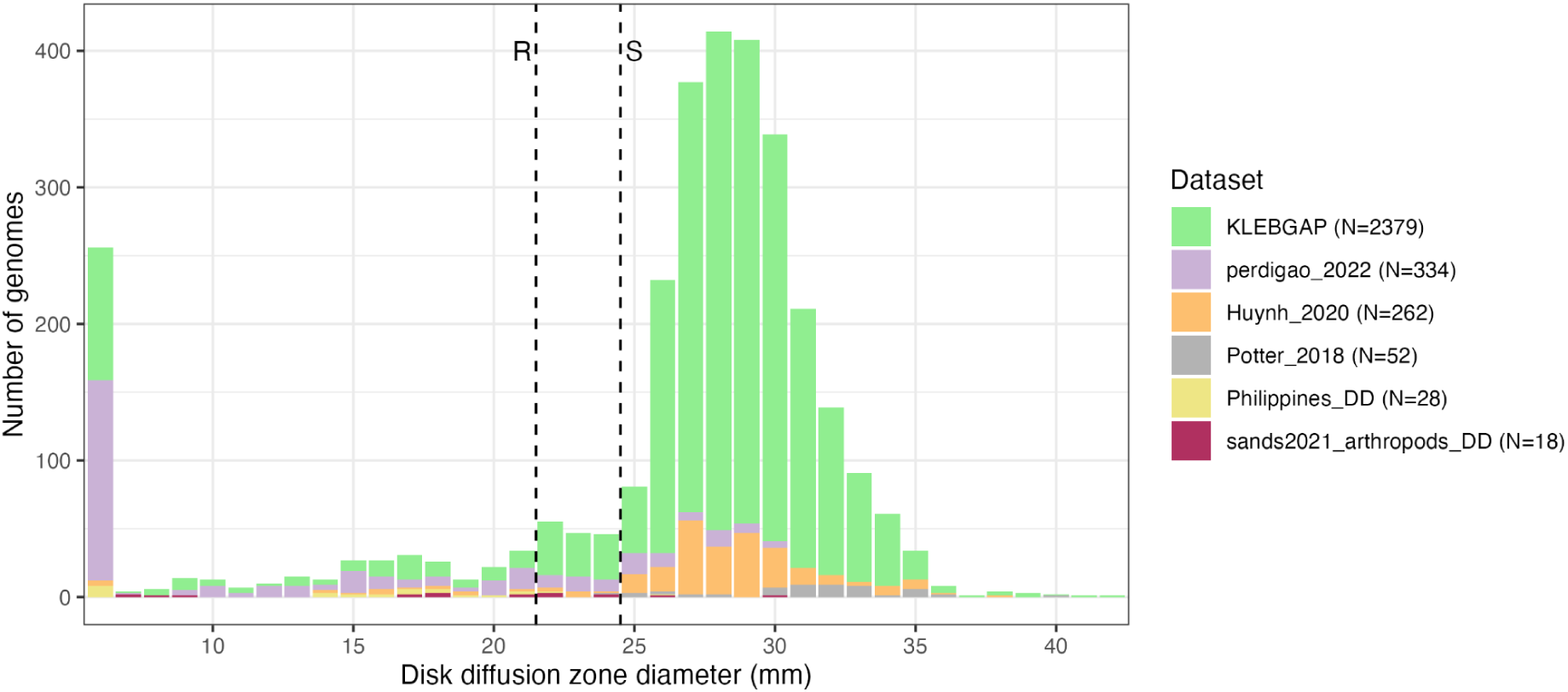
Disk diffusion distributions for discovery datasets. The number of isolates tested with disk diffusion are indicated in brackets. Vertical lines indicate the EUCAST/CLSI R breakpoint (<22 mm) and the EUCAST S breakpoint (≥25 mm). Note the CLSI breakpoint is ≥26 mm, and measurements generated using CLSI protocol (Philippines, N=28 and Potter 2018, N=52) were interpreted with this breakpoint.

**Figure S4.**
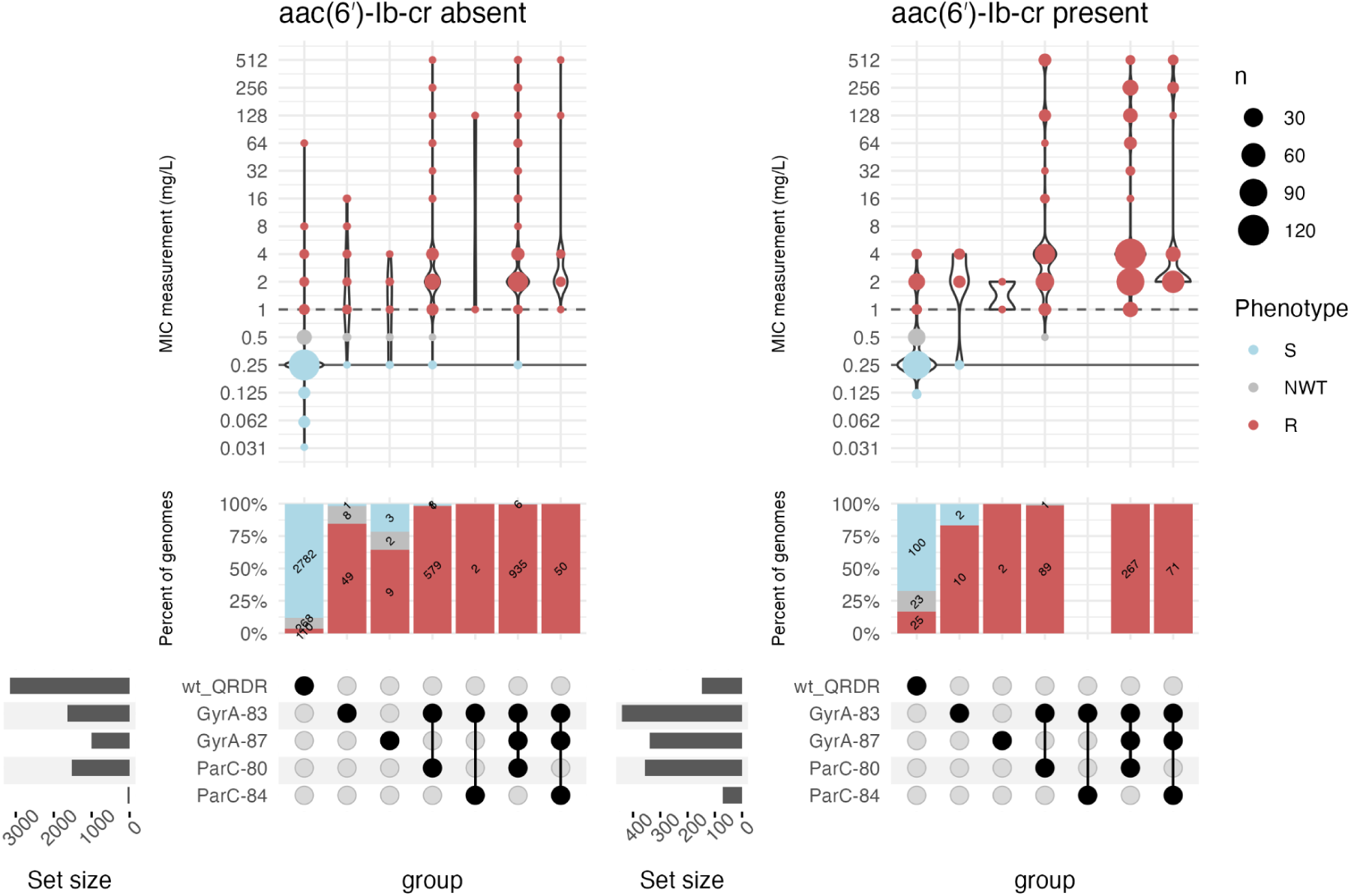
MIC distribution for QRDR mutations, in the absence of PMQR genes. Filled circles in the upset plot indicate the genotype as indicated in the row label. wt_QRDR indicates genomes with wildtype QRDRs.

**Figure S5.**
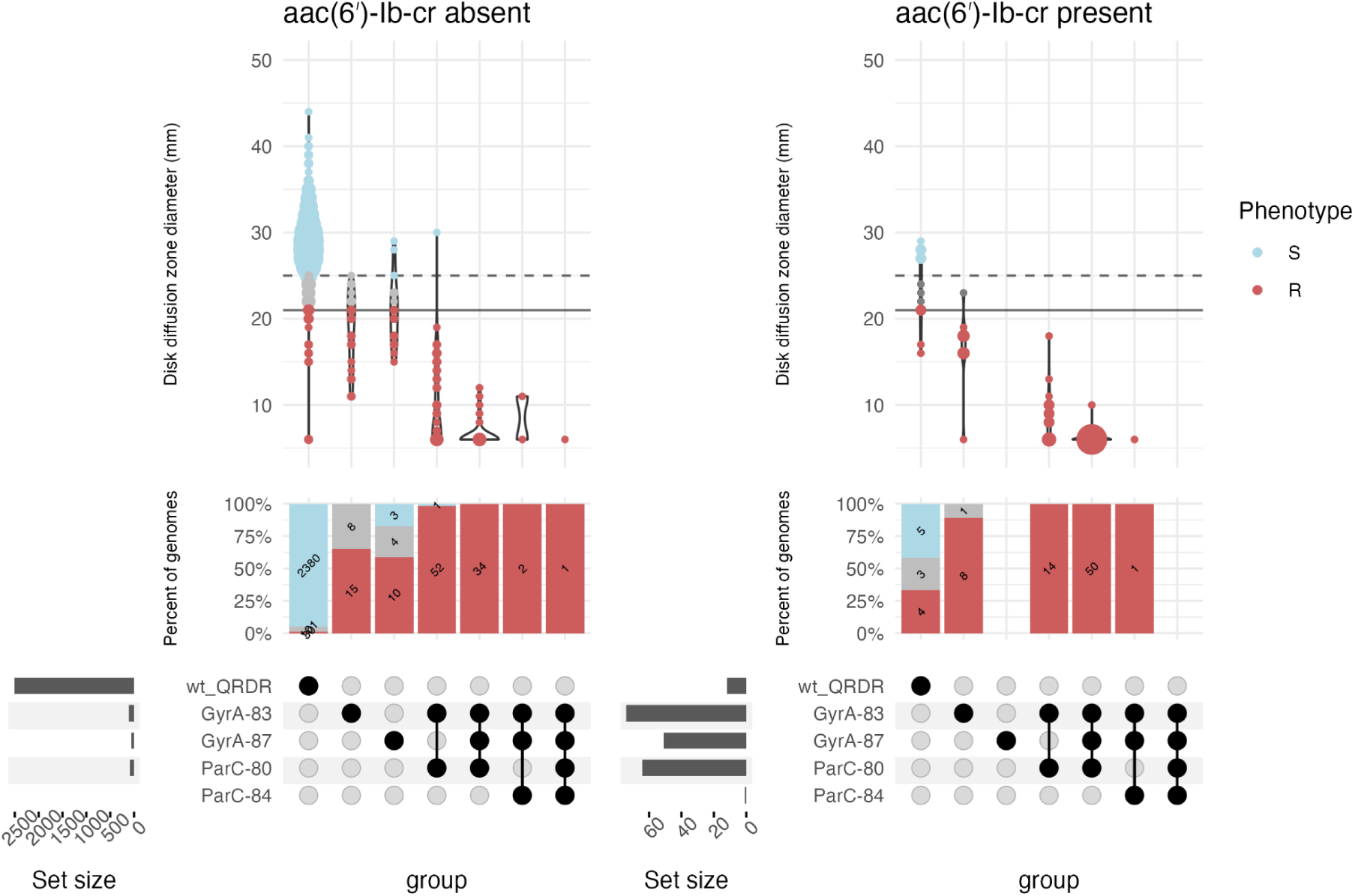
Disk diffusion zone diameter distribution for QRDR mutations, in the absence of PMQR genes. Filled circles in the upset plot indicate the genotype as indicated in the row label. wt_QRDR indicates genomes with wildtype QRDRs.

**Figure S6.**
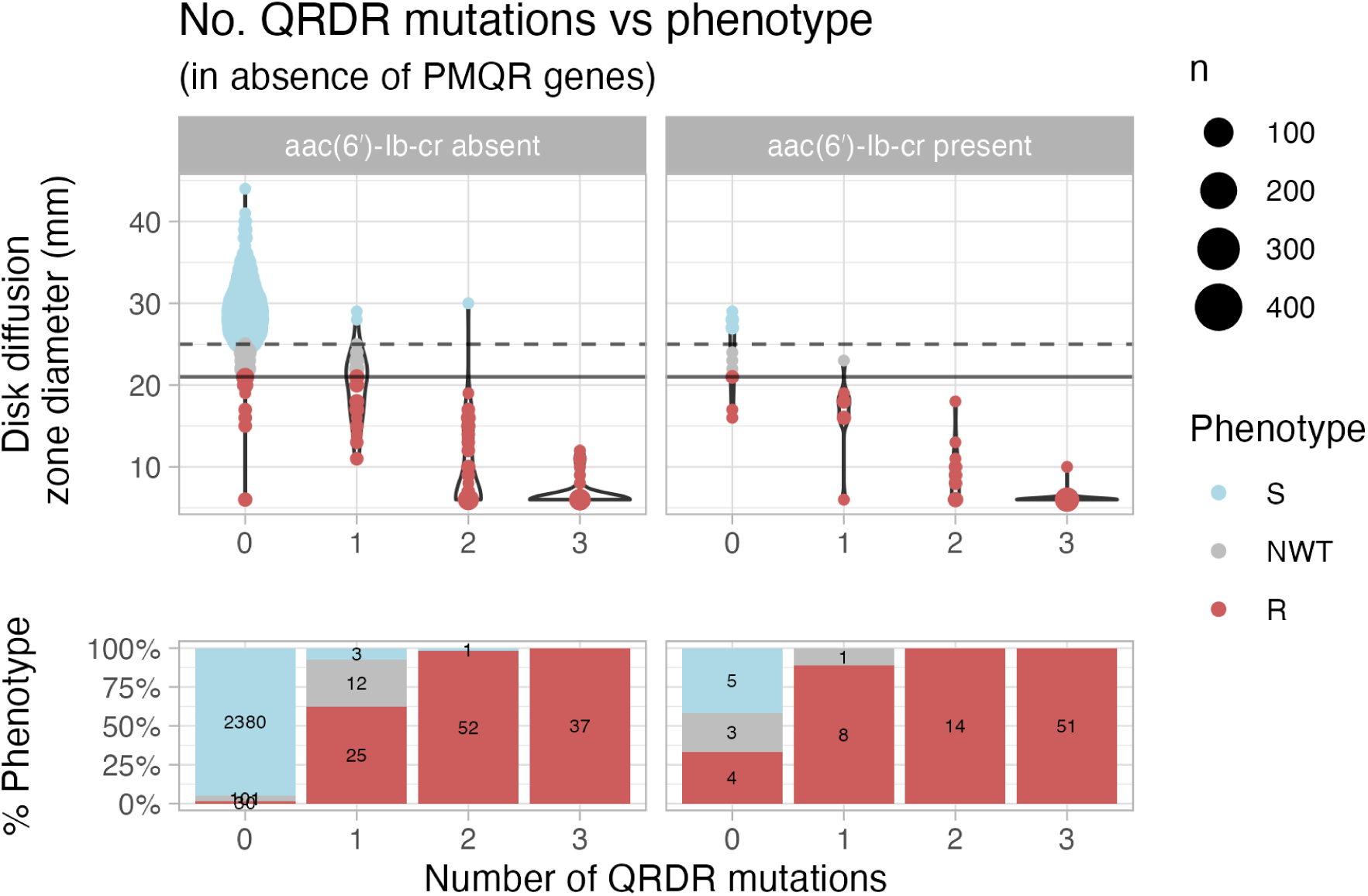
Number of QRDR mutations vs phenotype, for isolates with disk diffusion measurements. Filled circles in the upset plot indicate the genotype as indicated in the row label.

**Figure S7.**
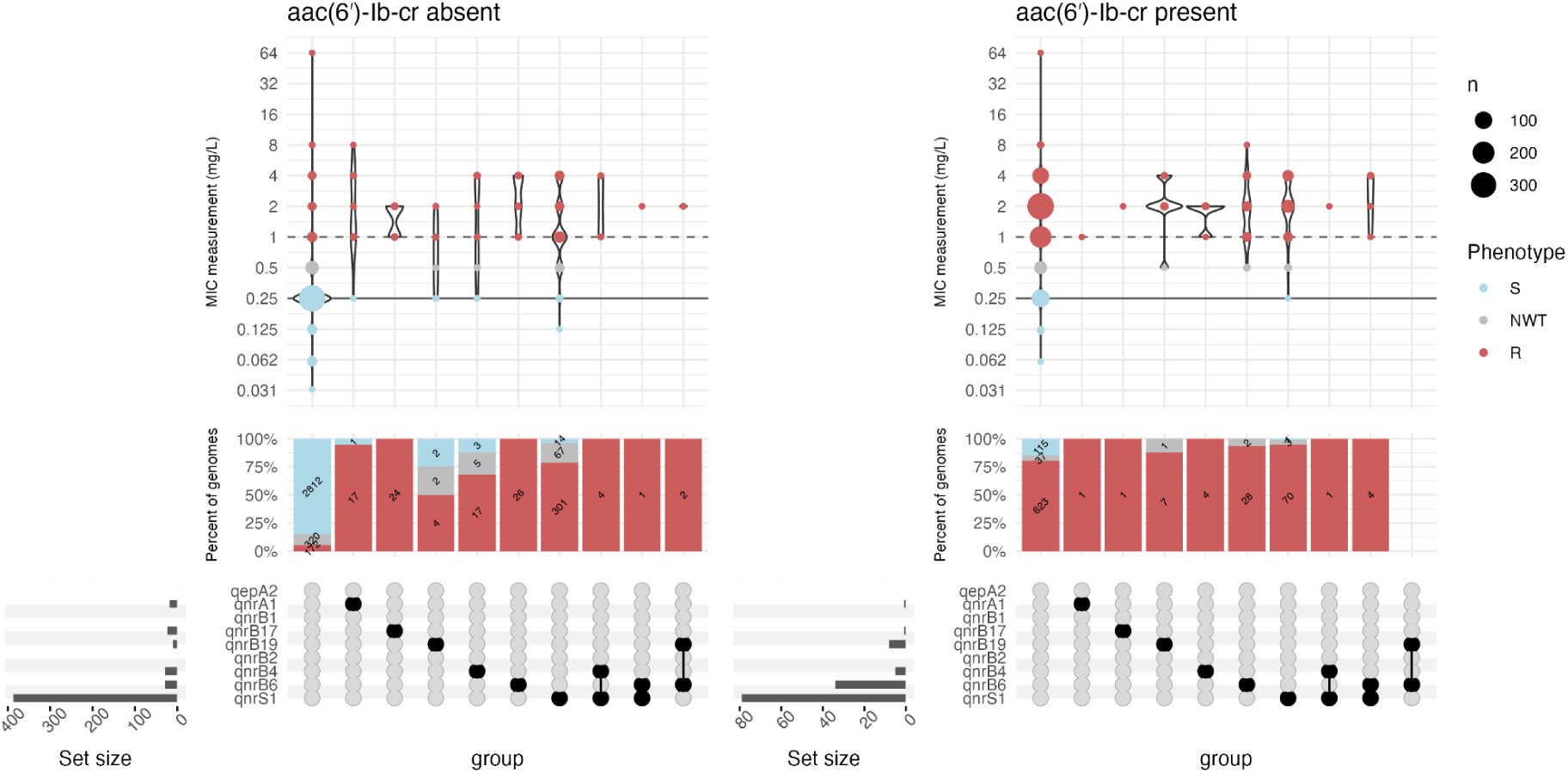
MIC distribution for PMQR gene combinations, in the absence of QRDR mutations. Filled circles in the upset plot indicate the genotype as indicated in the row label.

**Figure S8.**
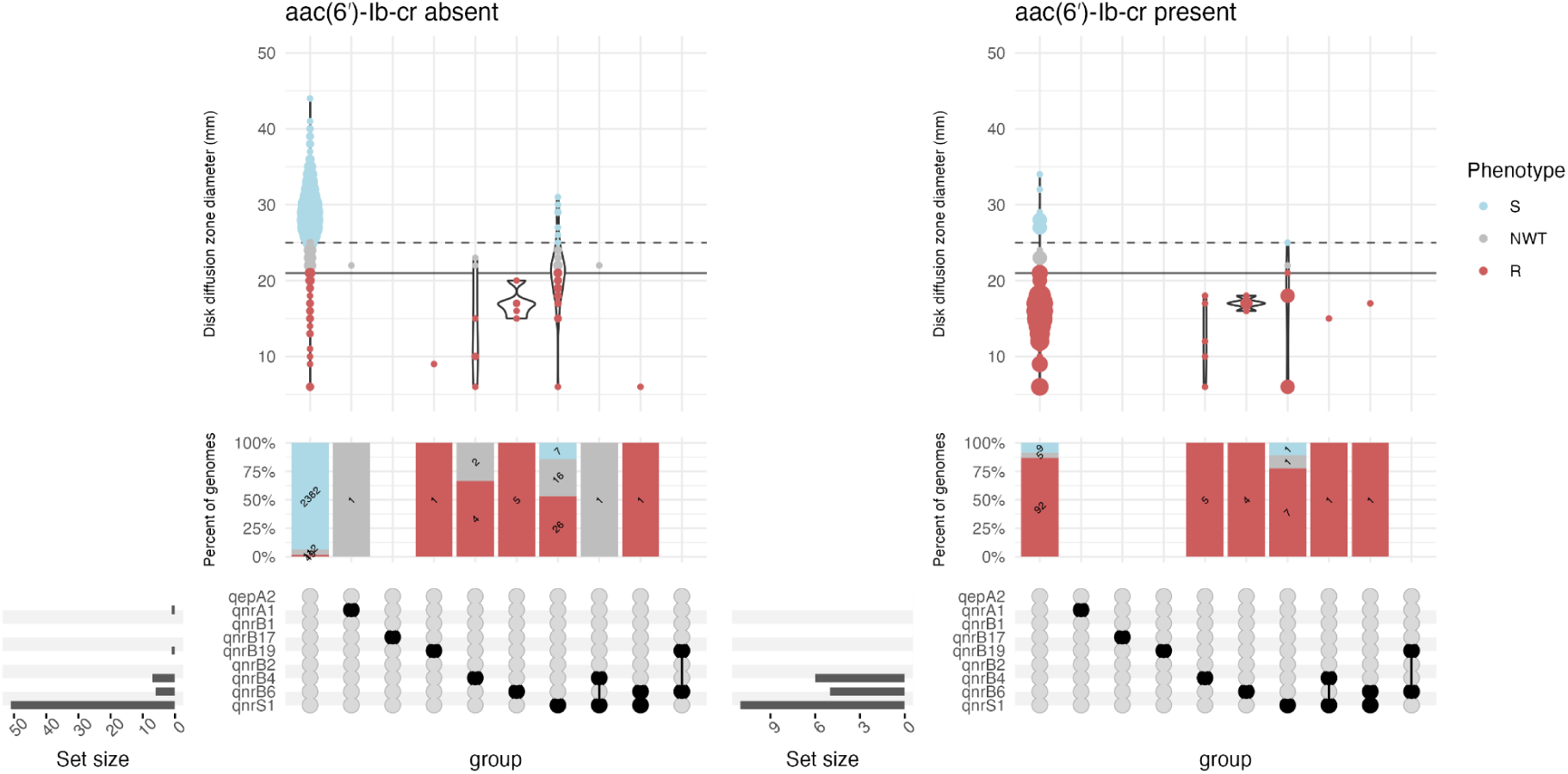
**Disk diffusion zone diameter distribution for PMQR gene combinations, in the absence of QRDR mutations.**

**Figure S9.**
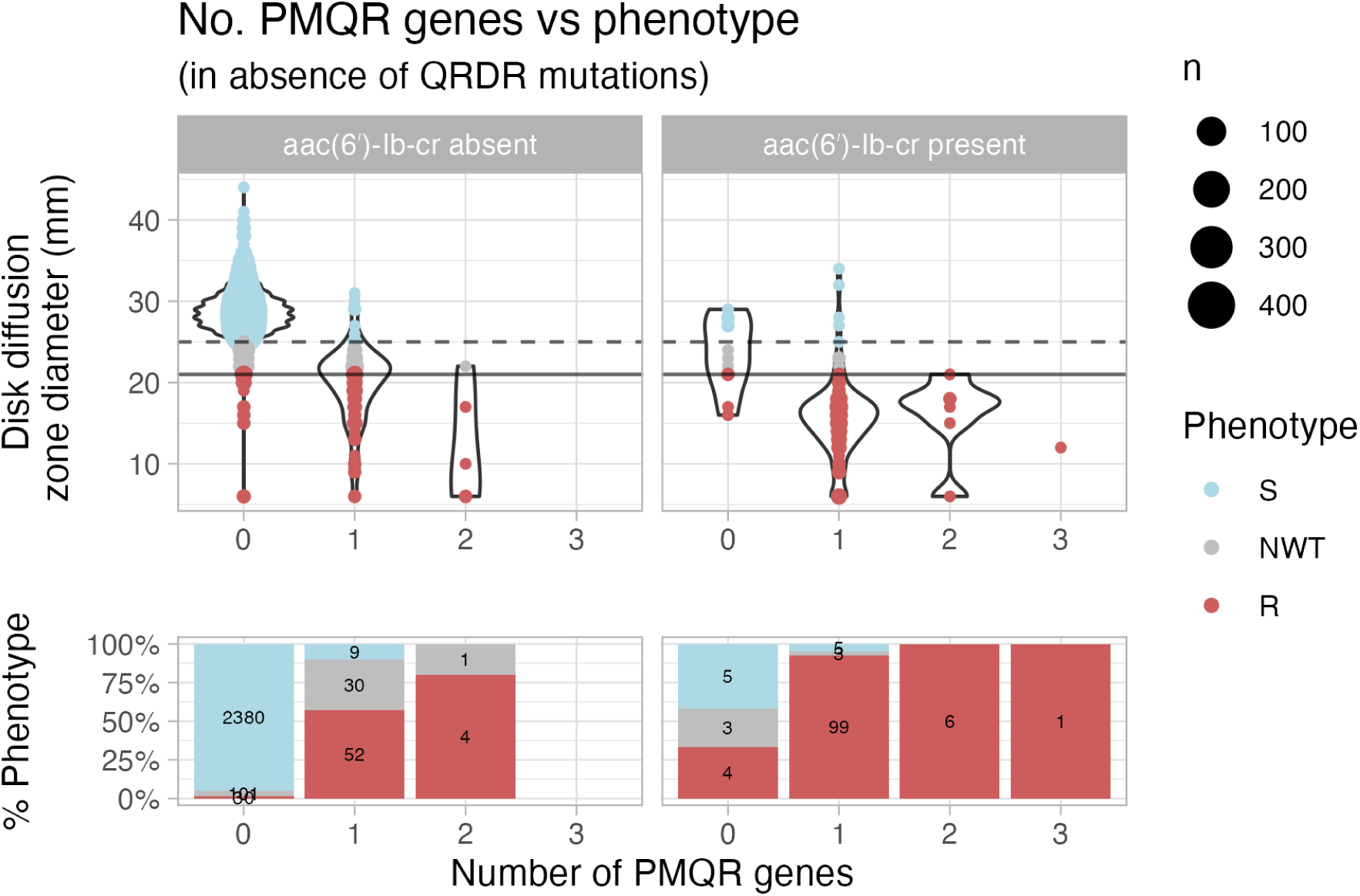
Number of *qnr*/*qep* genes vs phenotype, for isolates with disk diffusion measurements.

**Figure S10.**
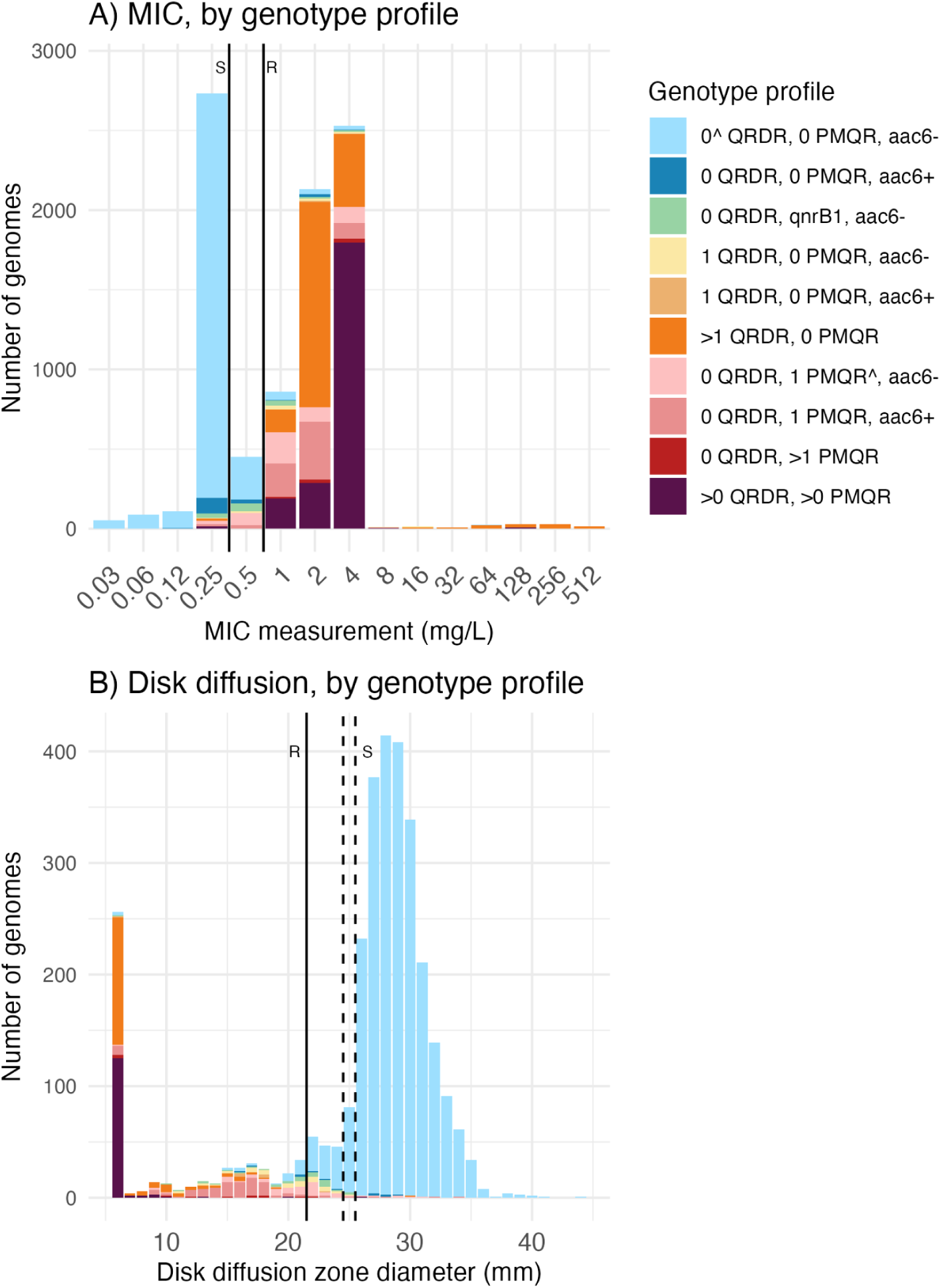
Phenotype distributions stratified by genotype profiles. (A) MIC measurement (mg/L) and (B) disk diffusion zone diameter (mm) distributions coloured according to the genotype profiles assigned by the rules-based classifier. All MIC datasets have lower bounds (</≤) and upper bounds (>/≥) with the exception of two datasets (totalling N=255 isolates, see **Figure S1, Table S3** for more detail). The vertical solid line indicates the EUCAST/CLSI R breakpoint (<22mm) and the vertical dotted lines indicate the EUCAST S breakpoint (≥25 mm) and CLSI S breakpoint (≥26 mm).

**Figure S11.**
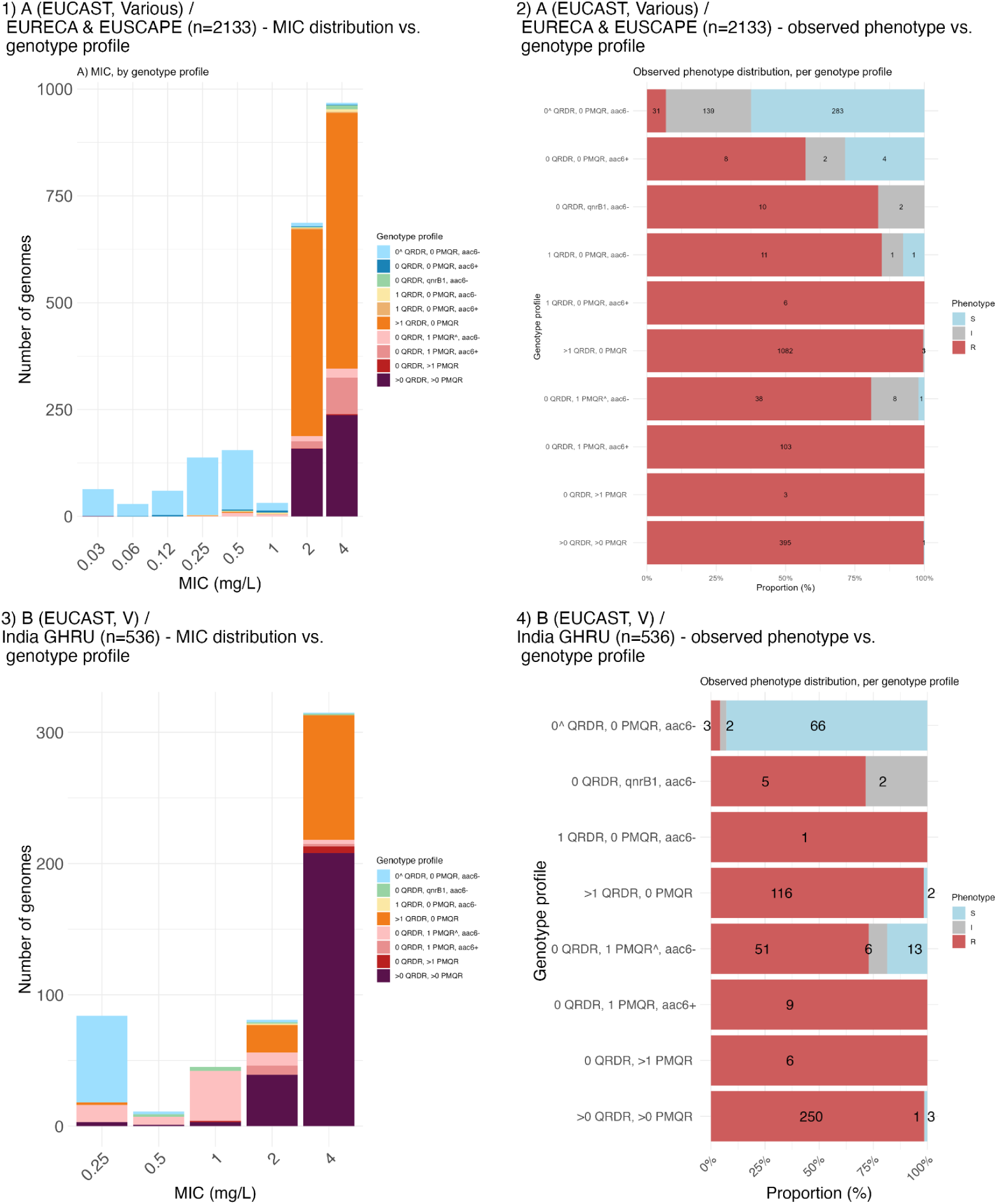
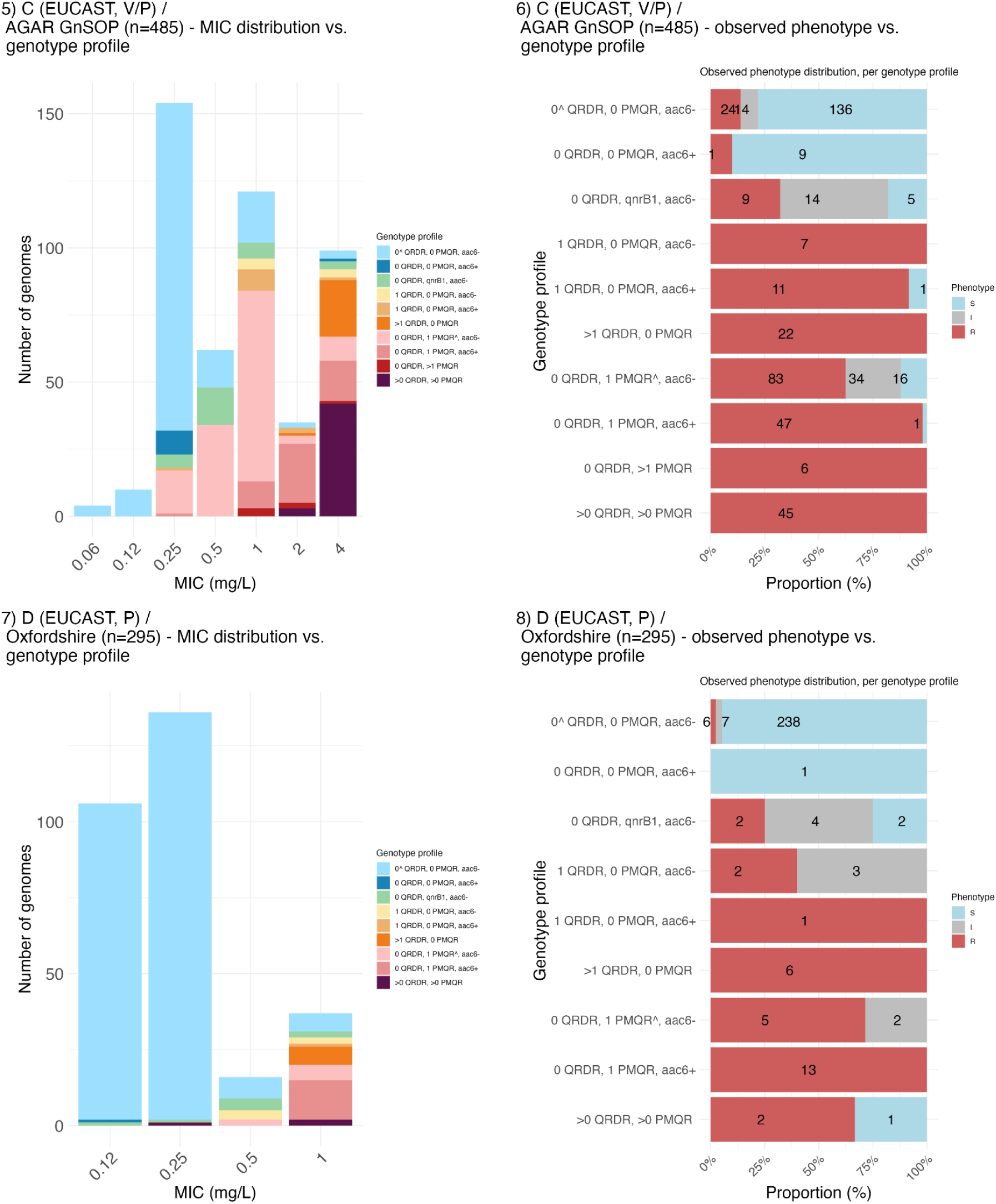
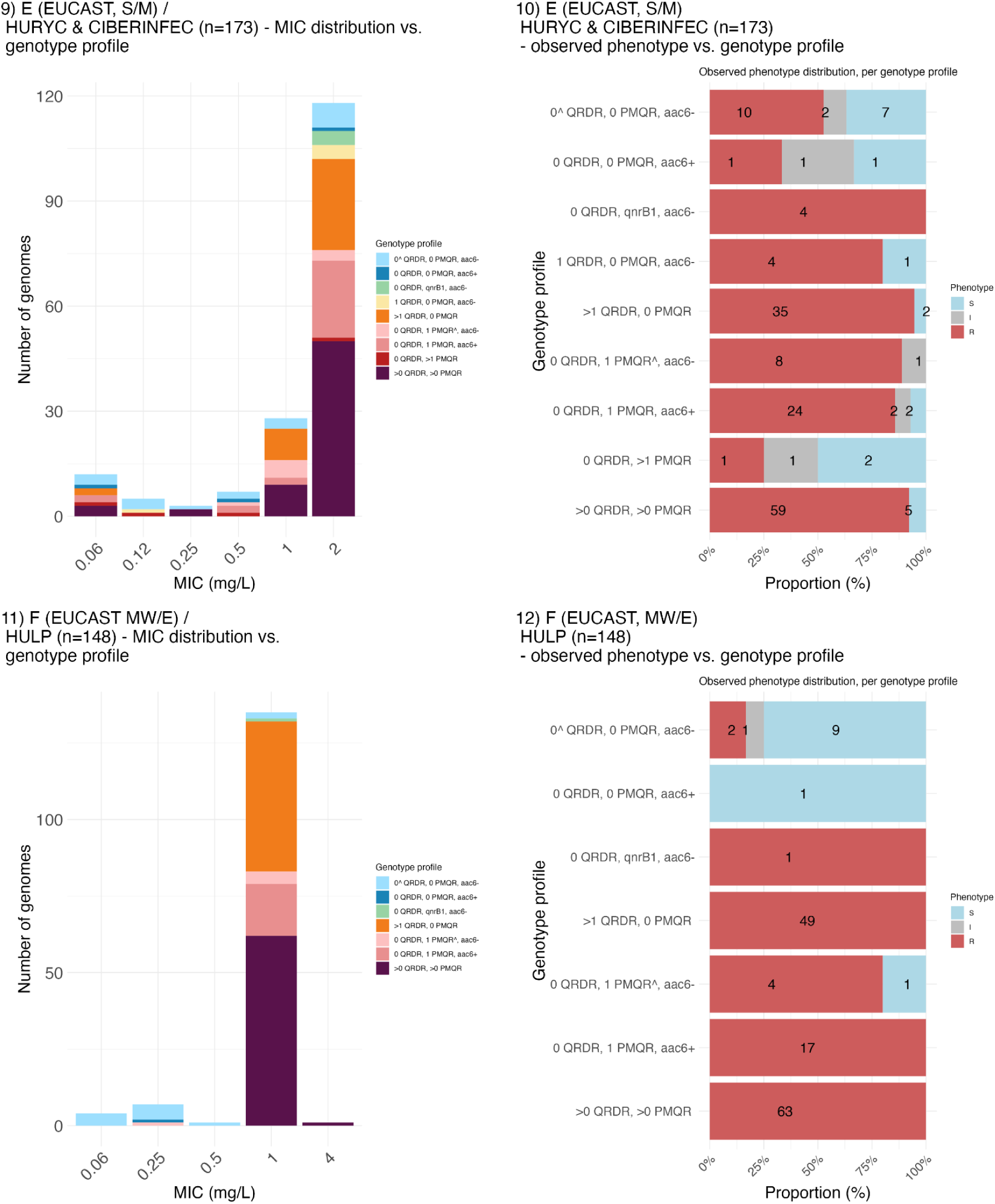
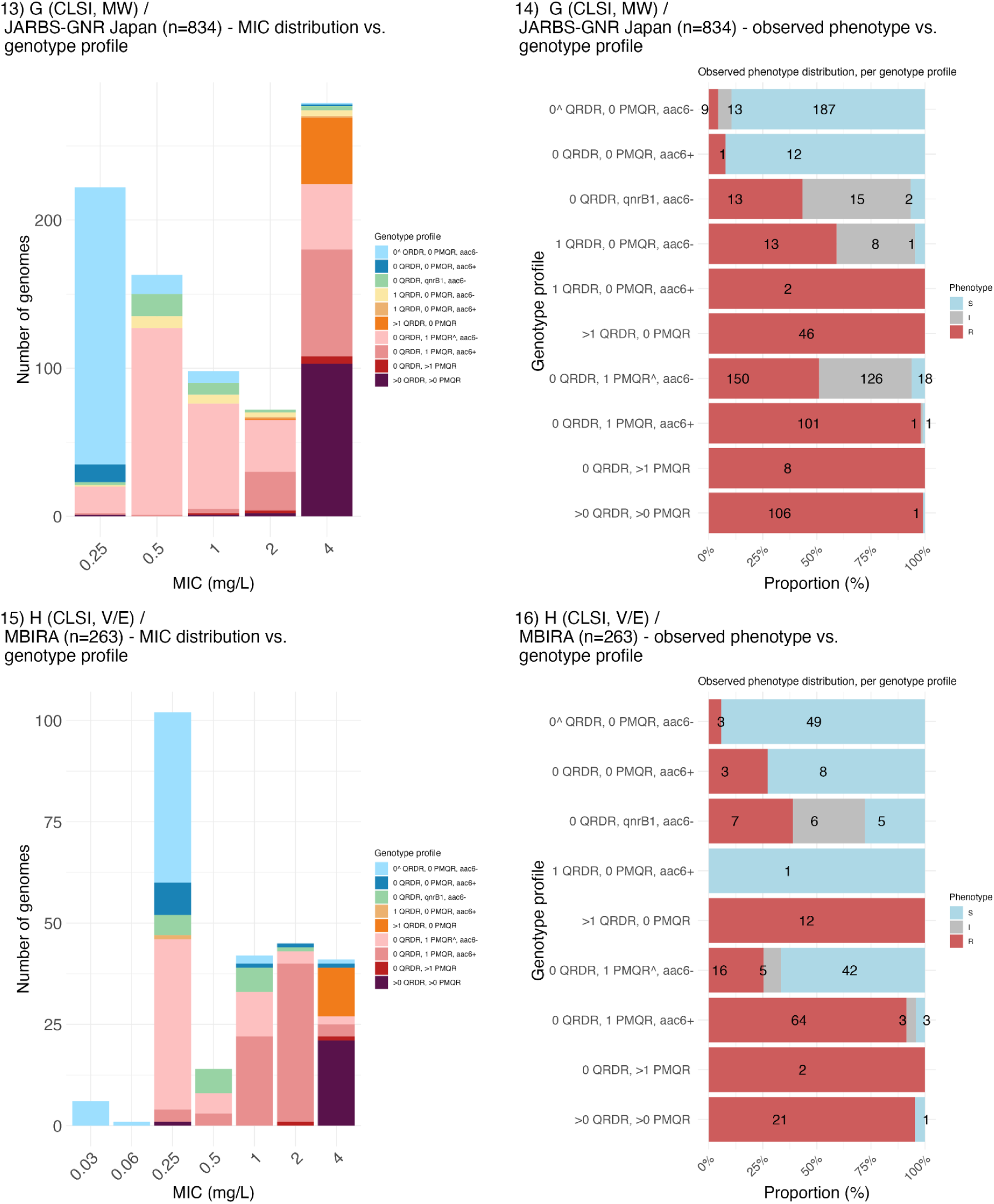
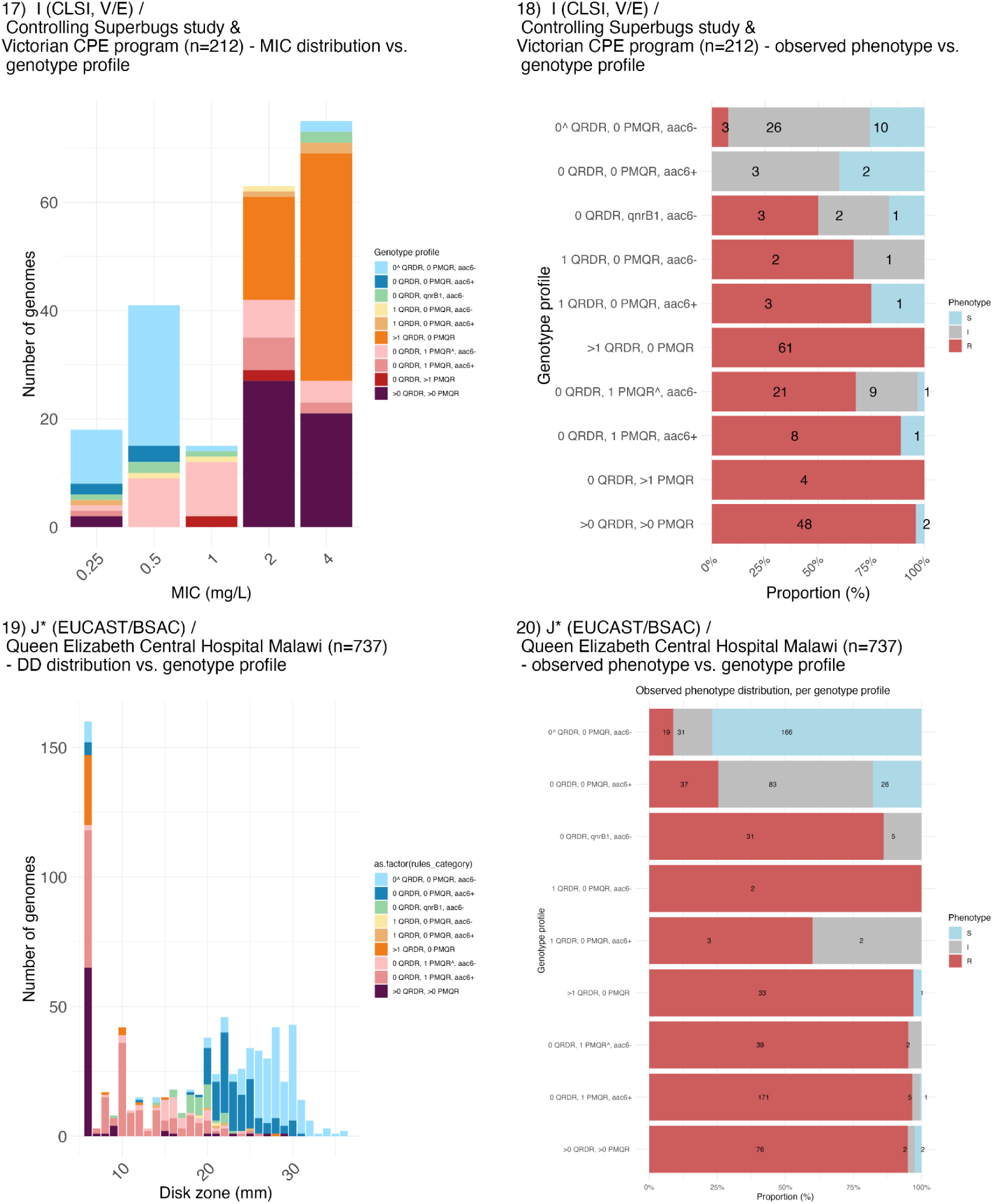
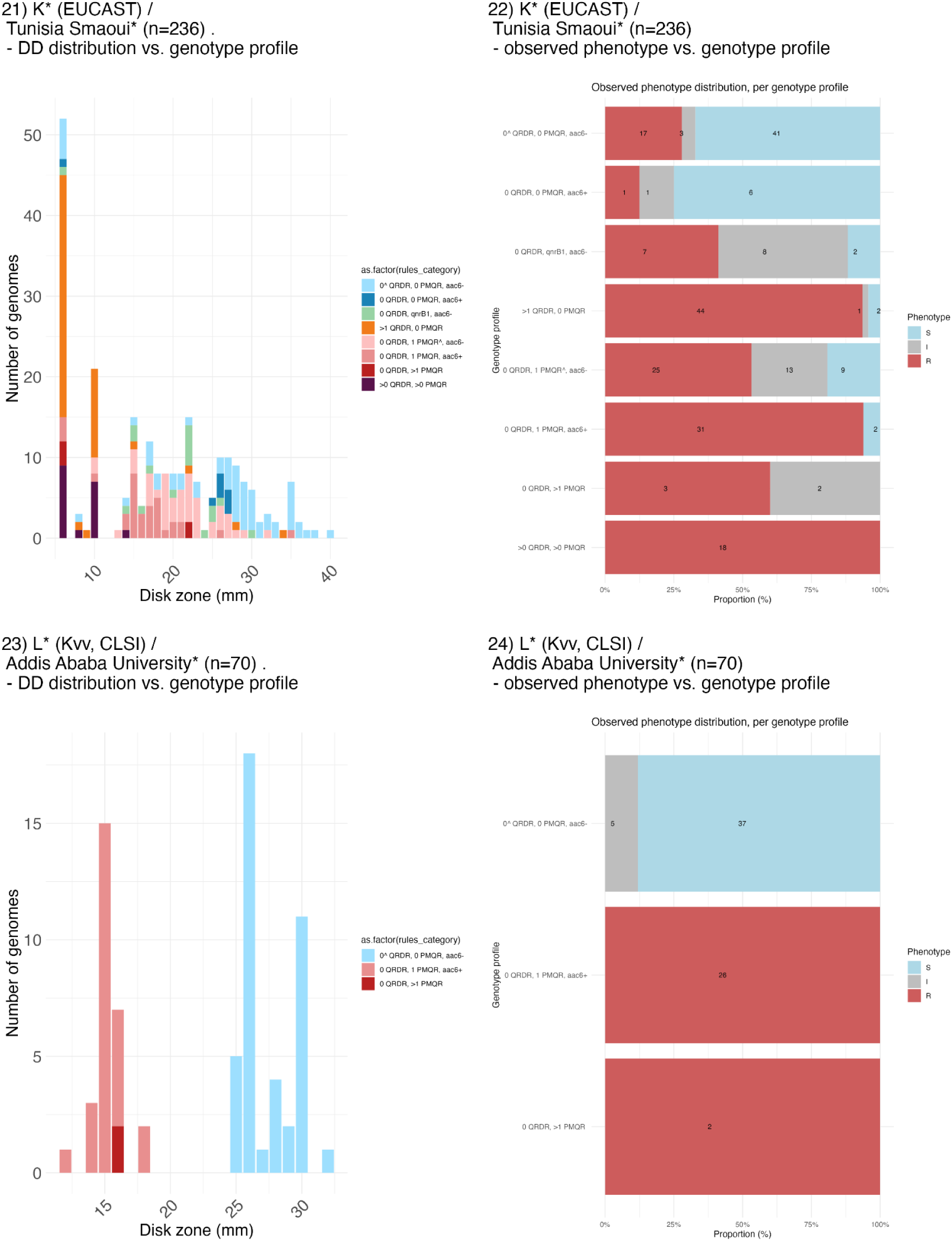
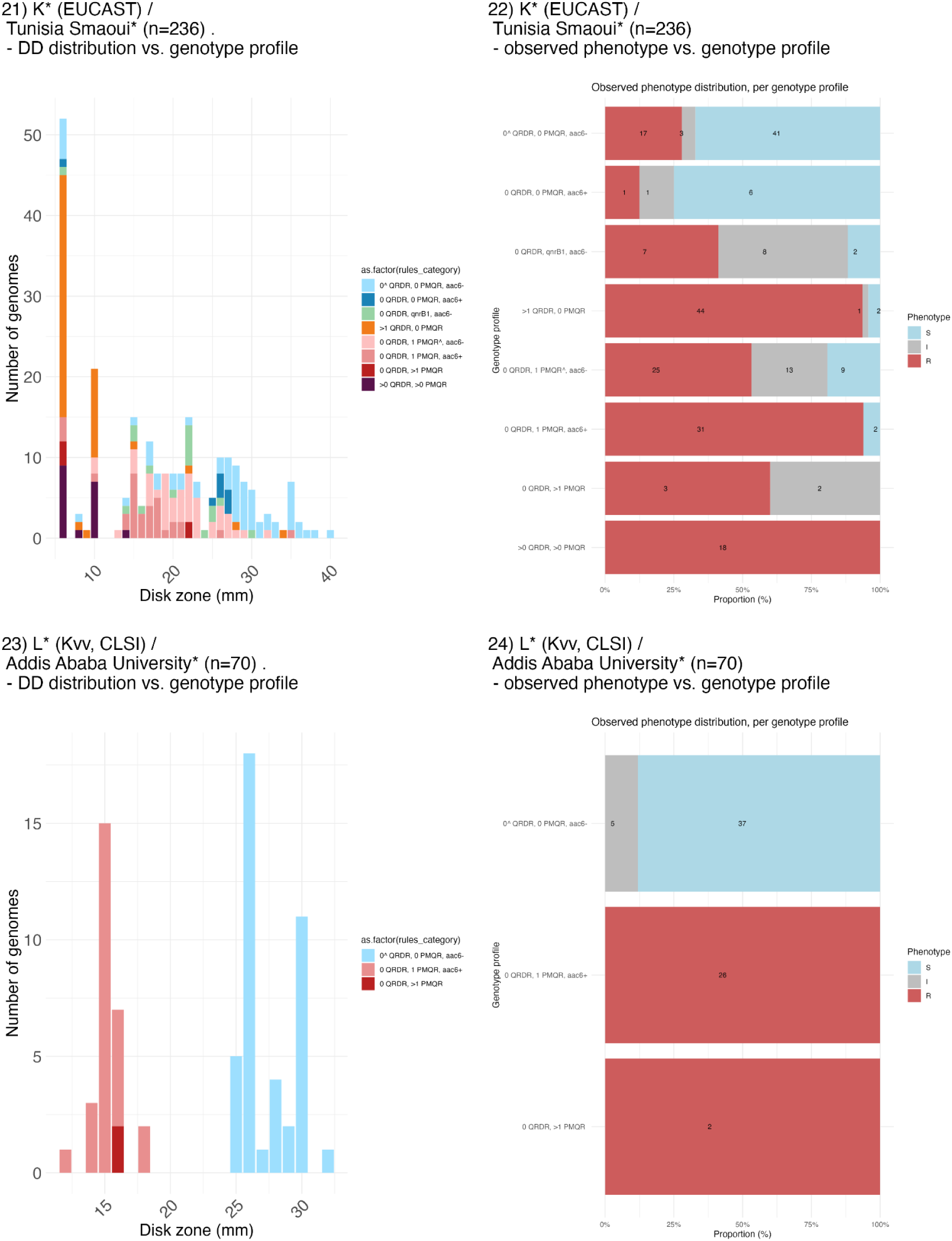
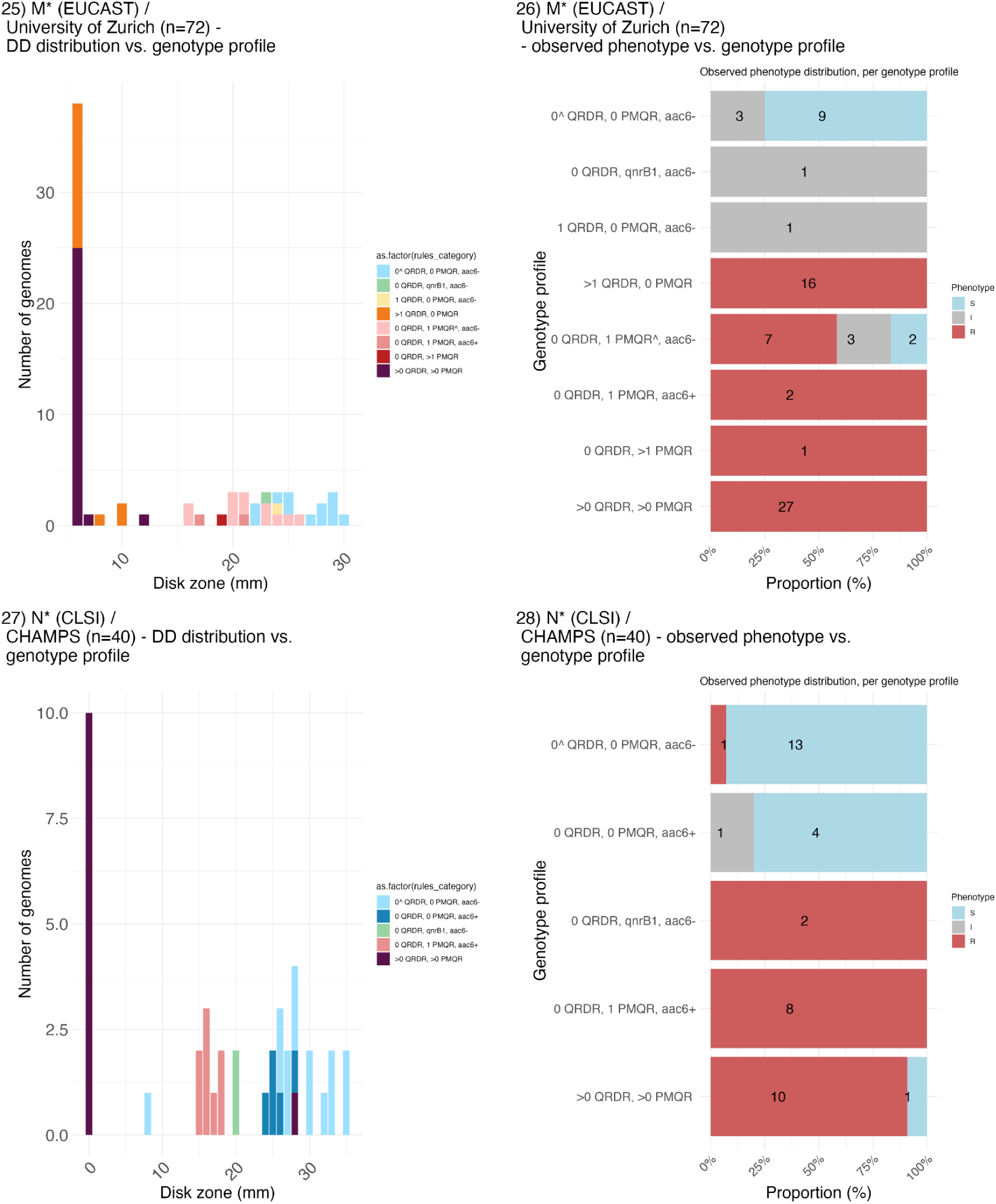
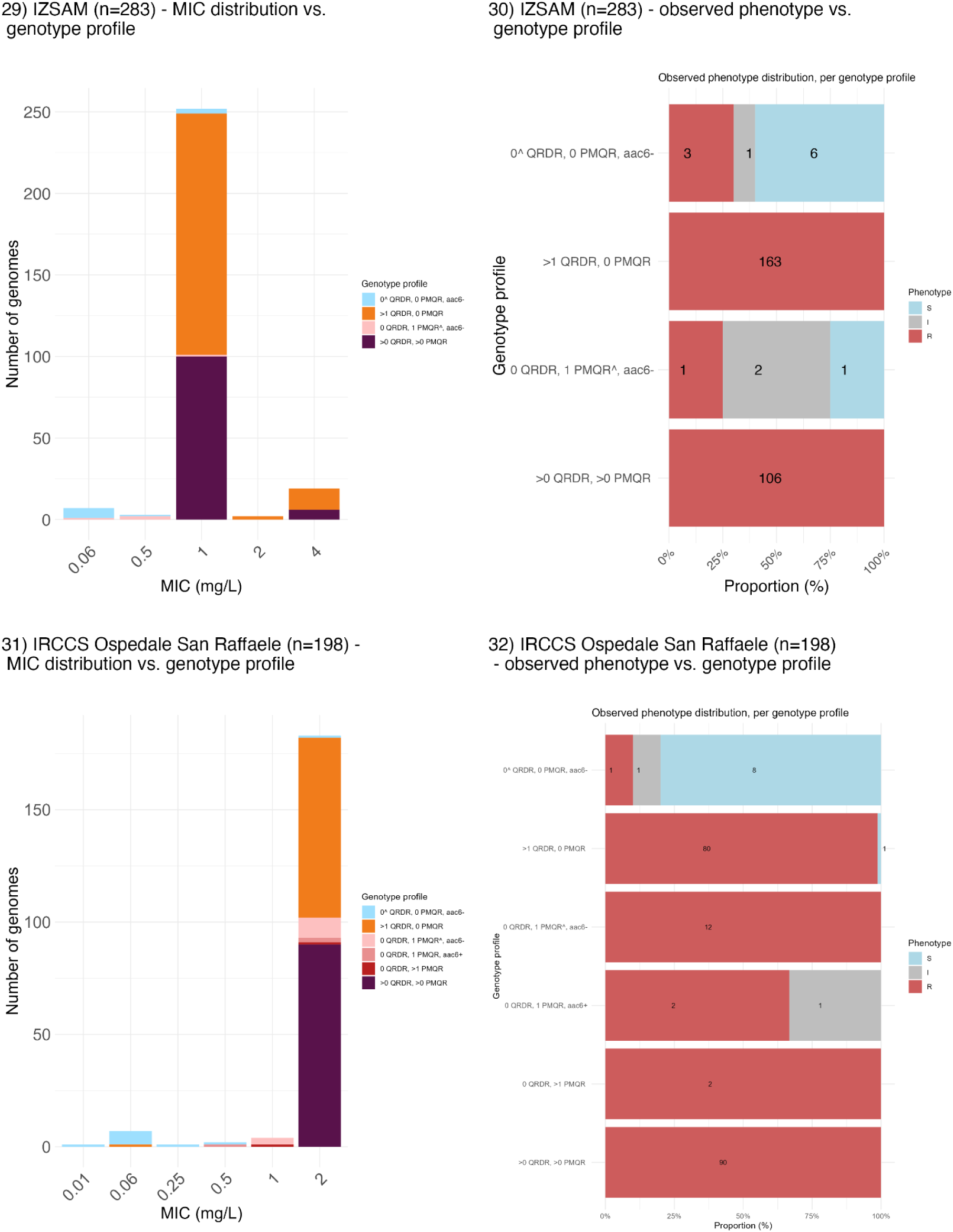
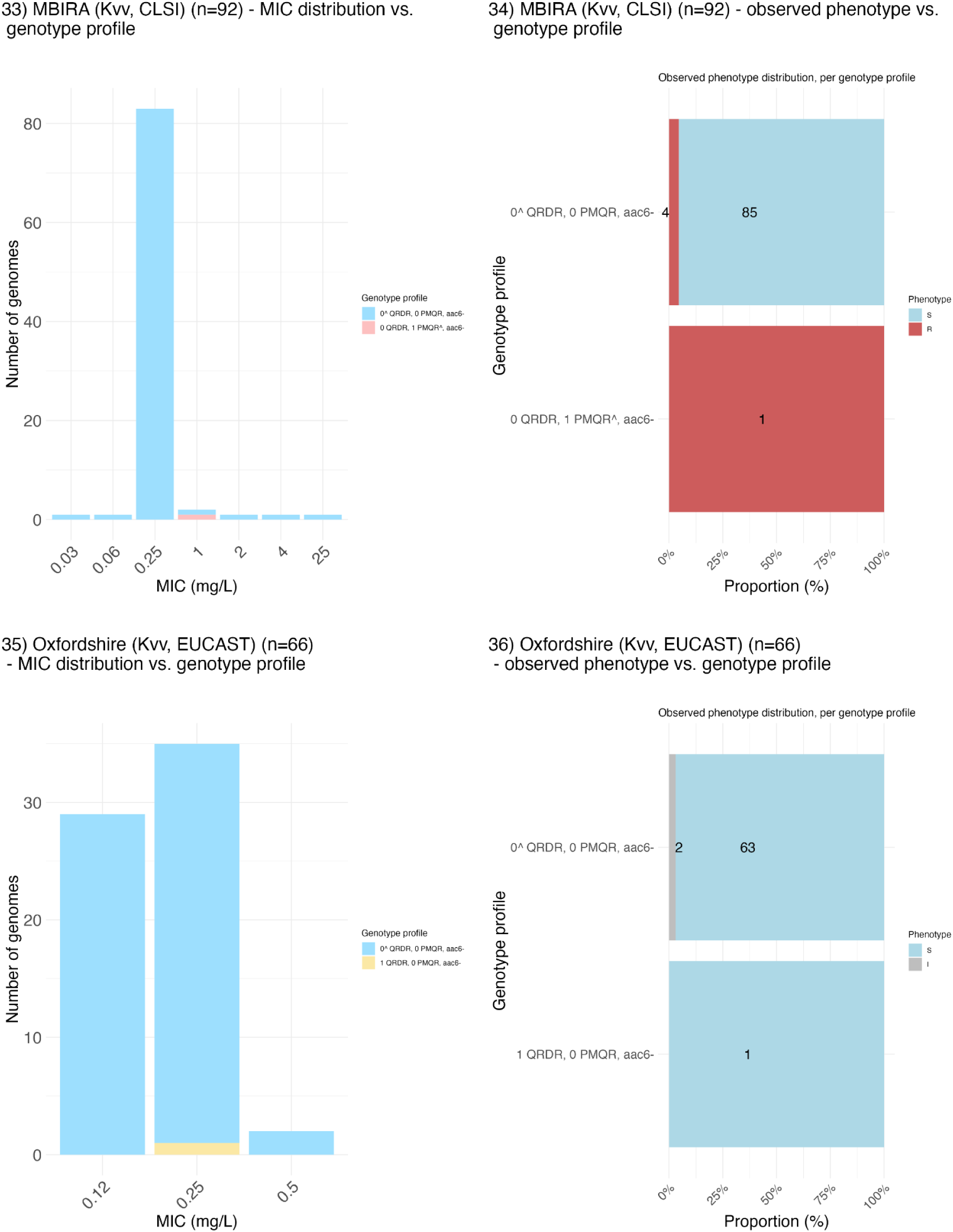
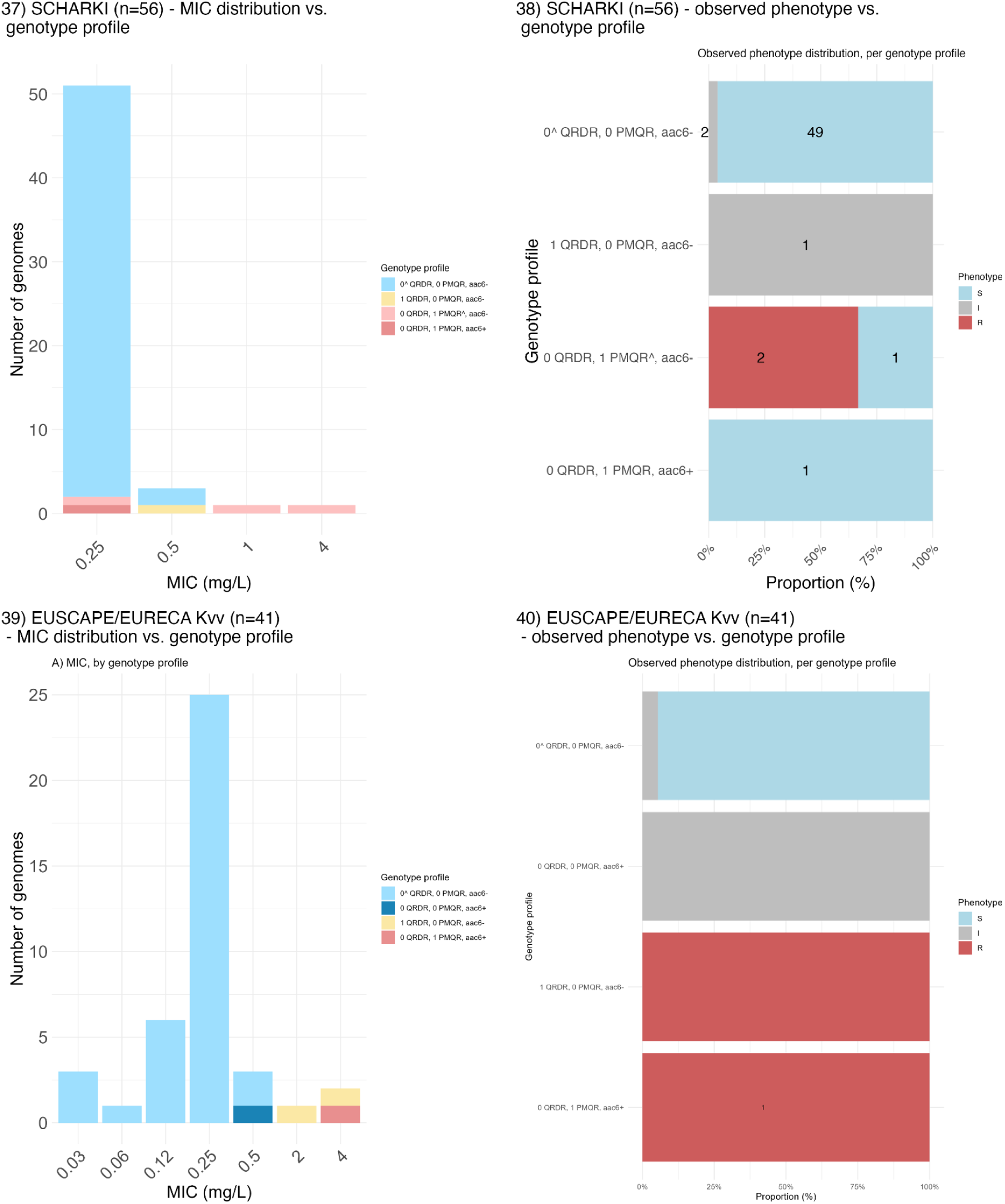
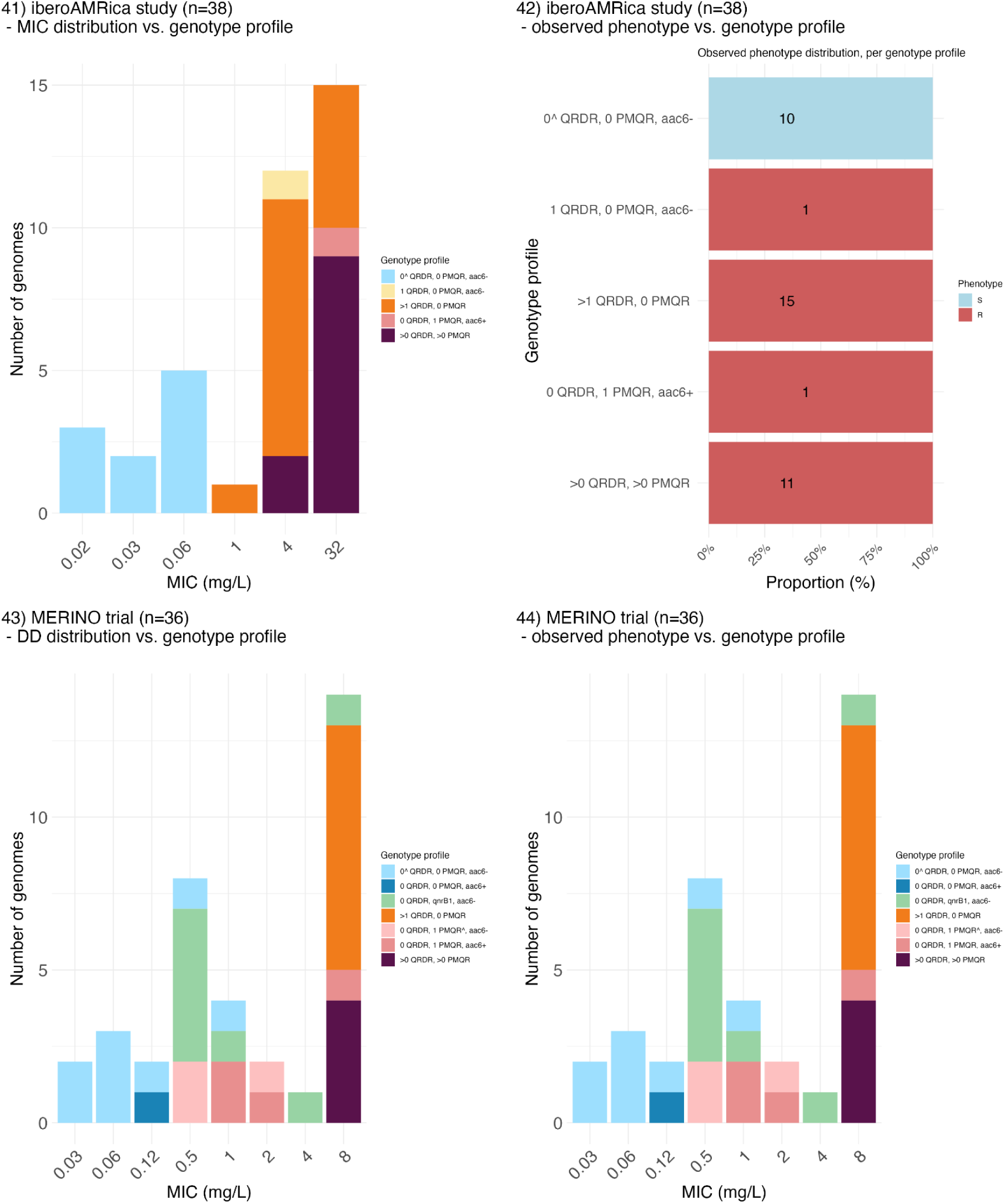
Genotype profiles of external validation datasets. (Odd numbered panels, e.g., 1, 3, 5, etc.) MIC or disk diffusion zone (DD) distributions coloured by genotype profile assigned by the rules-based classifier per dataset. (Even numbered panels, e.g., 2, 4, 6, etc.) Frequency of true, observed S/I/R phenotypes observed for genomes categorized into each genotype profile used in the rules-based classifier. Ciprofloxacin susceptibility phenotypes were measured using MIC or disk diffusion (indicated by *). For each dataset, the breakpoint guidelines used are indicated in the first bracket, followed by MIC method (Vitek (V), Phoenix (P), Sensititre (S), Microscan Siemens (M), Microscan WalkAway Beckman Coulter (MW), and E-test (E)). Dataset A includes data from laboratories across Europe and they used various MIC methods.

**Figure S12.**
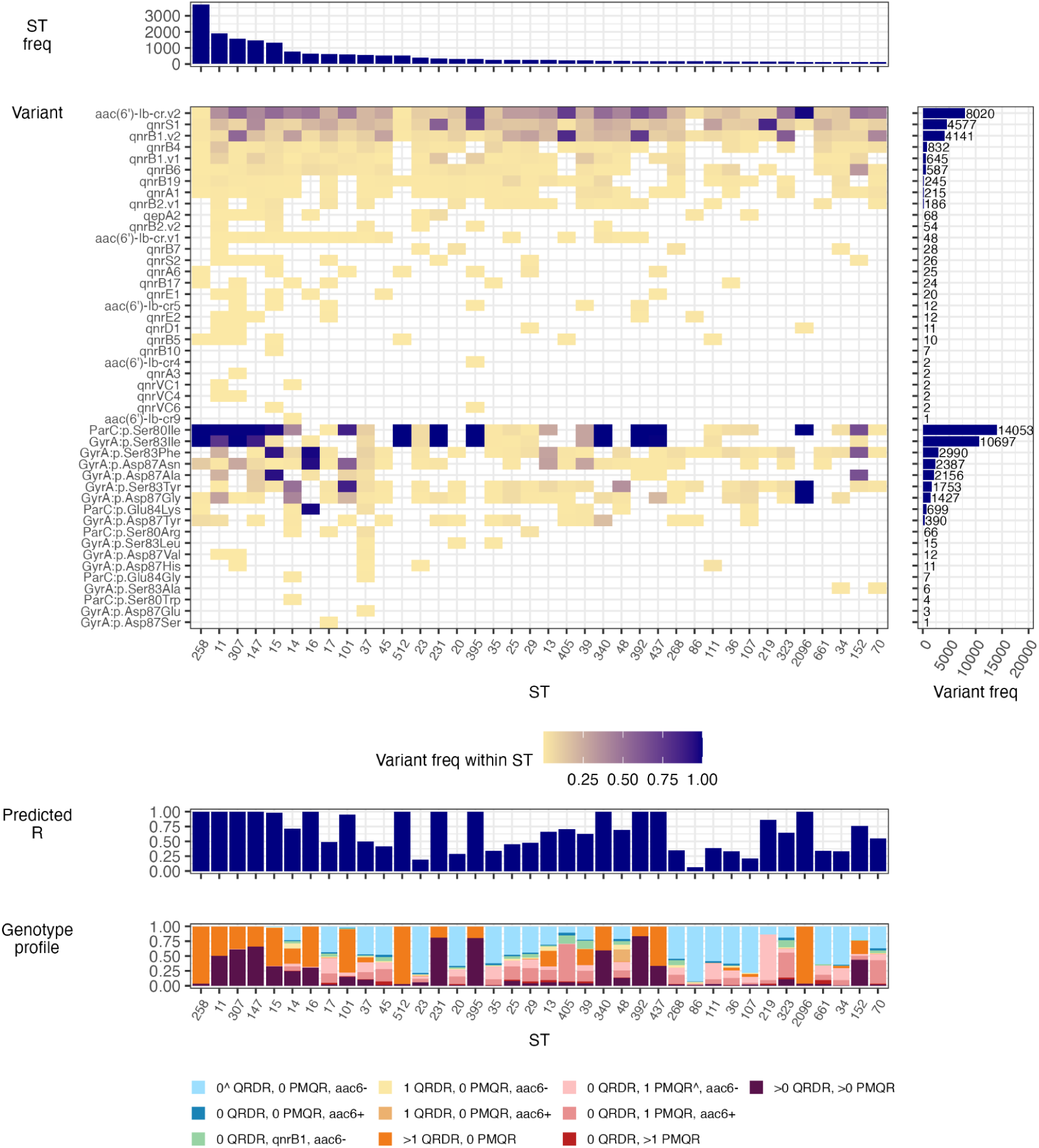
Ciprofloxacin resistance determinants and predicted resistance in public genome data for *K. pneumoniae*. Data were downloaded from Pathogenwatch, which assembled all read sets available in ENA as of March 29, 2023 and used Kleborate v3.2.4 to identify resistance variants and 7-locus sequence types (STs), and predict ciprofloxacin resistance using our rules-based classifier. Details are shown for STs with n>100 genomes (n=38 STs, n=18,212 genomes). Variant frequency (freq) in the right-hand panel indicates the total number of genomes in which each determinant was identified (out of n=18,212 *K. pneumoniae* genomes). ‘Predicted R’ indicates the proportion of genomes predicted as resistant based on genotype profile, using the rules-based classifier.

**Figure S13.**
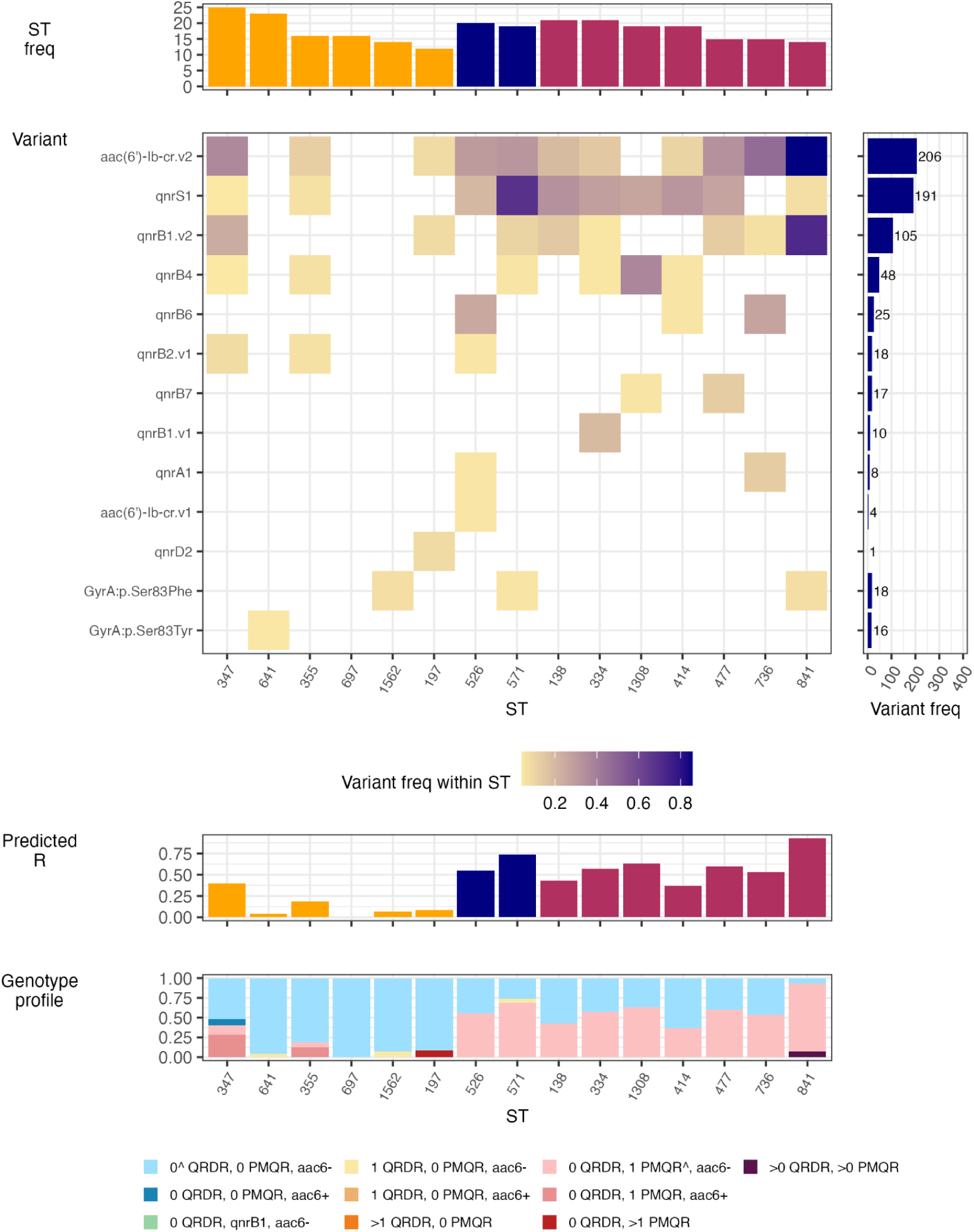
Ciprofloxacin resistance determinants and predicted resistance in public genome data for other members of the *K. pneumoniae* species complex. Data were downloaded from Pathogenwatch, which assembled all read sets available in ENA as of March 29, 2023 and used Kleborate v3.2.4 to identify resistance variants and 7-locus sequence types (STs), and predict ciprofloxacin resistance using our rules-based classifier. Details are shown for STs with N>10 genomes (n=15 STs, n=244 genomes). Variant frequency (freq) in the right-hand panel indicates the total number of determinants in which each genome was identified (out of n=244 non-*K. pneumoniae* genomes). ‘Predicted R’ indicates the proportion of genomes predicted as resistant based on genotype profile, using the rules-based classifier. In the top ST freq plot, yellow=*Kv*; navy=*Kqq*, and maroon=*Kqs*.

**Figure S14.**
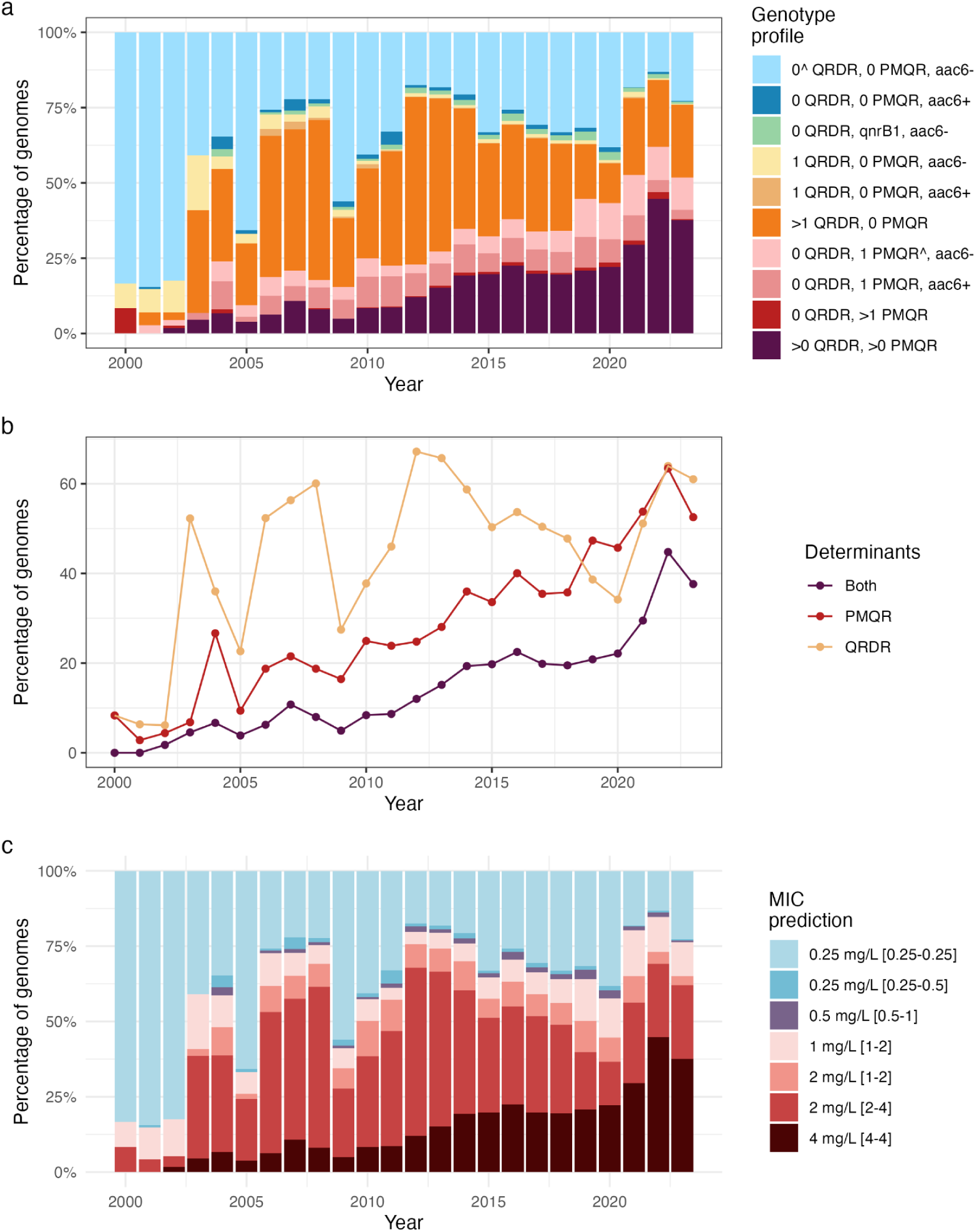
Ciprofloxacin resistance genotype profile, determinants, and MIC predictions per year. **(a)** Distribution of genotype profiles per year, inferred from 28,022 *K. pneumoniae* genomes and Kleborate-derived typing information, downloaded from Pathogenwatch. **(b)** Proportion of genomes per year carrying any quinolone resistance determining region (QRDR) mutation (yellow), any plasmid-mediated quinolone resistance (PMQR) gene (red), or both (maroon); inferred from the same data set. **(c)** Proportion of genomes and their MIC predictions per year; inferred from the same data set and the rules-based classifier from this study (square brackets indicate interquartile range). Based on EUCAST, MIC values shaded blue would be interpreted as S, those shaded purple (0.5 mg/L) as I, and pink/red/brown shades as R.

## SUPPLEMENTARY TABLES

**Table S1. Summary of KlebNET-GSP AMR Genotype-Phenotype Group discovery datasets.**

File: TableS1_discovery_dataset_summary.tsv

**Table S2. Genome assembly metrics and Kleborate output.**

Kleborate output only includes relevant columns for ciprofloxacin resistance prediction. File: TableS2_genotypes.tsv

**Table S3. Ciprofloxacin susceptibility phenotypes, metadata and public accessions.**

File: TableS3_phenotypes_metadata.tsv

**Table S4. Summary of datasets collected for validation.**

File: TableS4_validation_summary.tsv

**Table S5. External validation datasets genome assembly metrics and Kleborate output.**

Note that the data owners of the EUSCAPE & EURECA (referred to as dataset: A (EUCAST)) and IRCCS Ospedale San Raffaele datasets have independently performed external validation (i.e., no data was shared with the KlebNET-GSP AMR Genotype-Phenotype Group). Kleborate output only includes relevant columns for ciprofloxacin resistance prediction.

File: TableS5_genotypes_validation.tsv

**Table S6. External validation datasets ciprofloxacin susceptibility phenotypes and metadata.** Note that the data owners of the EUSCAPE & EURECA (referred to as dataset: A (EUCAST)) and IRCCS Ospedale San Raffaele datasets have independently performed external validation (i.e., no data was shared with the KlebNET-GSP AMR Genotype-Phenotype Group).

File: TableS6_phenotype_metadata_validation.tsv

**Table S7. Frequency of ciprofloxacin resistance determinants.**

S/I/R counts and positive predictive value (PPV) for resistance (R) and non-wild type (NWT), for each determinant found solo (i.e. in genomes with no other determinants present); in combination with *aac(6ʹ)-Ib-cr* only; overall.

File: TableS7_determinantStats.csv

**Table S8. Single-variable and multivariable associations of individual determinants with ciprofloxacin resistance.**

Regression coefficients (estimate, and 95% confidence interval) and p-values for individual determinants, alone or as an interaction term with *aac(6ʹ)-Ib-cr,* in logistic models of resistance (R) or nonwildtype (NWT). Model 1a: simple additive model including all determinants identified in ≥20 genomes; Model 1b: additive model with an interaction term for each determinant with *aac(6ʹ)-Ib-cr*.

File: TableS8_LogRegCoef.csv

**Table S9.** Logistic regression model coefficients for counts-based models.

| Prediction | Model | Determinant | est | ci.lower | ci.upper | pval |
| --- | --- | --- | --- | --- | --- | --- |
| R | 2a | (Intercept) | -3.2875322 | -3.4221909 | -3.1578852 | 0 |
|  |  | QRDR mutation count | 3.16606391 | 3.00406547 | 3.34008007 | 0 |
|  |  | acquired gene count | 4.00679912 | 3.81940358 | 4.19788437 | 0 |
|  |  | *aac(6')-Ib-cr* | 1.74526208 | 1.50847316 | 1.9848646 | 0 |
| R | 2b | (Intercept) | -3.3423694 | -3.4861258 | -3.2045937 | 0 |
|  |  | QRDR mutation count | 3.33578117 | 3.15087133 | 3.53612711 | 0 |
|  |  | *aac(6')-Ib-cr* | 2.14896219 | 1.76306212 | 2.51798909 | 0 |
|  |  | acquired gene count | 4.05737467 | 3.8500008 | 4.2691213 | 0 |
|  |  | QRDR mutation count:<br>*aac(6')-Ib-cr* | -1.2649665 | -1.6171746 | -0.8794719 | 1.69E-08 |
|  |  | acquired_gene count:<br>*aac(6')-Ib-cr* | -0.3760158 | -0.8434429 | 0.10729433 | 0.12577518 |
| NWT | 2a | (Intercept) | -2.1754551 | -2.2607275 | -2.0920161 | 0 |
|  |  | QRDR mutation count | 2.75229337 | 2.58940562 | 2.93031699 | 0 |
|  |  | acquired gene count | 4.19043288 | 3.97693009 | 4.41275439 | 0 |
|  |  | *aac(6')-Ib-cr* | 1.27006951 | 1.00615641 | 1.53390165 | 0 |
| NWT | 2b | (Intercept) | -2.2106988 | -2.298413 | -2.1249749 | 0 |
|  |  | QRDR mutation count | 2.88533173 | 2.70053261 | 3.08932239 | 0 |
|  |  | *aac(6')-Ib-cr* | 1.75889136 | 1.43525425 | 2.07632527 | 0 |
|  |  | acquired gene count | 4.37167763 | 4.12220106 | 4.63513269 | 0 |
|  |  | QRDR mutation count:<br>*aac(6')-Ib-cr* | -1.0251916 | -1.3870792 | -0.6206465 | 8.55E-06 |
|  |  | acquired_gene count:<br>*aac(6')-Ib-cr* | -0.9268128 | -1.4267755 | -0.415625 | 4.44E-04 |

File: TableS9\_CountLogRegCoef.csv

Table S10. Positive predictive values of *oqxAB* operon defects.

*oqxAB* operon defects in genomes without any other known ciprofloxacin resistance determinant. Number of genomes with or without the defect (N), susceptible (S) genomes, intermediate /ATU (I) genomes, resistant (R) genomes, non-wild type (NWT) genomes, and the positive predictive value (PPV) for R and NWT.

File: TableS10_oqxAB_contribution_PPV.tsv

**Table S11. Evaluation metrics of validation sets.**

Categorical agreement, major errors (ME), very major errors (VME), sensitivity, and specificity for predicting resistance (R) (excluding intermediate (I) phenotypes), R (including I phenotypes), and non-wildtype (NWT (i.e., R+I))) per dataset and all datasets combined (i.e., Pooled).

File: TableS11_extValidation_evalMetrics_all.tsv

**Table S12. Genotype profiles and median MIC of validation sets.**

Positive predictive values (PPV) and median MIC [interquartile range, IQR] of ciprofloxacin resistance genotype profiles (as per the rules-based classifier) stratified by dataset. Resistant PPV excluding intermediate (I) phenotype (RexcludeI_PPV), resistant PPV including I phenotype (R_PPV), and non-wildtype (NWT) PPV (i.e., R+I) are described in the table. Median MIC [IQR] per dataset are indicated in the table with I phenotype (MIC median [IQR]) and without I phenotype (MIC median [IQR] (exclude I)) for genotype profiles with more than 5 genomes/MIC values.

File: TableS12_extValidation_genotypeProfiles.tsv

## REFERENCES

1. Paczosa, M. K. & Mecsas, J. Klebsiella pneumoniae: Going on the Offense with a Strong Defense. Microbiol. Mol. Biol. Rev. 80, 629 (2016).

2. Russo, T. A. & Marr, C. M. Hypervirulent Klebsiella pneumoniae. Clin. Microbiol. Rev. 32, (2019).

3. Choby, J. E., Howard-Anderson, J. & Weiss, D. S. Hypervirulent Klebsiella pneumoniae – clinical and molecular perspectives. J. Intern. Med. 287, 283 (2019).

4. Naghavi, M. et al. Global burden of bacterial antimicrobial resistance 1990–2021: a systematic analysis with forecasts to 2050. Lancet 404, 1199–1226 (2024).

5. Institute for Health Metrics and Evaluation (IHME), University of Oxford. MICROBE. Seattle, WA: IHME, University of Washington, 2024. Available at: https://vizhub.healthdata.org/microbe. (Accessed: 6th November 2024)

6. World Health Organization. WHO Bacterial Priority Pathogens List, 2024: bacterial pathogens of public health importance to guide research, development and strategies to prevent and control antimicrobial resistance. (2024).

7. Hasso-Agopsowicz, M. et al. Identifying WHO global priority endemic pathogens for vaccine research and development (R&D) using multi-criteria decision analysis (MCDA): an objective of the Immunization Agenda 2030. eBioMedicine 0, 105424 (2024).

8. World Health Organization. The WHO AWaRe (Access, Watch, Reserve) antibiotic book. (2022).

9. FDA. FDA Access Data: CIPRO (Ref: 5449591). (2021).

10. World Health Organization. Global antimicrobial resistance and use surveillance system (GLASS) report 2022. (2022).

11. Turnidge, J. Epidemiology of quinolone resistance. Eastern hemisphere. Drugs 49 Suppl 2, 43–47 (1995).

12. Thomson, C. J. The global epidemiology of resistance to ciprofloxacin and the changing nature of antibiotic resistance: a 10 year perspective. J. Antimicrob. Chemother. 43 Suppl A, 31–40 (1999).

13. Blondeau, J. M., Yaschuk, Y., Suter, M. & Vaughan, D. In-vitro susceptibility of 1982 respiratory tract pathogens and 1921 urinary tract pathogens against 19 antimicrobial agents: a Canadian multicentre study. Canadian Antimicrobial Study Group. J. Antimicrob. Chemother. 43 Suppl A, 3–23 (1999).

14. Brisse, S., Milatovic, D., Fluit, A. C., Verhoef, J. & Schmitz, F. J. Epidemiology of quinolone resistance of *Klebsiella pneumoniae* and *Klebsiella oxytoca* in Europe. Eur. J. Clin. Microbiol. Infect. Dis. 19, 64–68 (2000).

15. Fàbrega, A., Madurga, S., Giralt, E. & Vila, J. Mechanism of action of and resistance to quinolones. Microb. Biotechnol. 2, 40–61 (2009).

16. Redgrave, L. S., Sutton, S. B., Webber, M. A. & Piddock, L. J. V. Fluoroquinolone resistance: mechanisms, impact on bacteria, and role in evolutionary success. Trends Microbiol. 22, 438–445 (2014).

17. Brisse, S. et al. Comparative In Vitro Activities of Ciprofloxacin, Clinafloxacin, Gatifloxacin, Levofloxacin, Moxifloxacin, and Trovafloxacin against *Klebsiella pneumoniae, Klebsiella oxytoca, Enterobacter cloacae*, and *Enterobacter aerogenes* Clinical Isolates with Alterations in GyrA and ParC Proteins. Antimicrob. Agents Chemother. 43, 2051 (1999).

18. Kato, J. I., Suzuki, H. & Ikeda, H. Purification and characterization of DNA topoisomerase IV in *Escherichia coli*. J. Biol. Chem. 267, 25676–25684 (1992).

19. Piddock, L. J. V. Fluoroquinolone resistance: Overuse of fluoroquinolones in human and veterinary medicine can breed resistance. BMJ Br. Med. J. 317, 1029 (1998).

20. Tsai, Y. K. et al. *Klebsiella pneumoniae* Outer Membrane Porins OmpK35 and OmpK36 Play Roles in both Antimicrobial Resistance and Virulence. Antimicrob. Agents Chemother. 55, 1485 (2011).

21. Machuca, J. et al. Impact of AAC(6′)-Ib-cr in combination with chromosomal-mediated mechanisms on clinical quinolone resistance in *Escherichia coli*. J. Antimicrob. Chemother. 71, 3066–3071 (2016).

22. Schneiders, T., Amyes, S. G. B. & Levy, S. B. Role of AcrR and RamA in fluoroquinolone resistance in clinical *Klebsiella pneumoniae* isolates from Singapore. Antimicrob. Agents Chemother. 47, 2831–2837 (2003).

23. Yamane, K. et al. New plasmid-mediated fluoroquinolone efflux pump, QepA, found in an *Escherichia coli* clinical isolate. Antimicrob. Agents Chemother. 51, 3354–3360 (2007).

24. Ruiz, J. Transferable mechanisms of quinolone resistance from 1998 onward. Clin. Microbiol. Rev. 32, (2019).

25. Jacoby, G. A., Strahilevitz, J. & Hooper, D. C. Plasmid-mediated quinolone resistance. Microbiol. Spectr. 2, (2014).

26. Rodríguez-Martínez, J. M. et al. Activity of ciprofloxacin and levofloxacin in experimental pneumonia caused by *Klebsiella pneumoniae* deficient in porins, expressing active efflux and producing QnrA1. Clin. Microbiol. Infect. 14, 691–697 (2008).

27. Wang, M., Sahm, D. F., Jacoby, G. A. & Hooper, D. C. Emerging Plasmid-Mediated Quinolone Resistance Associated with the qnr Gene in *Klebsiella pneumoniae* Clinical Isolates in the United States. Antimicrob. Agents Chemother. 48, 1295–1299 (2004).

28. Robicsek, A. et al. Fluoroquinolone-modifying enzyme: a new adaptation of a common aminoglycoside acetyltransferase. Nat. Med. 12, 83–88 (2006).

29. Ramirez, M. S. & Tolmasky, M. E. Aminoglycoside modifying enzymes. Drug Resist. Updat. 13, 151–171 (2010).

30. Shin, S. Y. et al. Characteristics of *aac(6’)-Ib-cr* gene in extended-spectrum beta-lactamase-producing *Escherichia coli* and *Klebsiella pneumoniae* isolated from Chungnam area. Korean J. Lab. Med. 29, 541–550 (2009).

31. Park, C. H., Robicsek, A., Jacoby, G. A., Sahm, D. & Hooper, D. C. Prevalence in the United States of *aac(6′)-Ib-cr* Encoding a Ciprofloxacin-Modifying Enzyme. Antimicrob. Agents Chemother. 50, 3953–3955 (2006).

32. Baker, K. S. et al. Evidence review and recommendations for the implementation of genomics for antimicrobial resistance surveillance: reports from an international expert group. The Lancet Microbe 4, e1035–e1039 (2023).

33. Djordjevic, S. P. et al. Genomic surveillance for antimicrobial resistance — a One Health perspective. Nat. Rev. Genet. 2023 252 25, 142–157 (2023).

34. Waddington, C. et al. Exploiting genomics to mitigate the public health impact of antimicrobial resistance. Genome Med. 2022 141 14, 1–14 (2022).

35. Okeke, I. N. et al. Leapfrogging laboratories: the promise and pitfalls of high-tech solutions for antimicrobial resistance surveillance in low-income settings. BMJ Glob. Heal. 5, 3622 (2020).

36. Wheeler, N. E. et al. Innovations in genomic antimicrobial resistance surveillance. The Lancet Microbe 4, e1063–e1070 (2023).

37. Sherry, N. L. et al. An ISO-certified genomics workflow for identification and surveillance of antimicrobial resistance. Nat. Commun. 2023 141 14, 1–12 (2023).

38. Kim, J. I. et al. Machine Learning for Antimicrobial Resistance Prediction: Current Practice, Limitations, and Clinical Perspective. Clin. Microbiol. Rev. 35, (2022).

39. Zhang, S. et al. Phenotypic and genotypic characterization of *Klebsiella pneumoniae* isolated from retail foods in China. Front. Microbiol. 9, 289 (2018).

40. Park, D. J., Yu, J. K., Park, K. G. & Park, Y. J. Genotypes of Ciprofloxacin-Resistant *Klebsiella pneumoniae* in Korea and Their Characteristics According to the Genetic Lineages. Microb. Drug Resist. 21, 622–630 (2015).

41. Ruppé, E. et al. From genotype to antibiotic susceptibility phenotype in the order *Enterobacterales*: a clinical perspective. Clin. Microbiol. Infect. 26, 643.e1–643.e7 (2020).

42. Stoesser, N. et al. Genome sequencing of an extended series of NDM-producing *Klebsiella pneumoniae* isolates from neonatal infections in a Nepali hospital characterizes the extent of community-versus hospital-associated transmission in an endemic setting. Antimicrob. Agents Chemother. 58, 7347–7357 (2014).

43. Gorrie, C. L. et al. Genomic dissection of *Klebsiella pneumoniae* infections in hospital patients reveals insights into an opportunistic pathogen. Nat. Commun. 2022 131 13, 1–17 (2022).

44. Harryvan, T. J., Schaftenaar, E., Franssens, B. T., Melles, D. C. & Rentenaar, R. J. EUCAST ciprofloxacin area of technical uncertainty in *Escherichia coli* and *Klebsiella pneumoniae* using BD Phoenix^TM^ and VITEK® 2 automated susceptibility systems. J. Antimicrob. Chemother. 79, 2718–2719 (2024).

45. Nguyen, M. et al. Developing an in silico minimum inhibitory concentration panel test for *Klebsiella pneumoniae*. Sci. Rep. 8, 421 (2018).

46. Thorpe, H. A. et al. A large-scale genomic snapshot of Klebsiella spp. isolates in Northern Italy reveals limited transmission between clinical and non-clinical settings. Nat. Microbiol. 2022 712 7, 2054–2067 (2022).

47. Biffignandi, G. B. et al. Optimising machine learning prediction of minimum inhibitory concentrations in *Klebsiella pneumoniae*. *Microb*. genomics 10, (2024).

48. Wick, R. R., Judd, L. M., Gorrie, C. L. & Holt, K. E. Unicycler: Resolving bacterial genome assemblies from short and long sequencing reads. PLOS Comput. Biol. 13, e1005595 (2017).

49. Hasman, H. et al. Rapid whole-genome sequencing for detection and characterization of microorganisms directly from clinical samples. J. Clin. Microbiol. 52, 139–146 (2014).

50. Lam, M. M. C. et al. A genomic surveillance framework and genotyping tool for *Klebsiella pneumoniae* and its related species complex. Nat. Commun. 2021 121 12, 1–16 (2021).

51. Diancourt, L., Passet, V., Verhoef, J., Grimont, P. A. D. & Brisse, S. Multilocus sequence typing of *Klebsiella pneumoniae* nosocomial isolates. J. Clin. Microbiol. 43, 4178–4182 (2005).

52. Inouye, M. et al. SRST2: Rapid genomic surveillance for public health and hospital microbiology labs. Genome Med. 6, 1–16 (2014).

53. Feldgarden, M. et al. AMRFinderPlus and the Reference Gene Catalog facilitate examination of the genomic links among antimicrobial resistance, stress response, and virulence. Sci. Reports 2021 111 11, 12728 (2021).

54. CLSI M100. Performance Standards for Antimicrobial Susceptibility Testing, 35th Edition. Clinical and Laboratory Standards Institute (2025).

55. EUCAST: Clinical breakpoints and dosing of antibiotics. European Committee on Antimicrobial Susceptibility Testing (2025). Available at: https://www.eucast.org/clinical_breakpoints/. (Accessed: 28th February 2025)

56. European Committee on Antimicrobial Susceptibility Testing (EUCAST). Breakpoint tables for interpretation of MICs and zone diameters*. Version 1.0. December* 2009.

57. Barry, A. L. et al. Ciprofloxacin disk susceptibility tests: Interpretive zone size standards for 5-μg disks. J. Clin. Microbiol. 21, 880–883 (1985).

58. Argimón, S. et al. Rapid Genomic Characterization and Global Surveillance of *Klebsiella* Using Pathogenwatch. Clin. Infect. Dis. 73, S325–S335 (2021).

59. Browne, A. J. et al. Global antibiotic consumption and usage in humans, 2000–18: a spatial modelling study. Lancet Planet. Heal. 5, e893–e904 (2021).

60. U.S. Department of Health and Human. Antimicrobial Susceptibility Test (AST) Systems - Class II Special Controls Guidance for Industry and FDA. (2009).

61. Pataki, B. Á. et al. Understanding and predicting ciprofloxacin minimum inhibitory concentration in *Escherichia coli* with machine learning. Sci. Reports 2020 101 10, 1–9 (2020).

62. Lipworth, S. et al. Estimating the effect of antimicrobial resistance genes on minimum inhibitory concentration in *Escherichia coli*. medRxiv 2024.05.15.24307162 (2024). doi:10.1101/2024.05.15.24307162

63. Matlock, W. et al. E. coli phylogeny drives co-amoxiclav resistance through variable expression of blaTEM-1. bioRxiv 2024.08.12.607562 (2024). doi:10.1101/2024.08.12.607562

64. Petit, R. A. & Read, T. D. Bactopia: a Flexible Pipeline for Complete Analysis of Bacterial Genomes. mSystems 5, (2020).

65. Jolley, K. A., Bray, J. E. & Maiden, M. C. J. Open-access bacterial population genomics: BIGSdb software, the PubMLST.org website and their applications. Wellcome Open Res. 3, (2018).

66. Raherison, S. et al. Expression of the *aac(6′)-Ib-cr* Gene in Class 1 Integrons. Antimicrob. Agents Chemother. 61, e02704–16 (2017).

67. Da Re, S. et al. The SOS response promotes qnrB quinolone-resistance determinant expression. EMBO Rep. 10, 929–933 (2009).

68. Jones, R. N., Erwin, M. E. & Croco, J. L. Critical appraisal of E test for the detection of fluoroquinolone resistance. J. Antimicrob. Chemother. 38, 21–25 (1996).

69. Rodriguez-Martinez, J. M. et al. Challenges to accurate susceptibility testing and interpretation of quinolone resistance in *Enterobacteriaceae*: results of a Spanish multicentre study. J. Antimicrob. Chemother. 70, 2038–2047 (2015).

70. Salam, M. A., Al-Amin, M. Y., Pawar, J. S., Akhter, N. & Lucy, I. B. Conventional methods and future trends in antimicrobial susceptibility testing. Saudi J. Biol. Sci. 30, 103582 (2023).

71. Leegaard, T. M., Justesen, U. S., Matuschek, E. & Giske, C. G. Performance of automated antimicrobial susceptibility testing for the detection of antimicrobial resistance in Gram-negative bacteria: a NordicAST study. APMIS 131, 543–551 (2023).

72. Zhou, M. et al. Comparison of five commonly used automated susceptibility testing methods for accuracy in the China Antimicrobial Resistance Surveillance System (CARSS) hospitals. Infect. Drug Resist. 11, 1347 (2018).

73. Kahlmeter, G. & Turnidge, J. Wild-type distributions of minimum inhibitory concentrations and epidemiological cut-off values—laboratory and Clinical Utility. Clinical Microbiology Reviews 36, (2023).

74. Martínez, J.L., Coque, T.M. and Baquero, F. ‘What is a resistance gene? ranking risk in resistomes’. Nature Reviews Microbiology, 13(2), pp. 116–123, (2014).

75. Chindelevitch, L. et al. Ten simple rules for the sharing of bacterial genotype-Phenotype data on antimicrobial resistance. PLoS Comput. Biol. 19, (2023).

76. World Health Organization. Catalogue of mutations in Mycobacterium tuberculosis complex and their association with drug resistance. (2023).

